# A comprehensive survey on the nature of ring:ring nucleobase stacking interactions in RNA: occurrence, structural variability and classification of the associated contacts

**DOI:** 10.1101/2020.02.28.970798

**Authors:** Ayush Jhunjhunwala, Zakir Ali, Sohini Bhattacharya, Antarip Halder, Abhijit Mitra, Purshotam Sharma

## Abstract

The astonishing diversity in folding patterns of RNA 3D structures is crafted by myriads of noncovalent contacts, of which base pairing and stacking are the most prominent. Although the classification scheme proposed by Leontis and Westhof (*RNA* (2001), 7, 499) has been widely accepted for the annotation of RNA base pairs, the absence of an unambiguous classification system for base stacks appears to be a roadblock for exploring the stacking diversity in RNA. Here we provide an unambiguous and structurally-intuitive scheme for a geometry cum topology based classification of base stacking, where a stack is essentially classified in terms of the topology of the interacting nucleobase faces and the geometry described by the relative orientation of the glycosidic bonds. For heterodimeric stacks, this generates eight basic stacking geometric families, whereas for homodimeric stacks, this generates six of those. Further annotation in terms of the identity of the bases and the region of involvement of purines (5-membered, 6-membered or both rings), leads to the enumeration of 384 distinct RNA base stacks. Based on our classification scheme, we also present an algorithm for automated identification of stacks in RNA crystal structures. Overall, the work described here is expected to greatly facilitate structure-based RNA research.

## INTRODUCTION

The recent exponential growth in the availability of atomic resolution structures of RNA molecules has not only highlighted the astonishing structural diversity of RNA, it has also created the basis for molecular level investigations into their functional diversity, which includes their regulatory and catalytic roles. The associated challenge of dealing with the complexity of RNA 3D structures, has spurred major overhauling of the inherited methodological apparatus developed earlier in the context of investigating the structural details of double stranded DNA.

The apparently complex RNA structures can be best understood in terms of a hierarchy, by considering the 3D structures as a collection of covalently-linked 2D stem-loop motifs interconnected through a network of noncovalent interactions between non-consecutive ribonucleotides in the form of stacking and base pairing (Fig. 1). Although stacking interactions involve *nonspecific* face-to-face van der Waals contacts between nucleobases, base pairing consists of *specific* edge-to-edge hydrogen bonding between two nucleotides. Since both these attractive interactions give rise to diverse geometries, in terms of accessible nucleobase orientations, the development of a reliable method for automated identification and annotation of base pairs and base stacks is essential for investigating the structural diversity of RNA.

**FIGURE 1.**
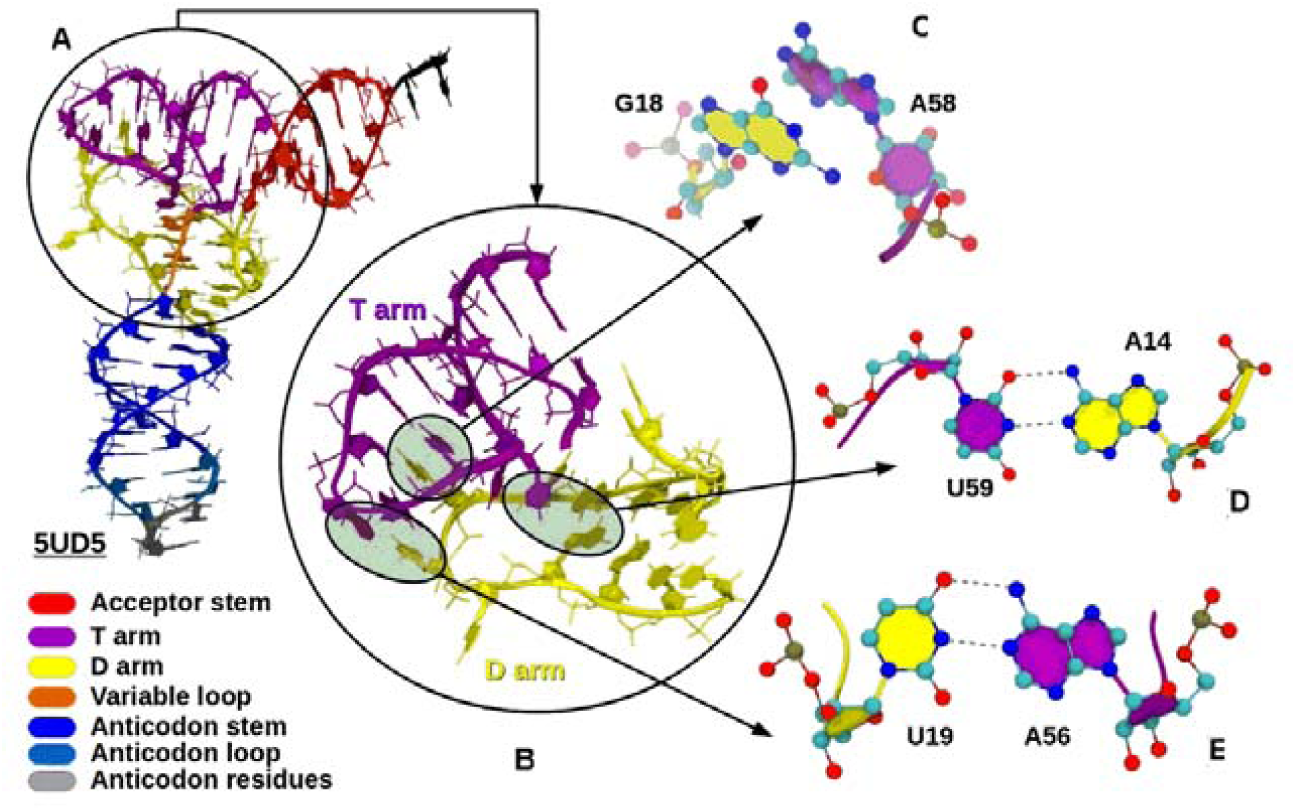
(A) Representation of a tRNA structure (PDB code: 5UD5) illustrating the noncovalent interactions between (B) T arm and D arm of tRNA. (C and D) Examples of Base pairing (C) and stacking interactions ((D) between nucleobases of D arm and T arm.

Though the two complementary strands of DNA double helices are held together mainly by canonical A:T (A:U in RNA) and G:C base pairs, it has long since been recognised that there are hot spots involving noncanonical base pairs in DNA that involve different nucleobase edges and different pairs of nucleobases. Accordingly, there have been several attempts, most notably by Saenger (Saenger 1984), at annotating the geometries of these noncanonical base pairs. Saenger’s annotation or nomenclature scheme, however, had problems in dealing with RNA base pairs which involve a significantly larger variety of noncanonical base pairs (Leontis and Westhof 2001). In this context, shortly after the publication of the high-resolution crystal structures of ribosome at the turn of century (Ban et al. 2000, Cate et al. 1999; Nissen et al. 2000; Schluenzen et al. 2000; Wimberly et al. 2000), Leontis and Westhof proposed a comprehensive geometrical classification scheme for the naming of all possible RNA base pairs (Leontis and Westhof 2001). This scheme characterizes RNA base pairs in terms of three interaction edges (Watson-Crick (W) edge, Hoogsteen (H) edge, and Sugar (S) edge) and the mutual glycosidic orientation (*cis* or *trans*) of the interacting nucleotides, and classifies RNA base-pairs into 12 distinct geometric families (i.e. W:W, W:H, W:S, H:H, H:S or S:S families in *cis* or *trans* orientations). This scheme facilitated the analysis of intrinsic structural characteristics and dynamical stabilities of different types of RNA base-pairs (see, for example, refs. Chawla et al. 2015; Halder and Bhattacharyya 2013; Havrila et al. 2013; Mládek et al. 2009; Seelam et al. 2017), and thus also provided a framework for understanding the structural implications of sequence-based homology between RNA molecules, in terms of geometries and stabilities of base pairs involved. Subsequent studies, while extending this scheme to classify and annotate higher order structures such as base triplets and quartets in RNA (Bhattacharya et al. 2019; Abu Almakarem et al. 2011), highlighted the need for supplementing the geometric framework with topological considerations.

The task of annotation of attractive interactions stabilizing RNA 3D structures is however not complete unless the intricate variations in base stacking are appropriately captured. Towards this end, there are at least four nucleic-acid annotation and visualization tools that identify stacking interactions. These include MC-Annotate (Gendron et al. 2001), ClaRNA (Waleń et al. 2014), 3DNA (Lu et al. 2008) and FR3D (Find RNA 3D, Sarver et al. 2008). Among these, MC-Annotate and ClaRNA use the stacking algorithm previously developed by Gabb *et al*. (Gabb et al. 1996), which considers two bases to be stacked if the distance between their ring centres is less than 5.5 Å, the angle between the normals to their base planes is less than 30°, and the angle between the normal of one base plane and the vector connecting the centre of the two rings is less than 40°. In addition to detecting stacking, ClaRNA introduces an additional parameter to measure the overlap area of bases (Waleń et al. 2014), albeit without specifying the identity of the overlapping rings (5- or 6-membered). On the other hand, 3DNA program developed by Lu and co-workers classifies stacking interaction from planar projection of each nucleobase ring, inclusive of their respective exocyclic atoms, in consecutive bases or base pairs (Lu et al. 2008). To detect stacking interactions, this algorithm defines a middle reference frame between two potentially stacked bases or base pairs and quantifies the stacking interaction by evaluating the shared overlap areas of the extended polygons obtained as projections of the base rings along their respective exocyclic substituents. This approach is indeed useful for capturing N-H•••π type interactions involving exocyclic amino groups, in addition to the π – π stacking instances. However, the output of this program does not specify the interacting faces of the stacking nucleobases, and thus does not distinguish between the different possible stacking topologies.

In a way, the issue of the identity of interacting faces was addressed by Sarver *et al*. who classified the stacked bases, that occur in a regular helix within the FR3D database (Sarver et al. 2008), according to the orientation of the interacting faces in the context of the strand directions in the helix. Specifically, FR3D identifies the stacking interactions based on non-zero overlap of the convex hull of each base projected vertically onto the convex hull of the other. The side of nucleobase that orients towards the 3′-end in a regular helix is designated as the 3′ face, and the other face is called the 5′ face. The base stacks are accordingly classified using an alpha-numeric notation as X-Y s35/s55/s33, where X and Y represent the stacked nucleotides, the letter ‘s’ denotes the stacking interaction, and the numbers 35, 55 or 33 denote the identity of the interacting faces (3′ or 5′) of the stacked bases. Thus, this scheme distinguishes between interstrand and intrastrand stacking, and identifies stacking interactions involving exocyclic base atoms. However, because of variations in the torsion angle χ associated with the stacked bases, defining stacking faces based on the strand direction can often be nonintuitive in complex RNA structures. Further, the scheme does not consider the extent of overlap of the stacked nucleobase rings and thus does not to comprehensively annotate the inherently complex intermolecular interaction hyperspace of stacked bases characterized by multiple, closely-spaced minima that strongly depend on the relative orientations and the extent of overlap of the interacting bases (Šponer et al. 2008). Though the annotation of relative orientation of the stacked bases is a general concern for both purines and pyrimidines, the extent of overlap is specifically problematic in case of a purine, which can either stack with another purine/pyrimidine through its 5-membered ring, 6-membered ring or both rings.

To overcome these shortcomings and to facilitate a comprehensive description and annotation of RNA base stacks, here we propose a geometry cum topology based classification scheme which is both simple as well as unambiguous. We first label the two faces of the nucleobases, independent of strand directions, as α and β, (Fig. 2), using the IUPAC numbering of the nucleobase atoms (Rose et al. 1980), and topologically classify each RNA base stack in terms of the nucleobase face participating in stacking. We then, in analogy with the base pairing classification scheme (Leontis and Westhof, 2001), introduce the mutual orientation of the glycosidic bonds (*cis* or *trans*, Fig. 3) of the stacked bases as an additional geometric parameter for annotation. Further sub-classification of the stacking geometric families thus generated, by specifying the identity of the ring (5-membered, 6-membered or both) of the purine bases participating in stacking, leads to a number of distinct stacking types that depend on the identity of the stacked nucleobases (Fig. 4). Each nucleobase stack, thus identified, is further annotated as *consecutive* or *non-consecutive*, depending on the location of the stacked nucleotides in the sequence space (Fig. 5).

**FIGURE 2.**
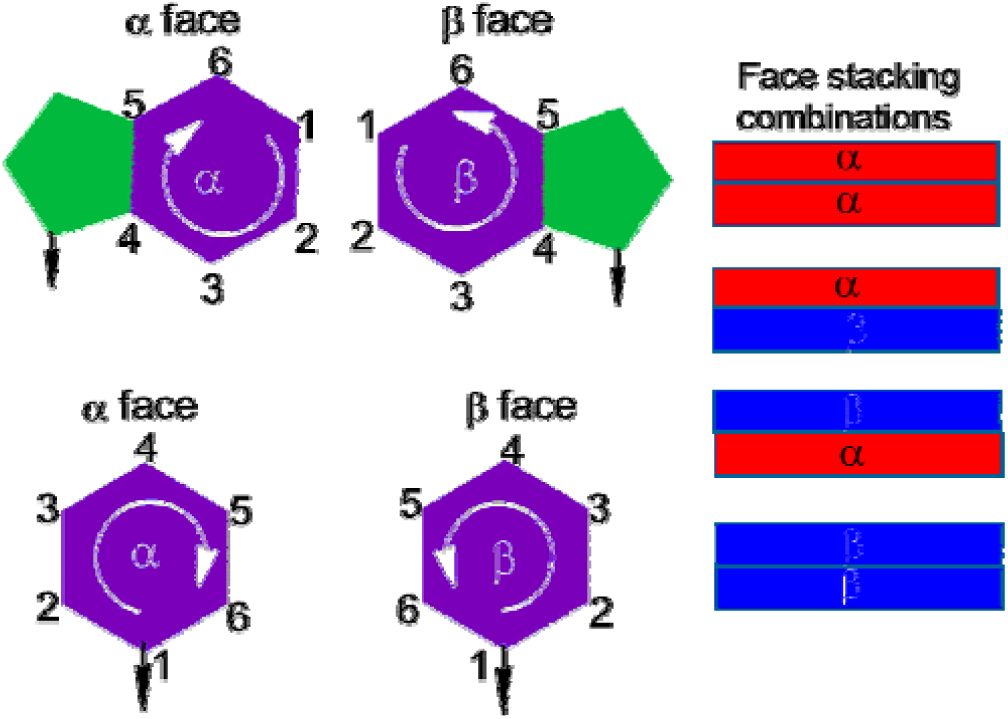
Representation of the α and β faces of purines and pyrimidines (left), and four possible face stacking combinations (right). Ring atoms are numbered according to the standard numbering nucleobases. Six membered rings and five-membered rings are represented by purple and green colors, respectively. Red colored rectangles represent the α nucleobase face and blue rectangles represent the β nucleobase face.

**FIGURE 3.**
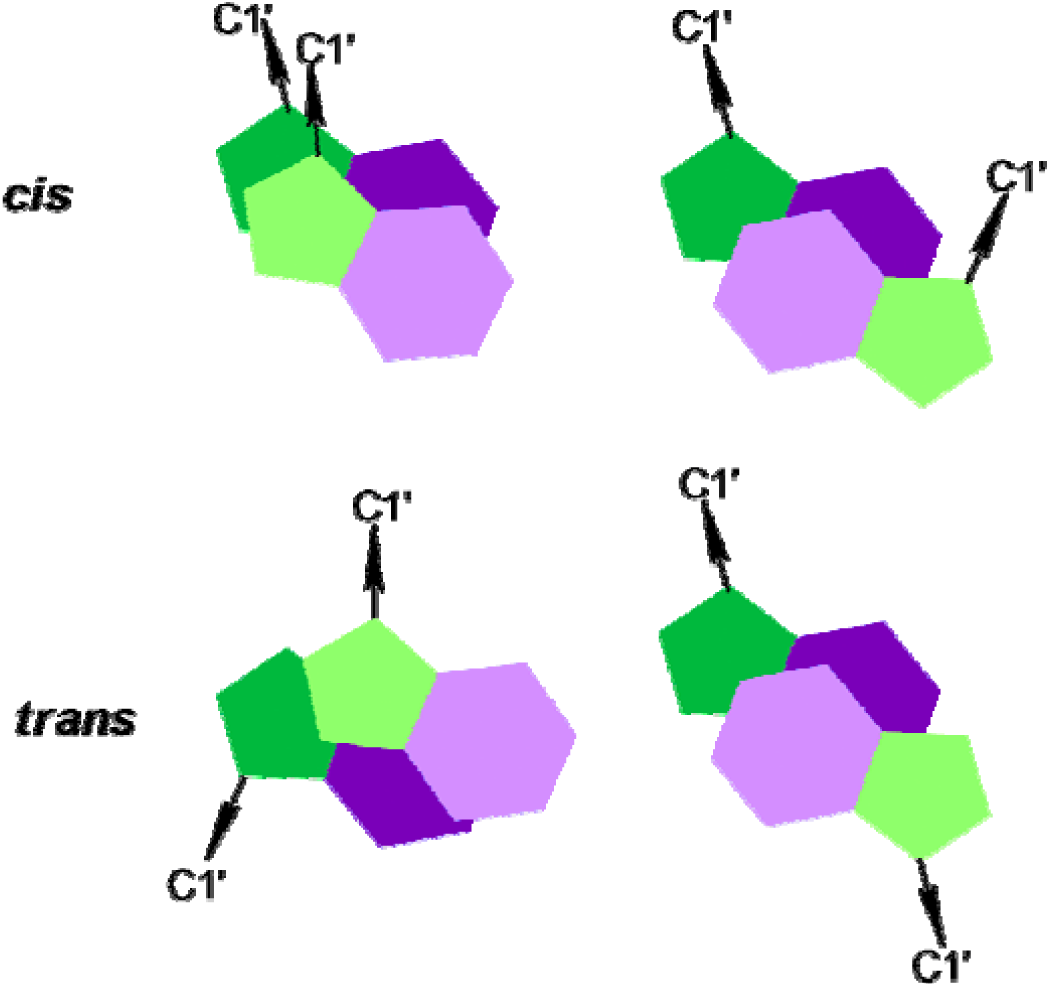
Representation of *cis* and *trans* orientation in RNA base stacks. Six membered rings and five-membered rings are represented by purple and green colors, respectively. Rings of upper base of the stack are shown in lighter in color compared to the lower base for clarity.

**FIGURE 4.**
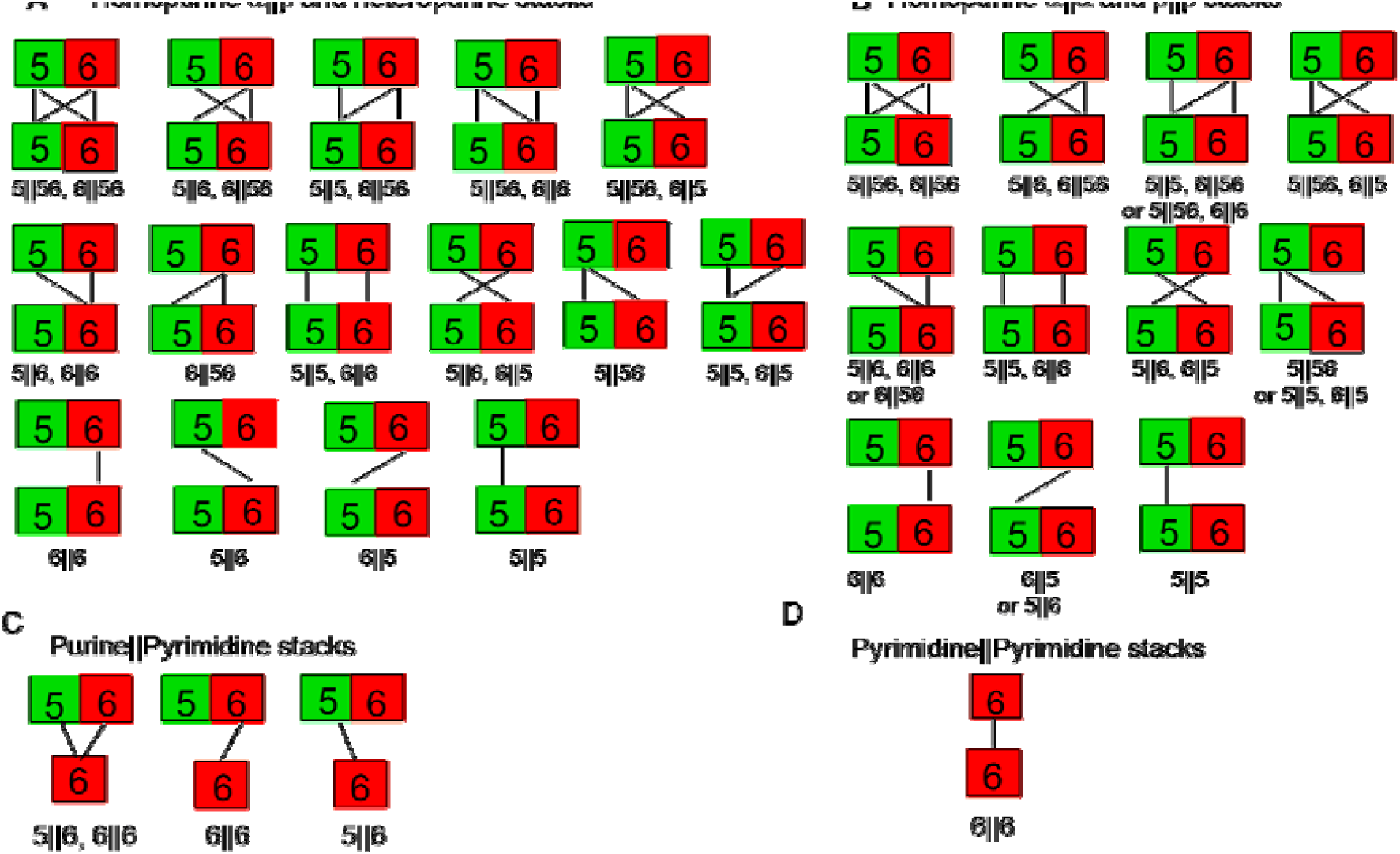
Topological variants of (A) homopurine stacks with α∥α or β∥β face orientations, (B) homopurine stacks with α∥β face orientations as well as heteropurine stacks with all possible face orientations, (C) purine∥pyrimidine stacks and (D) pyrimidine∥pyrimidine stacks.

**FIGURE 5.**
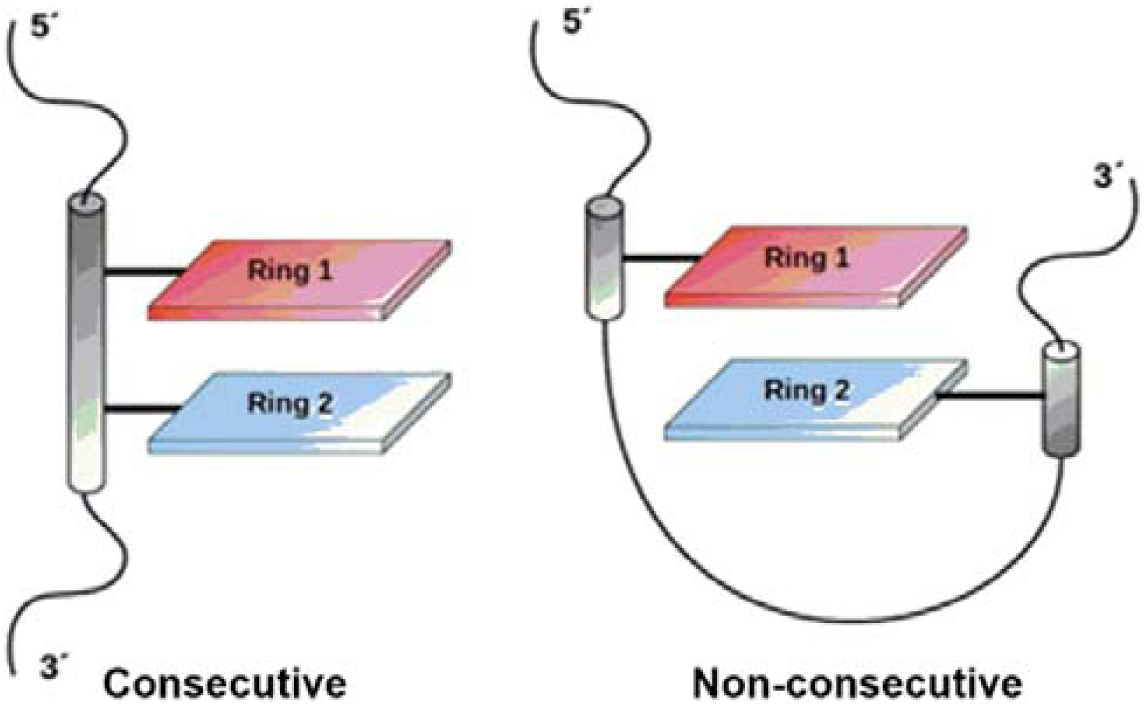
Cartoon representation of *consecutive* and *non-consecutive* stacks.

Based on this classification scheme, and by appropriately adapting from existing approaches towards detecting stacking, we have developed an algorithm for the automated identification of examples of different stacking geometries in RNA crystal structures and have classified them accordingly. In this article, we describe the classification scheme and the algorithm in detail. We also apply it to annotate base stacks present in selected RNA motifs and demonstrate the utility of our proposed classification scheme.

## RESULTS AND DISCUSSION

### Classification of RNA base stacks

As discussed in **Materials and Methods** section, our algorithm uses a slightly modified version of the stack detection protocol developed by Gabb *et. al*. (Gabb et al. 1996), to detect stacks in RNA structures in terms of the identities and residue number of the bases involved. All possible stacks, thus detected, can be classified and comprehensively annotated by recursively applying four different criteria in three stages. In the first stage, we assign the base stack to a basic stacking geometric family, based on the topology of interacting nucleobase faces and the relative orientation of respective glycosidic bonds. In the second stage, geometric variants within the basic geometric families are assigned to subclasses based on the involvement of rings (five-membered, six-membered or both) of purines in the stack. In the third stage, the stacks are annotated based on whether the stacked bases are *consecutive* or *nonconsecutive* in the sequence space. These three stages are briefly described below.

### Stage 1: Basic base stacking geometric families

#### Topology of stacking nucleobase faces

RNA nucleobases present two stacking faces α and β, which can be designated in absolute terms, i.e. without any reference to strand directions (Fig. 2). This leads to four combinations of face to face orientations (α∥α, α∥β, β∥α and β∥β, Fig. 2) within sequence stretches having heterodimeric base stacks. However, for sequence stretches with homodimeric base stacks, (i.e. for A∥A, C∥C, G∥G and U∥U), the α∥β and β∥α arrangements being identical, we get only three distinct face orientations, namely, α∥α, α∥β and β∥β.

#### Relative orientation of the glycosidic bonds

Each of the three (homodimeric) or four (heterodimeric) stacking-face combinations can be further sub-classified in terms of the (*cis* and *trans*) glycosidic bond orientations (Fig. 3). If the glycosidic bonds of the stacked nucleotides point in the same direction, the orientation is designated as *cis*, whereas the other orientation is designated as *trans*. This is geometrically determined by calculating the torsion angle σ_ab_ (∠(C1^′^)_a_–X_a_–X_b_–(C1^′^)_b_), where (C1′)_a_ and (C1′)_b_ represent the C1′ atoms of the stacked bases a and b, and X_a_ and X_b_ are the coordinates of the centroids of the respective base rings that connect to the sugar through glycosidic bond (five membered ring in case of purines and six membered ring in case of pyrimidines). The orientation of the stacked bases is annotated as *cis* if 90° ≥ σ_ab_ ≥ 0° and *trans* if 180° ≥ σ_ab_ ≥ 90°. Analysis of the distribution of σ torsion angle values reveals that while the majority of the consecutive stacked pairs exhibit *cis* orientation, the non-consecutive stacks span a wide variation in σ, with values ranging from 0° to 180° (Fig. 6). Regardless, it is quite apparent that the specification of relative orientation of the respective glycosidic bonds adds value towards an unambiguous and comprehensive annotation of stacked base pairs in RNA structures.

**FIGURE 6.**
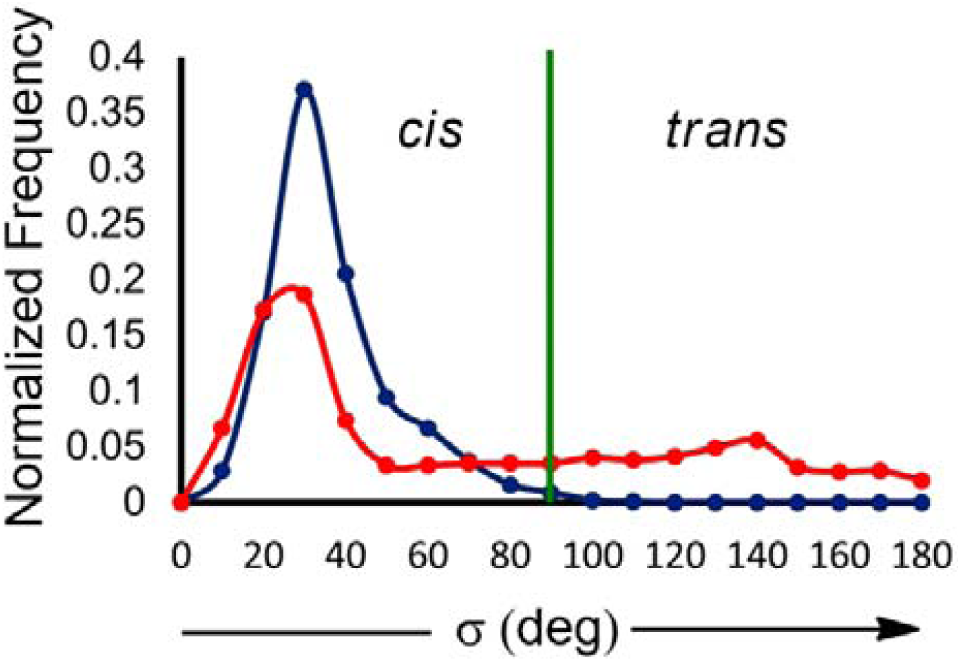
Distribution of σ torsion angle in consecutive (blue) and non-consecutive (red) stacks in the analyzed structures.

Overall, consideration of relative orientation of glycosidic bonds, in conjunction with the stacking-face combinations, lead to 8 theoretically possible stacking geometric families for heterodimer stacks and 6 such families for homodimer stacks.

### Stages 2 and 3: Geometric variant sub classification and final annotation

#### Purine∥Purine stacks

Depending on whether the five-membered, six-membered or both rings participate in stacking, a heteropurine stack (i.e. A∥G) may involve 15 distinct subclasses. (Fig. 4A). In contrast, each homopurine stack (A∥A or G∥G) may involve 11 geometric subclasses if both the purines interact with the same face (i.e. α∥α or β∥β, Fig. 4B). However, if the interacting faces are different (i.e. α∥β or β∥α), the homopurine stacks may involve 15 subclasses (Fig. 4A). The combination of 15 A∥G subclasses with four face orientations (α∥α, α∥β, β∥α and β∥β) and two glycosidic orientations (*cis* and *trans*) leads to 120 distinct stacking arrangements. In contrast, the combination of 11 subclasses each for α∥α and β∥β face combinations, and 15 subclasses for α∥β faces, with two glycosidic orientations, leads to 74 types of stacks for A∥A and G∥G respectively. Overall, this leads to (120 + 74*2) 268 distinct purine∥purine stacking arrangements.

Among the purine∥purine stacks, the three most-dominant subclasses involve the *cis* orientation (Supplemental Table S2). The most prominent subclass among these is provided by the 5∥5, 6∥56 α∥β *cis* stacking arrangement, since it maximizes the stacking overlap, and can occur between two consecutive stacked bases (Supplemental Table S2). However, 10 *cis* stacking arrangements and 10 *trans* arrangements are extremely uncommon (Supplemental Table S3), since they can occur only at uncommon σ values (close to 90°). Consequently, one example each of only two of these topologies were found in the crystal structures at σ of 83.5° and 85.3° (Supplemental Table S3). In contrast, 20 additional stacking arrangements possible at reasonable σ values were not found in the analysed structures (Supplemental Table S4). These arrangements can, however, possibly occur in RNA crystal structures available in future. Overall, examples of 230 out of 268 purine∥purine stacking arrangements were found in the dataset (Supplemental Tables S5 – S124).

#### Purine∥Pyrimidine and Pyrimidine∥Pyrimidine stacks

Since pyrimidines contain only one (i.e. six-membered) ring, the stacking of pyrimidines with five-membered and/or six-membered ring of purines leads to 3 distinct topologies (Fig. 4C). The combination of these 3 topologies with 4 face orientations (Fig. 2) and 2 glycosidic orientations (Fig. 3) leads to 24 each of A∥C, A∥U, C∥G and G∥U stacks. Overall, this leads to 96 distinct purine∥pyrimidine stacking arrangements. Of these, the arrangement involving the combination of the maximum-overlap topology (5∥6, 6∥6) and α∥α cis orientation is the most dominant, and occurs in more than a third of purine∥pyrimidine stacks detected (Supplemental Table S125). However, unlike purine∥purine stacks, examples of all 96 possible stacking arrangements were encountered in the analysed dataset (Supplemental Tables S126 – S149).

In contrast, pyrimidine∥pyrimidine stacks involve only one topology (Fig. 4D). This, in conjunction with 4 face orientations and 2 glycosidic orientations leads to 8 distinct C∥U stacking geometries. Similarly, the combination of 3 face orientations and 2 glycosidic orientations leads to 6 C∥C and 6 U∥U stacks. This leads to 20 distinct pyrimidine∥pyrimidine stacking interactions. Among these, the α∥β (or β∥α) *cis* is the most-prominent stacking arrangement, which occurs in majority (∼93%) of the cases (Supplemental Table S150). As in the case of purine∥pyrimidine stacks, examples of all 20 stacking arrangements were also found in the RNA structures investigated (Supplemental Tables S151 – S158).

Overall 268 purine∥purine, 96 purine∥pyrimidine and 20 pyrimidine∥pyrimidine stacks give rise to 384 distinct topological variants of RNA base stacks.

#### Identification of base stacks in RNA crystal structures

We further developed an automated method for classification of the base stacks present in RNA structures. Specifically, to detect stacking interactions, PDB files of each RNA structure was parsed to extract the Cartesian coordinates of the atoms that constitute a purine five-membered ring (i.e. C4, C5, N7, C8 and N9) atoms and a purine or pyrimidine six-membered ring (N1, C2, N3, C4, C5 and C6), as well as the C1′ of the attached sugar. Subsequently, the distance vector 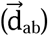 connecting the centroids of the two (potentially) stacked rings a and b, was calculated (Fig. 7). Further, a mean plane was defined for each ring (Cremer and Pople, 1975), and the vector normal to the mean plane of each ring (i.e. 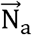or 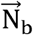) was calculated (Fig. 7). In addition, as previously suggested (Gabb et al., 1996), the relative tilt of the stacked rings (i.e. θ_ab_) was calculated as the angle between the normals to the mean planes of the two rings (Fig. 7). Finally, the angle between 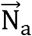 or 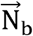 and 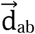 (i.e. τ_a_ or τ_b_) was calculated (Fig. 7), which in conjunction with 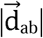, determines the horizontal shift of one monomer with respect to the other. For locating all the stacked pairs in RNA crystal structures, the following previously established criteria (Gabb et al., 1996), was used: 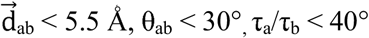.

**FIGURE 7.**
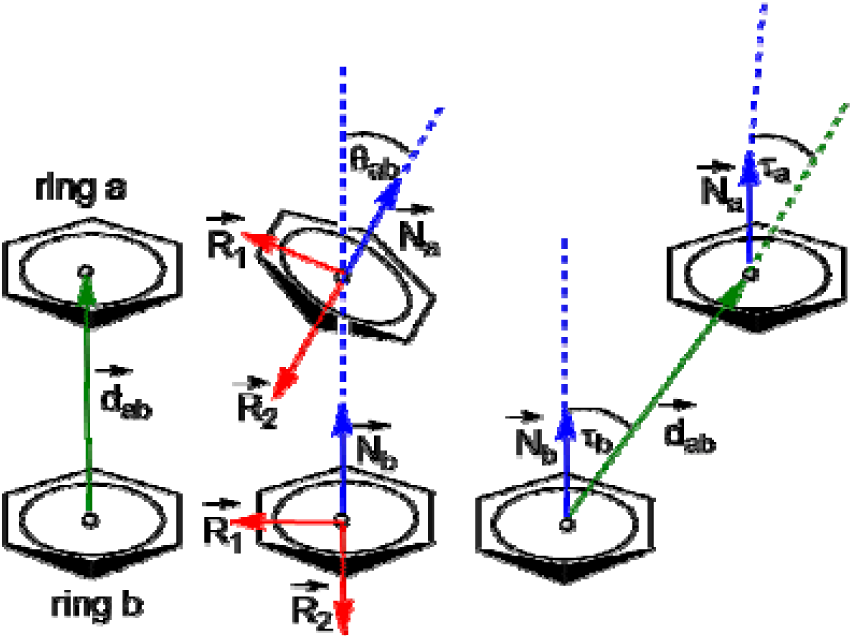
Representation of geometrical parameters (_ab_, θ_ab_, τ_a_ and τ_b_) used to locate base∥base stacking in the crystal structures of RNA.

In purine containing base stacks, topology was assigned on the basis of identity of the stacked purine ring (5-membered or 6-membered or both) (Fig. 4). Subsequently, the identity of the interacting face of the nucleobase (i.e. α or β, Fig. 2) were defined (Rose et al. 1980). Specifically, the interacting purine face is designated as α if the standard atom numbering of the 6-membered purine ring increases in the clockwise direction when viewed from the space between the stacked bases, and the other face is called β (Fig. 2). Similar protocol is followed while assigning faces to pyrimidines (Fig. 2). Accordingly, the faces were designated computationally, using the direction of normal to the mean plane vectors defined for each stacked ring. Since the mean plane vectors are calculated from atom positions around the ring (N1 to N6), the direction of the normal vectors (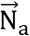 or 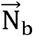) is such that it always extends from the β face. Thus, if the angle between the distance vector 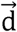 and 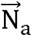 or 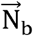 is greater than 90° the face is α, else it is assigned as β. Subsequently, the orientation of the stacked bases was annotated, depending on the value of σ angle (*cis* if 90° ≥ σ ≥ 0° and *trans* if 180° ≥ σ ≥ 90°, Fig. 3). Finally, each of the identified nucleotide was annotated as *consecutive* or *distant*, depending on the absolute value of the difference between the nucleotide residue numbers in the PDB file. Here, if the absolute value of the difference between the nucleotide numbers equals 1, then the bases are annotated as *consecutive* stacks, else they are annotated as *non-consecutive* stacks (Fig. 5).

The proposed classification has been executed on the dataset of structures downloaded from PDB (see Materials and Methods section). Further, this automated method for classification is made accessible through the web server (http://stackdetect.iiit.ac.in), where a user can generate an output file containing information on each base stack present in a given RNA PDB file, along with the average values of ring:ring stacking parameters (i.e. 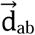, θ_ab_, τ_a_, τ_b_ and σ_ab_) for each topology. For example, for a 5∥56 stack involving two distinct ring:ring stacking contacts (i.e. 5∥5 and 5∥6), the output will display the average of the stacking parameters determined for each of the two contacts.

The consolidated output files that characterize each stack identified in PDB dataset in terms of its type and the geometrical parameters are provided on the web portal (http://stackdetect.iiit.ac.in/tnd/).

#### Examples of base stacking geometries in RNA crystal structures

Fig. 8 explains the protocol for identification of interacting faces of a base stack using the U513∥C512 α∥β *cis* stack found in glutamyl-tRNA (PDB: 1g59, (Sekine et al. 2001)). When viewed from the direction of the interacting face in the stack, the numbering of atoms in C512 increments in the counterclockwise direction. Hence, the interacting face is designated as β (Fig. 8). On the other hand, when similarly viewed, the numbering of atoms in U513 increments in clockwise progression (Fig. 8) and the face is assigned as α. This, in combination with the *cis* glycosidic orientation (Fig. 8) leads to annotation of this particular stack as U513∥C512 α∥β *cis*.

**FIGURE 8.**
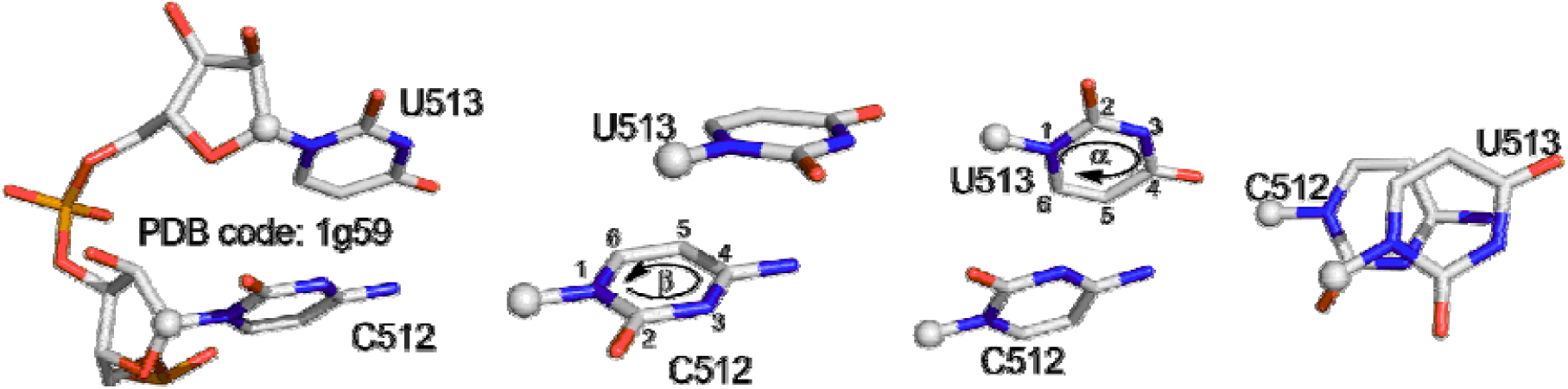
Illustration of the assignment of α and β face orientations with the help of a U∥U α∥β *cis* stack.

Although similar naming convention can be used in case of purine∥purine and purine∥pyrimidine stacks, additional information regarding the stacking topology also needs to be specified in these cases. For example, the designation of A2841∥G2842 α∥α 5∥56 *cis* stack found in PDB 1kc8, indicates that the stacking of A and G occur through their respective α faces, where the five membered ring of A stacks with both the five membered and six membered rings of G in a *cis* glycosidic orientation. Similarly, G1163∥U1164 α∥β 5∥6, 6∥6 *trans* stack found in PDB 3ccu, implies the stacking of α face of G with the β face of U in a *trans* glycosidic orientation, where both five- and six-membered rings of G stack with the six-membered ring of U.

Each of the 384 theoretically possible stacking arrangements can occur between nucleotides which are either in tandem position in sequence space or are separated by other nucleotides within the sequence space. For example, Fig. 9 shows the PDB examples of six possible stacking arrangements of a U∥U stack. Although the first three arrangements illustrate the stacking of two Us that are *consecutive* in sequence space, the remaining three arrangements are taken from structures where the stacked Us are *non-consecutive* in sequence space. Examples of *consecutive* and *non-consecutive* stacking corresponding to all base stacks identified from analysis of the complete RNA dataset are provided in the Supplementary Information (Supplemental Tables S5 – S124, S126 – S149 and S151 – S158). Further, a comparison of nomenclature of stacks identified by our method with the other available methods is provided in Supplemental Table S159.

**FIGURE 9.**
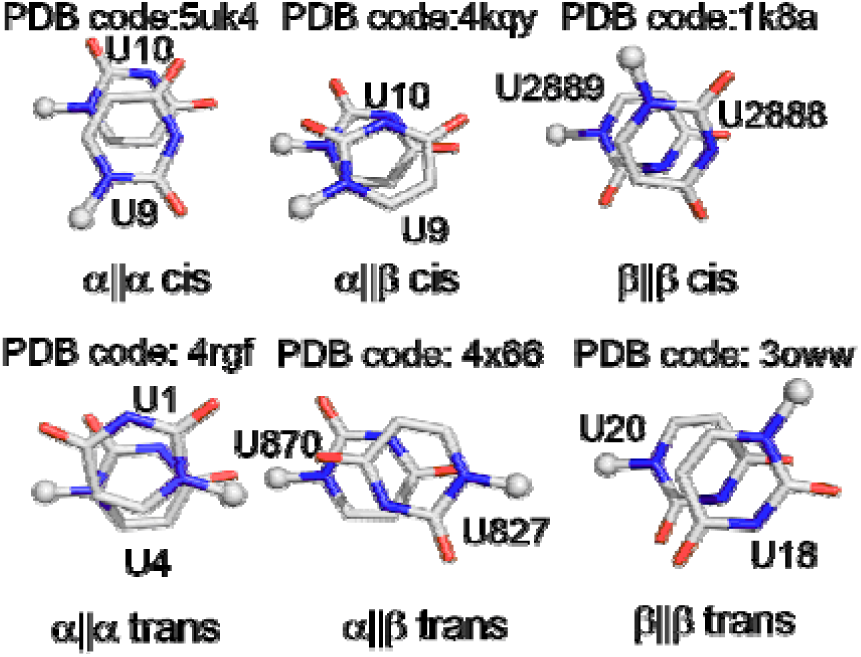
Six possible stacking arrangements of U∥U stacks. Top three arrangements are shown using *consecutive* stacks, whereas bottom three arrangements illustrate non-consecutive stacks occurring in RNA crystal structures.

### Annotation of stacking contacts in RNA 2D representations

As previously suggested for base pairs (Leontis and Westhof 2001), it is desirable to annotate the stacking arrangements in a RNA molecule on a standard 2D secondary structure representation. This would help to visually recognize the essential features of stacking interactions in an RNA motif. In this context, we propose that to designate base stacking in general, one could use the symbol “∥”. Further, we suggest filled or hollow arrow symbol to unambiguously specify the stacking face. Thus, when the head of the arrow symbol interacts with the other arrow, it represents the α interacting face of the base. Alternatively, if the tail of the arrow symbol interacts with the other arrow, it represents the β interacting face. Further, two filled arrows denote *cis*, whereas hollow arrows represent *trans*, glycosidic stacking orientation. For example, the symbol 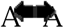 and 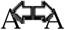 would represent β∥β *cis* and β∥β *trans* arrangement respectively. Symbols for all the variants of distinct mutual stacking arrangements can be seen in Table 1.

**Table 1.**
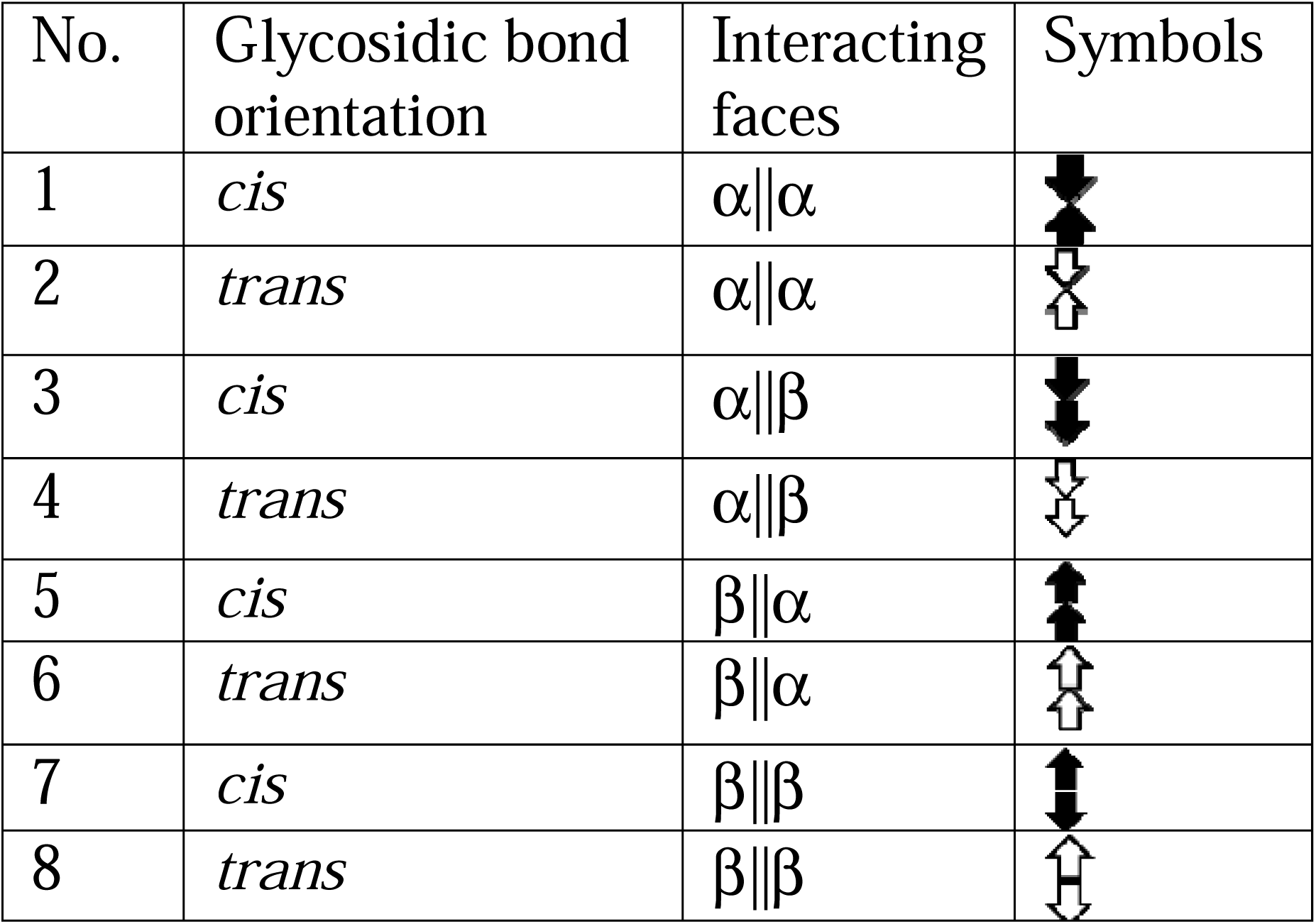
Symbols for representations of eight stacking geometrical families.

### Stacking contacts in representative 2D RNA structures

To further illustrate the symbol convention for denoting stacking variants, we present examples of 2D representations of three well-known RNA motifs used previously to represent distinct base pairing geometries (Leontis and Westhof 2001). These include the loop E of bacterial 5S rRNA (PDB code: 364d (Correll et al. 1997)), the sarcin/ricin motif from the large bacterial ribosomal subunit (PDB code: 430d, (Correll et al. 1998)) and the Domain IV of Signal Recognition Particle (SRP) 4.5 S RNA (PDB codes: 1dul and 1duh).

#### Loop E of bacterial 5S rRNA

Panel A in Fig. 10 represents the secondary structure of the loop E of bacterial 5S rRNA (Correll et al. 1997). The highly conserved loop E is the binding site for the ribosomal protein L24 in the *E. coli* ribosome (Leontis and Westhof 1998). All bases of this symmetric “internal loop” are paired. Further, all the base pairs are stacked on top of each other to form a chain like structure (Fig. 10A). Although base pairing interactions within the motif are well known and have been previously annotated by Leontis and Westhof (see green symbols in Fig. 10A, (Leontis and Westhof 2001)), here we provide complete description of complex stacking interactions within the motif using our classification scheme.

**FIGURE 10.**
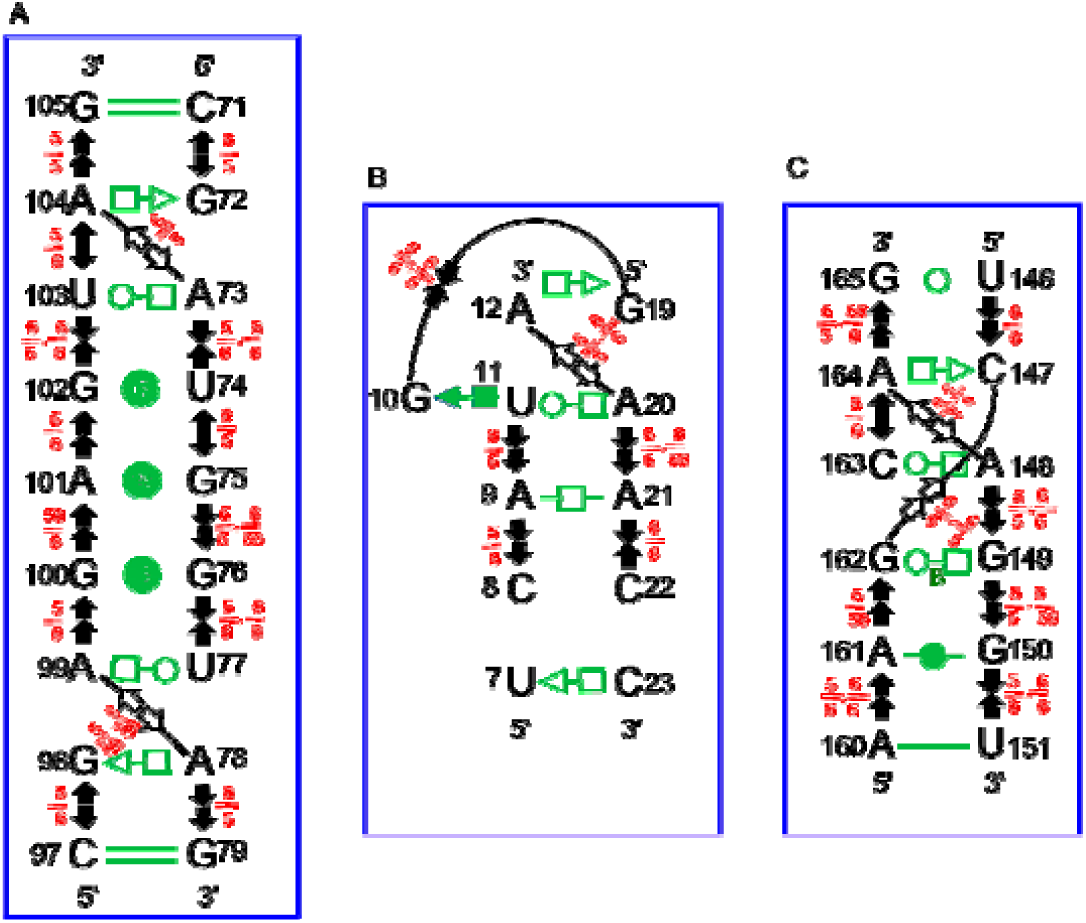
Annotation of stacking interactions using arrow representations in the (A) loop E motif of bacterial 5S rRNA (PDB code: 364D), (B) sarcin/ricin motif of the large ribosomal subunit (PDB code: 430D) and (C) domain IV of SRP 4.5 S rRNA (PDB code: 1DUL and 1DUH). Green symbols represent base pairing interactions previously described by Leontis and Westhof (Leontis and Westhof 2001).

As shown in Fig. 10A, the A104∥G105 *consecutive* stack, which involves a α∥β 5∥5 *cis* interaction, is designated using filled arrows (indicating the *cis* geometry), with the arrow placed above A104 has its head (representing its α face) pointing towards G105, and the other arrow placed below G105 with its tail (representing its β face) facing A104 (Fig. 10A). However, A104∥A73 6∥5 and A99∥A78 5∥56, 6∥56 are β∥β *trans* cross-strand stacking interactions, which are indicated by hollow arrow symbols sharing their tails respectively. Fig. 10A also shows how the numerical topological descriptors can indicate that different set of rings are involved in stacking interactions.

On similar lines, remaining base stacks are annotated in Fig. 10A. However, although FR3D algorithm detects an interstrand N-H•••π contact between G75 and G112, such a contact is not identified by our algorithm, since we focus on ring stacking in this work (Supplemental Table S160). Nevertheless, the classification of such contacts can easily be done along similar lines. Further, considering only base-pairing interactions, Leontis and Westhof observed that the bacterial loop E motif comprises of two isosteric submotifs oriented in opposite (palindromic) directions (Leontis and Westhof 2001). While this may apparently imply the presence of a C2 symmetry, this is clearly ruled out when stacking interactions within the motif are taken into consideration. Thus, these relationships can easily be depicted, in RNA motifs, with the help of the symbols we have proposed.

#### Sarcin/ricin motif of the large ribosomal subunit

The highly conserved sarcin/ricin motif (Leontis and Westhof 1998) in Domain E of eukaryotic 5S rRNA structure comprises of a GAGA hairpin loop (not shown in figure) and an asymmetric internal loop (Fig. 10B). Here the base G10 lies in the same plane as the base pair U11∥A20 and thus represent a “bulge”. Although the base pairing geometries of the five base pairs (i.e. A12:G19, G10:U11, U11:A20, A9:A21 and U7:C33) were previously described by Leontis and Westhof (see green symbols in Fig. 10 (Leontis and Westhof 2001)), analysis of the intra-motif stacking relationships between bases reveals six distinct base∥base stacks. Two of these are, the A12∥A20 β∥β 56∥5 *trans* cross-strand stacking interaction, and G19 (base pairing with A12) stacked upon G10 in α∥α 56∥5 *cis* geometry. Further, the stacking interaction between U11 and A9 reported earlier by Leontis and Westhof (Leontis and Westhof 2001) is topologically annotated as α∥β 6∥5 according to our classification scheme. In addition, A9∥C8, A20∥A21 and A21∥C22 stacks may be annotated, respectively, as α∥β 5∥6 *cis*, α∥β 5∥5,6∥56 *cis* and α∥α 6∥6 *cis*.

#### Domain IV of SRP 4.5 S rRNA

This symmetric internal loop motif is very similar to the submotifs of the bacterial loop E motif (Figs. 10A and C). Our analysis reveals that this motif comprises of a G165∥A164 β∥α 6∥6,5∥56 *cis* stacking along with A164∥C163 β∥β 5∥6 *cis* stacking and a A164∥A148 β∥β 6∥56 *trans* cross-strand stacking interaction (Fig. 10C). These interactions correspond to G105∥A104, A104∥U103 and A104∥A73 stacking interactions respectively in the E submotif, albeit with different set of rings stacked (Fig. 10A). Interestingly, in SRP, C147 and G162 show cross-strand stacking interaction in β∥α 5∥56 *trans* geometry, which does not occur in the loop E motif (Fig. 10A).

### Occurrence of selected stacking topologies in 3D RNA structures

To illustrate how the proposed classification scheme can enhance our understanding of stacking interactions in RNA 3D structures, we annotate the stacking interactions in certain specific RNA motifs. First, we annotate the stacking interactions around the G:C linchpin pair (i.e. G2:C74 pair (blue) in (Supplemental Fig. S2) in the tRNA-like structure from the 3′-end of the turnip yellow mosaic virus (PDB id: 4p5j (Colussi et al. 2014)). This pair has been previously compared with the U64:A85 base pair, observed in SAM-I riboswitch from *Thermoanaerobacter tengcongensis* (PDB:2gis (Montange et al. 2006)), and similarities in the base-stacking interactions around the two pairs have been noted (Lu et al. 2015). However, detailed comparison of the two structures reveals geometrical differences in the stacking patterns around the G:C linchpin pair and the U:A pair (Supplemental Fig. S2 (Lu et al. 2015)).

We further compare the stacking interactions in three specific diloop motifs-the CUAG (C55 to G58) loop of the CRISPR Cas9-sgRNA-DNA ternary complex (chains B and C, PDB id: 4oo8 (Nishimasu et al. 2014)), the UUGA diloop (Lu et al. 2015), which is recognized by the sequence specific RNA binding sterile alpha motif domain of yeast post *trans*criptional regulator Vts1p (PDB id: 2f8k (Aviv et al. 2006)) and the CUUG diloop (Lu et al. 2015), which was previously characterized as a tetraloop (PDB code: 1rng (Jucker and Pardi 1995)). A previous comparison had revealed that although all three loops belong to the ‘diloop’ category, UUGA and CUAG possess similar structural features that markedly differ from those of CUUG (Lu et al. 2015). This is further substantiated by our analysis, which reveals significantly smaller stacking overlap in CUUG loop compared to the other two loops (Supplemental Fig. S3). In addition, our analysis highlights subtle differences in the extent of stacking overlap within UUGA and CUUG loops. Overall, both these examples illustrate how annotation of stacking contacts in RNA 3D structures can aid in visual understanding of the subtle structural differences in apparently similar motifs.

## CONCLUSIONS

In the present work, we have proposed a classification scheme for annotation and characterization of base stacks in RNA structures, and, based on it, have presented an automated method for classification of base stacks. This is expected to highlight the rich diversity in RNA base stacking and enhance our understanding of the role of stacking interactions in biology. We demonstrate this by analysing specific RNA structural and functional contexts that involve the role of base stacks. In addition, we have proposed symbolic representations for easy visualization of stacking interactions in secondary structures as well as in 3D structural diagrams of RNA. Overall, the work described here is expected to greatly facilitate structure-based understanding of RNA functions.

## MATERIALS AND METHODS

A dataset containing all RNA crystal structures deposited in the Protein Data Bank (PDB) till 21^st^ March 2019 with the resolution of up to 3.5 Å was used to identify the occurrences of base stacking geometries considered in the present work (Table S1). Each PDB file was parsed to extract the Cartesian coordinates of atoms constituting the base rings, as well as the C1′ of the attached sugar. As part of exception handling, nucleotides with incomplete information (generally missing ring atoms), or cases where two nucleotides belonging to different chains possess same atomic coordinates (e.g. in PDB code: 488d), were ignored.

To calculate ring-ring stacking, two types of rings were considered-the six-membered heterocyclic ring of each canonical (purine or pyrimidine) nucleobase and the five-membered heterocyclic ring of each purine (adenine or guanine). As a first step, the coordinates of centroid of each ring (*i.e. x*_*c*_, *y*_*c*_, *z*_*c*_) were determined using the cartesian coordinates (*i.e*. (*x*_1_, *y*_1_, *z*_1_), (*x*_2_, *y*_2_, *z*_2_), (*x*_3_, *y*_3_, *z*_3_),……(*x*_*n*_, *y*_*n*_, *z*_*n*_)) of 1, 2, ….n (n = number of atoms that constitute the ring (5 or 6)) atoms that constitute the ring, using the following relations:

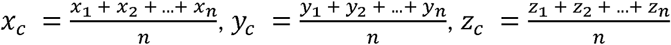

Subsequently, the position vector 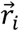 of each *i*^th^ ring atom was determined by using the cartesian coordinates (*x*_*i*_, *y*_*i*_, *z*_*i*_) of the atom *i*, and the coordinates of the ring centroid. Specifically,

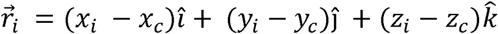

Further, the distance vector (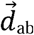, Fig. 7) that connects the centroids of two stacked rings *a* and *b* was determined using the coordinates of their centroids (*x*_*ac*_, *y*_*ac*_, *z*_*ac*_ and *x*_*bc*_, *y*_*bc*_, *z*_*bc*_ respectively).

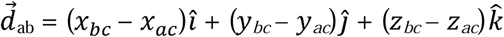

To calculate ring-ring stacking, a mean plane for each ring was defined using the two orthogonal vectors 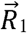 and 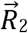 with the help of the method suggested by Cremer and Pople (Cremer and Pople 1975). Specifically,

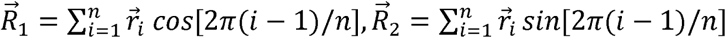

Further, 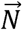, the vector normal to the mean plane vectors 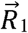 and 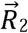 of each ring (Fig. 7), was determined as their cross product.

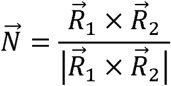

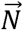 was then used to calculate *θ* _*ab*_ (Fig. 7), the angle between the normals (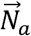 and 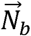) of the stacked rings *a* and *b*, as previously suggested by Gaab *et al* (Gabb et al. 1996). Specifically,

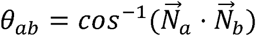

Further, the angles of 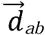 with 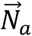 and 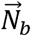 (*i.e. τ*_*a*_ and *τ*_*b*_ respectively, Fig. 7) for the stacked rings were calculated as:

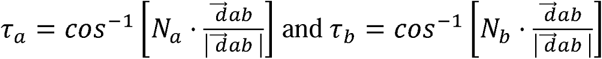

For locating all the stacked pairs in RNA crystal structures, the following criteria established in a previous study (Gabb et al. 1996), was used was used: 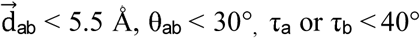.

## Notes

http://stackdetect.iiit.ac.in/

### Competing Interest Statement

The authors have declared no competing interest.

### Summary of Updates

The version as been modified for format of RNA journal. Further, a few editing changes have been done.

