## Supplementary Information for "A comprehensive survey on the nature of ring:ring nucleobase stacking interactions in RNA: occurrence, structural variability and classification of the associated contacts"

**to**

### Table of Contents

**Figure S1.** Structural representation and chemical numbering of the canonical RNA nucleobases…………..…………………………………………………………...…… S3
**Figure S2.** Annotation of stacking interactions within the base stacks around the linchpin pair of the viral tRNA mimic and the SAM-I riboswitch…………..………..………………..….S4
**Figure S3.** Annotation of stacking interactions with three different diloop motifs…...…..…S5
**Table S1.** PDB codes of 2669 RNA crystal structures studied in the present work……..…S6
**Table S2.** Percent occurrence frequency of three most dominant topologies of purine||purine stacks……………………………………………………………………………………..S10
**Table S3.** List of purine||purine stacking topologies that can occur only at extreme σ values (close to 90°)……………………………………………………………………………..S11
**Table S4.** List of purine||purine stacking topologies that are possible at reasonable σ values, but not found in RNA crystal structures…………………………………………………………………………...….S12
**Table S5-S124**. Examples of different types of purine||purine stacks identified from RNA crystal structures, along with their geometrical parameters…….……..……………..…...S13
**Table S125.** Percent occurrence frequency of three most dominant topologies of purine||pyrimidine stacks………………………………………………………..……….….S70
**Table S126-S149**. Examples of different types of purine||pyrimidine stacks identified from RNA crystal structures, along with their geometrical parameters…..………..…………..…S71
**Table S150.** Percent occurrence frequency of three most dominant topologies of pyrimidine||pyrimidine stacks……………………………………………………..…..S82
**Table S151-S158**. Examples of different types of pyrimidine||pyrimidine stacks identified from RNA crystal structures, along with their geometrical parameters……….……...….S83
**Table S159.** Comparison of the method of detection of all topologies of A||G stacks by our methods with other available methods……………………..……………………………S87
**Table S160.** Comparison of stacking interaction identified in the Loop E of bacterial 5S rRNA (PDB code: 364d) using our method and FR3D……………………...…………..S91

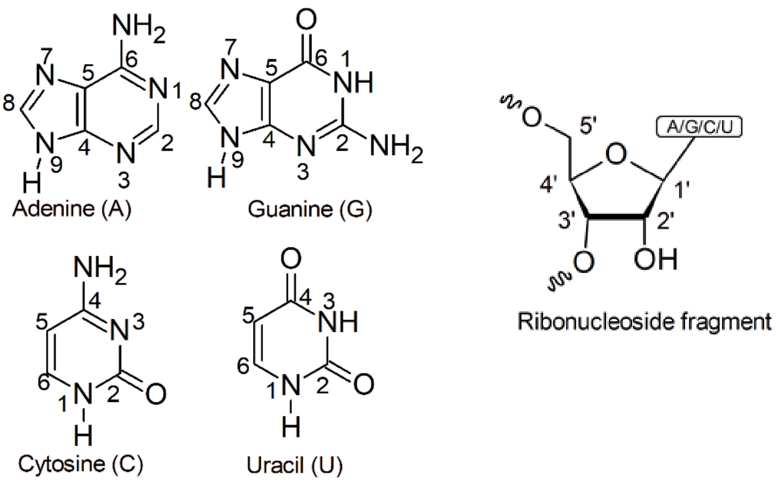

**Figure S1.** Structural representation and chemical numbering of the canonical RNA nucleobases and the ribonucleoside fragment of RNA.

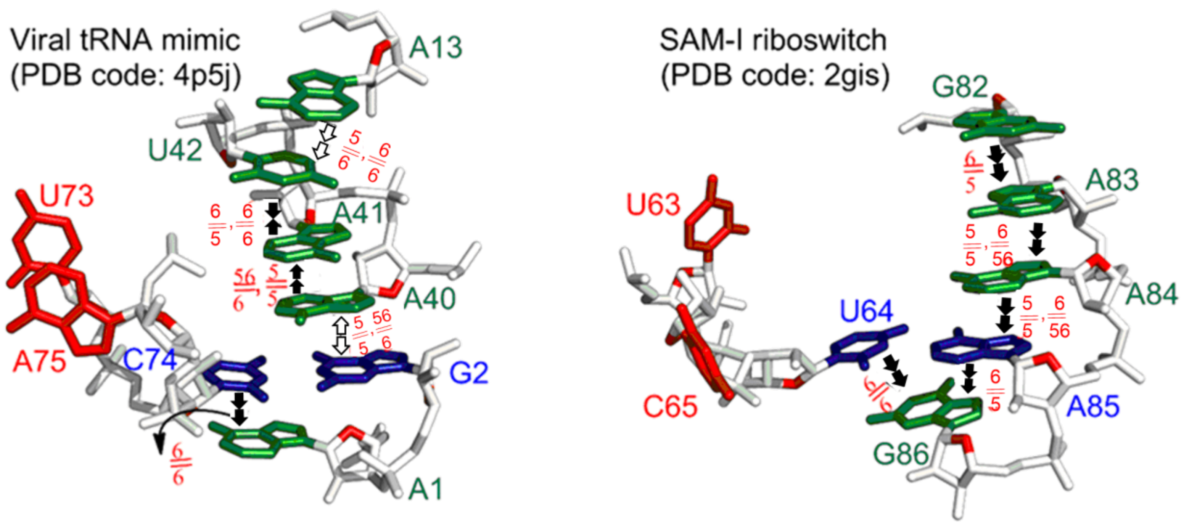

**Figure S2.** Annotation of stacking interactions within the base stacks around the linchpin pair of the viral tRNA mimic and the SAM-I riboswitch.

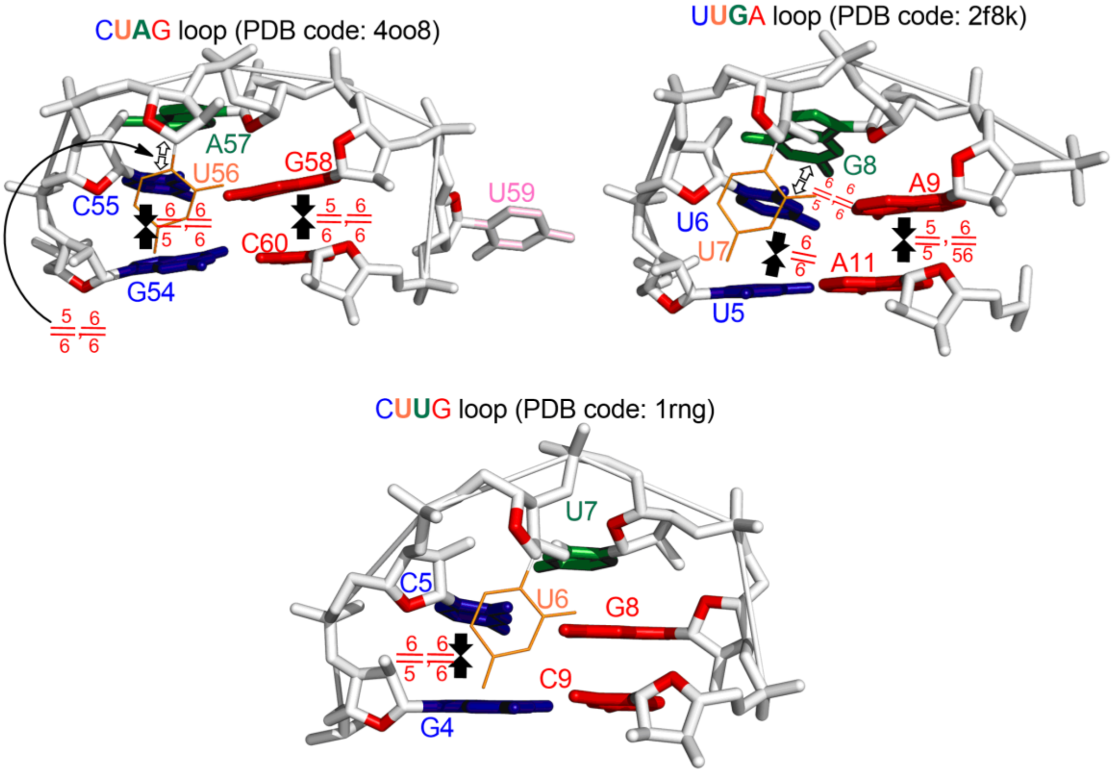

**Figure S3.** Annotation of stacking interactions with three different diloop motifs.

**Table S1.** PDB codes of 2669 RNA crystal structures studied in the present work.

4k31, 5tgm, 3rtj, 3d2s, 5ed1, 2xnr, 3bsn, 3ova, 3ccu, 3o7v, 5the, 2g5k, 1hr0, 3cun, 3bo4,
4kre, 3p22, 1uvi, 3oij, 1evv, 5fjc, 3cgs, 2jlw, 3vyx, 6noc, 4x4r, 4nfp, 6dol, 1pgl, 5wwe,
4k0k, 1nji, 2esj, 2qek, 5j8a, 6bm4, 2cky, 3ges, 2xs5, 4gxy, 1yij, 3v7e, 5fw2, 4v6c, 5b2q,
5jju, 1mms, 1tn2, 5it8, 1n35, 4ed5, 4io9, 5eev, 1s72, 6hc5, 1egk, 4v5c, 5hr7, 4c7o, 3c44,
2g32, 4wu1, 4as1, 4lf8, 5gjb, 4wqu, 3dil, 5hr6, 6c8e, 3ftm, 4hot, 3wfs, 4yvi, 6dpp, 2bcz,
5fk2, 4ngd, 6i3p, 3cgr, 3p6y, 2c51, 3o8r, 259d, 4kr3, 2f8t, 6cap, 5zsc, 4r3i, 4tuw, 4wtf,
3k5q, 2f4u, 6bjx, 1c9s, 4lvz, 4v5s, 1vy5, 4xww, 2r7x, 1r3e, 4u34, 2zm6, 3mjb, 6dpm,
5y85, 2q66, 1j8g, 5wnr, 3gpq, 1s77, 3wqy, 2ogm, 5wwf, 2qa4, 4wfa, 3skr, 2pjp, 4v5p,
5f5f, 5v6x, 1zdi, 3gs5, 6dmn, 1hc8, 4pdq, 3kna, 3pu4, 4v6f, 4wfl, 4rwp, 4u56, 4r4p, 2vrt,
2r7u, 4wqf, 4rdx, 2xb2, 466d, 4wzo, 3cw6, 1k9m, 4ji7, 4oau, 4e8q, 5lta, 5dfe, 1nta, 6gsl,
2zm5, 6dlr, 1zjw, 1ttt, 4u24, 3lrr, 2f8s, 5fdu, 1saq, 2h0s, 3ccm, 3kmq, 6c5l, 3irw, 1tn1,
1av6, 3zd4, 6fqr, 4pr6, 3eph, 5btm, 1csl, 5bym, 4oq9, 4lfb, 4dr2, 6dnr, 5ns3, 1qtq, 5e81,
5wzg, 2o44, 5uef, 5w3v, 3bbm, 1tfy, 3npq, 5zsb, 2x2q, 4y4o, 4v64, 3gvn, 6h9h, 1vqm,
4w2h, 4dwa, 2gjw, 4oqu, 6dos, 5ckk, 4krf, 4y27, 5wlh, 1kd5, 3r1e, 3hjw, 3t1y, 4lnt, 6c8d,
6bm2, 3r1h, 165d, 3gtl, 2zh3, 1vq6, 2vod, 1k73, 1mzp, 2ygh, 4zer, 5tdj, 1sa9, 1yit, 1wz2,
2zh7, 5vr4, 4p97, 6d3p, 5dox, 5nef, 4ohz, 4z0c, 2h0w, 4v8q, 5dun, 2b8s, 3s4p, 4bw0,
6doo, 1e7k, 1vqp, 4u35, 3nnc, 6c65, 5eme, 2w89, 1hys, 3q0l, 4al5, 1xmq, 2dr8, 4tux,
5hkc, 2c4y, 5de8, 3l25, 6bmd, 5hn2, 5e18, 2czj, 1h2c, 5aor, 5bz5, 3er9, 6dn3, 3ccv, 2zh9,
4x0b, 4v95, 5lr4, 1y6s, 3ibk, 4z4e, 1ooa, 2ykg, 4p5j, 4zc7, 3i2r, 1gtn, 4v9l, 5npm, 4a3k,
3q51, 3u2e, 2fcz, 4ola, 5j4d, 5w0m, 4ji1, 1q93, 4gkk, 5wt1, 1ddl, 4wzm, 4m7d, 2ozb, 2r7s,
2vqf, 5bz1, 2yif, 3szx, 4s2x, 5bws, 4ngc, 3v6y, 5ay2, 4fax, 3uld, 2f4t, 4u6f, 5wnu, 4frg,
5usg, 5zq8, 5eeu, 6cab, 4u3p, 4jya, 2ho7, 2z74, 4fen, 4wtd, 4zt9, 5dqk, 2dr5, 4rqf, 5l00,
4e59, 4m2z, 4kyy, 5elr, 5omw, 2pn4, 4x2b, 1xjr, 4wf1, 4yoe, 4v7j, 5t16, 5amr, 3hvr, 3mj0,
5ndj, 2deu, 5i9h, 1mfq, 4g7o, 1zx7, 1y39, 4duy, 4pcj, 3bo3, 1e8o, 4phy, 6c8l, 3foz, 483d,
3glp, 3s2d, 5nzd, 3skt, 5xwp, 2a64, 1y6t, 2vop, 1x9k, 5ud5, 5jvg, 5jcf, 4eya, 4a3d, 3hk2,
3l26, 5f0s, 4v67, 1c0a, 5b63, 3a6p, 4b5r, 1yty, 1r3o, 6dmv, 5ytt, 5bzu, 4woi, 6du4, 3avw,
5b2o, 5udk, 5iwa, 4u4z, 3mqk, 3cce, 3cqs, 2g8v, 6dod, 4u4u, 2dr7, 1xp7, 3hxm, 4u3o,
3iem, 2uxc, 3hoy, 2ees, 4v4f, 5jb2, 4yaz, 5lqt, 4v9o, 1wne, 5zq0, 5id6, 5bs3, 5kpy, 2xnw,
1m8x, 5xpa, 6cy4, 4w92, 3i5y, 5jc9, 4kqy, 5hd1, 3zd5, 4v9b, 5vi5, 3bnq, 3mdi, 4xw1,
4jxz, 4lj0, 5dar, 4gv9, 3bx2, 4v5q, 4e6b, 3ok2, 4wzd, 5e17, 1qvf, 405d, 4rmo, 3i61, 4kq0,
3g71, 3i5x, 3d2g, 3ciy, 5zem, 5k77, 4nyc, 5usa, 5ndi, 5ddp, 5xut, 5fdv, 3ol6, 2byt, 3tra,
5z71, 5ef0, 5f8g, 1i9x, 2pxf, 3b5s, 4yb0, 3f2w, 3ola, 3ktw, 3gao, 5fk5, 4tna, 406d, 4z4h,
1zz5, 5t8y, 3f4e, 2jja, 4g0a, 5xz1, 2qk9, 2q1r, 6dlq, 1f7y, 3avy, 6m7k, 1aq4, 2npy, 4kr9,
6dmc, 6evk, 4dzs, 2gcs, 5t7b, 5f5h, 5g4v, 3ccj, 3dw4, 4kxt, 1i6h, 2hoo, 1vqn, 2et3, 5bud,
3pu1, 4wr6, 5wzk, 6db8, 1hr2, 3q0n, 1fxl, 5ytv, 4k4y, 6dcl, 6dpf, 4ngg, 4z4g, 5ktj, 1j5e,
3iev, 6blp, 4w5t, 4lsk, 1un6, 2p7e, 3wqz, 2oe6, 1gts, 5o1y, 4ht8, 4xwf, 1ik5, 2xd0, 6cxz,
3snp, 3cf5, 3hga, 4yb1, 3la5, 4w2f, 3koa, 5v0k, 6bqf, 1zl3, 3wfr, 1ntb, 3s7c, 4aq7, 4v50,
6dpl, 4q5v, 4s3n, 5ay4, 4a93, 5j4c, 4v8g, 1j9h, 5jea, 3e2e, 3nd4, 3nkb, 6dp8, 1qc0, 1j6s,
6d30, 1dk1, 3qgb, 5zei, 1x9c, 4lvv, 3aev, 2dvi, 5ued, 3bno, 3lrn, 1n78, 5ip2, 5wzi, 5v7c,
6dpg, 5w5h, 1ytu, 6gsk, 3wzi, 5axn, 6htu, 2zh8, 4bwm, 4k4s, 3ccl, 2z75, 1eqq, 1duh, 4e8k,
5wjr, 2bcy, 3g96, 1nbs, 4v8h, 1l3z, 1yzd, 5o62, 5i9f, 5wty, 4b3o, 5yki, 4dv7, 5jxs, 1m5o,
4u3u, 2cv2, 3cz3, 6mj0, 1fir, 3ccs, 5wns, 4erl, 4zcf, 5do5, 2pxp, 5on3, 1y73, 4ol8, 3l3c,
3akz, 2jea, 2r7r, 1p79, 4qil, 1b7f, 5fw3, 3f30, 1q82, 5nom, 5d5l, 2bgg, 4dr6, 5dh7, 3cpw,
4xco, 3bnn, 1duq, 430d, 5gmg, 4x64, 1gax, 5d99, 5kk5, 4nha, 1y69, 1vq7, 1n1h, 2g92,
1tra, 5zeg, 3cw5, 2jlt, 3hou, 5xh6, 2ply, 5tpy, 3s1m, 3avx, 3skz, 5une, 1f1t, 2hoj, 6fq3,
2izm, 2xlj, 5dh6, 2zha, 3po2, 3erc, 3s2h, 6hbx, 2o3y, 5swm, 5j8b, 3q0o, 3ol8, 2nug, 1yyo,
5gxi, 4x4s, 1kuq, 2xdd, 3ds7, 3vrs, 3s16, 2uwm, 5hcq, 4p3u, 4ilm, 1ffk, 4jyz, 3owz,

3mum, 5d0a, 1fuf, 3avu, 6bsj, 4jxx, 4mgm, 6dp3, 1vbz, 2oih, 1g4q, 6dow, 4u4y, 3pip, 3s49, 2xpj, 4a3e, 5i4a, 1wsu, 2b2e, 5js1, 4u50, 3icq, 3k61, 5wzj, 157d, 5ng6, 1euq, 4wta, 4lgt, 6cfl, 6f4g, 3gx6, 1sds, 1mji, 4jrt, 4z31, 4ghl, 1ibl, 1kh6, 5vsu, 6bgb, 1jbt, 4n2s, 6nd6, 3g8t, 3syw, 4x4p, 3diz, 364d, 4fe5, 2r22, 4rne, 4v7s, 3a3a, 2drb, 4fsj, 4wal, 3d0m, 3r9x, 5mrx, 5wit, 4q5s, 2db3, 5l4o, 5o7h, 3iwn, 1utd, 4tvx, 4bpb, 2et5, 6gvy, 2fd0, 5vo8, 5f8h, 3q0s, 2bs1, 1kfo, 5ccx, 4wkr, 3sj2, 5hsw, 4qu6, 3dvv, 5wnv, 2pxd, 5fj4, 6b14, 2py9, 5fkd, 4tue, 4wc2, 4fwt, 2hhh, 1wpu, 3ouy, 2b2g, 4gg4, 1drz, 1m5v, 4v8a, 1zev, 1pjg, 1qrt, 2gq7, 3q0r, 1qu2, 472d, 4dr3, 1cvj, 4k4w, 5w51, 5udl, 5wwt, 5fk6, 4kr7, 5z1i, 3mja, 4j39, 5y7m, 1kc8, 6doq, 2grb, 3jxq, 2xs2, 2j0s, 3cgq, 4gv6, 2oeu, 4w5o, 4k4z, 2du3, 4v8f, 4enb, 1kog, 2f8k, 2pxk, 4ktg, 300d, 3siu, 2gxb, 4rc0, 1exd, 3ucu, 6car, 4jv5, 5elk, 3dis, 6d92, 4faw, 4lf7, 2e9t, 4wtm, 4ato, 3gca, 1vq4, 3cxc, 3m3y, 5zth, 3k62, 5m3h, 3mur, 6b3k, 6cc3, 1kq2, 4gpw, 6az4, 5ob3, 4un5, 1xpu, 2azx, 4k4x, 1z43, 3pf4, 5e6m, 5dv7, 1q2r, 5mga, 1q2s, 3pey, 5jbg, 5zsl, 1rpu, 3ow2, 6dls, 2v7r, 5d6g, 4bhh, 6nof, 6doh, 2bq5, 4wtj, 4tz6, 5fmz, 4a3f, 4jah, 4pqv, 5use, 1a9n, 3skw, 6d9k, 5xwg, 5zsa, 1yjw, 6c6k, 3fo6, 4nmg, 2nue, 4k4u, 3fht, 1z7f, 2hyi, 4g6p, 5lm7, 1yyw, 1b23, 6f4h, 5us2, 2pxt, 4qqb, 1uvj, 4l8r, 5vj9, 3ho1, 4u20, 3iqr, 5mei, 5czp, 1y3s, 2o5i, 6dob, 4fvu, 2pxb, 6ar3, 2e2j, 421d, 5onh, 5j88, 4wtk, 3oxj, 5eez, 5ef2, 5eaq, 5hbw, 4f02, 3x1l, 4v8x, 2et8, 5wwx, 1il2, 2oue, 1dfu, 4lvw, 3i56, 4u27, 5dea, 4h8k, 6dpi, 3lwo, 4wqy, 3k1v, 3slq, 4qm6, 2anr, 3bnp, 4q0b, 3owi, 5ytx, 6dof, 1vqo, 4tra, 5eey, 3zp8, 2atw, 5wwg, 5hbx, 4nyb, 2xs7, 4alp, 2r7w, 1yy0, 5t83, 2b2d, 4rgf, 3rc8, 1a34, 1hnz, 2a1r, 5f0q, 1njp, 2hvy, 4v85, 433d, 2zh6, 4ijs, 1vq8, 2e9r, 3rer, 462d, 4puo, 1zh5, 1fg0, 3qrp, 357d, 5cki, 6doa, 1k01, 5x22, 1xnq, 3f73, 1xbp, 2von, 4fb0, 6dog, 5bjo, 4zlr, 5udi, 5uj2, 3gs1, 2xzl, 2p7f, 5wti, 3loa, 4u37, 3dim, 4rcm, 3d0u, 3s15, 1lc4, 4v7u, 4lvy, 397d, 4wc4, 5z9x, 5mfx, 1gtr, 4ifd, 3ol9, 1rna, 4p3t, 6gyv, 5dh8, 3klv, 1ykv, 4l81, 3p4d, 5e08, 3qjl, 4w4g, 5der, 5j5b, 3gib, 2zzm, 2be0, 2et4, 3h5y, 5v3f, 2o3x, 5zsm, 5e3h, 5o69, 6d2z, 1m90, 3s1q, 3kfu, 3qsy, 1ddy, 4gcw, 6h9i, 301d, 2uxb, 1yjn, 5a0t, 4f1n, 4e78, 4c4w, 4www, 2du6, 4jng, 5v1l, 2eew, 4x4v, 5ndh, 5jji, 1qu3, 4u1v, 3qjp, 4e8r, 4xw0, 5hcp, 2fcx, 4en5, 5bte, 5n94, 3dvz, 5awh, 3bsu, 3agv, 3sd3, 1vtq, 2zhb, 4ill, 3qjj, 5w7o, 4jrc, 2gq4, 1asz, 5fci, 2g9c, 3q3z, 5ztm, 2pn3, 6dp6, 5czz, 2ec0, 2quw, 1q86, 3nj7, 4xjn, 2xo0, 5w6v, 6c8k, 6fpx, 5m73, 2ix1, 1wrq, 3moj, 3fs0, 5guh, 3bsb, 1i7j, 4b3t, 2qwy, 2g8i, 5v9z, 4oi1, 5ns4, 3amu, 6ifn, 1j1u, 4u55, 3fte, 6ifo, 5fkf, 4fxd, 2ppb, 6dp2, 4un3, 5w1i, 353d, 5z4a, 1di2, 420d, 2r7y, 280d, 1gsg, 5h3u, 2g3s, 4v88, 434d, 3t5q, 5xc6, 2xnz, 5e54, 1d4r, 1yi2, 4un4, 4c9d, 5tgp, 2b57, 5zsn, 4w5r, 4nia, 6doc, 1l9a, 4dr5, 4v2s, 2eet, 1zdh, 3k5y, 4u3m, 3ok4, 3bt7, 4pqu, 5j01, 4n48, 1ehz, 3lwq, 4v4h, 4rcj, 3slm, 1vy4, 6cfk, 4bxx, 4u47, 3olb, 1qcu, 5eew, 4lt8, 5ndk, 1vc6, 5f8i, 4ohy, 4qi2, 3hax, 4v9f, 4ycp, 4u26, 4v9h, 3f2q, 3oxb, 4v83, 2zjr, 1ibm, 5nxt, 1zse, 1ec6, 4u6k, 409d, 3p4b, 6dn1, 1m8w, 3q0m, 6g2k, 3td0, 2r20, 4jvh, 4w2g, 6i1k, 2oe8, 1yls, 4v5j, 3trz, 4wtg, 5bxk, 1xmo, 5tko, 1ze2, 3pew, 5kal, 4r4v, 2b3j, 1wvd, 4z92, 4wkj, 3p4a, 1r9f, 4prf, 4wsd, 283d, 6dor, 4v51, 205d, 4mgn, 6noh, 1vqk, 5fk4, 1i94, 6boh, 3oxe, 3k4e, 5u3g, 1vc5, 3m7n, 5dno, 6cas, 1y26, 6dok, 1w2b, 2zuf, 3nma, 1tfw, 2oij, 4rkv, 1feu, 1u1y, 3nna, 1uvn, 2zh1, 3zjt, 2zh5, 3ud4, 3adl, 6fkr, 1vfg, 4xbf, 1u9s, 5w7n, 6g7z, 2uu9, 4frn, 5dgv, 2zzn, 3s14, 387d, 4fnj, 1h3e, 2g8w, 6b4v, 5nwq, 6d8l, 3vjr, 2g8u, 4e58, 4ay2, 6dpj, 2bs0, 5yts, 2xo1, 6fhi, 2v6w, 3dw6, 5k8h, 6i0v, 2iz9, 5krg, 1mdg, 4ngb, 5xuz, 1qbp, 4nh5, 3zc0, 1xok, 1p9x, 3f4g, 5xog, 3tzr, 4wro, 5elx, 4v9q, 1rlg, 3ox0, 4wtc, 2f4s, 4b3s, 4v7x, 4wt8, 4a3g, 3ncu, 4wrt, 2oe5, 2gdi, 5dc3, 5c45, 3cgp, 6gsj, 3b91, 3gx2, 5elt, 1m8v, 2dxi, 3npn, 2xlk, 5wwr, 1sj4, 4nya, 5nqi, 5ibb, 1qrs, 3kms, 2o5j, 4x4n, 5zsd, 4yye, 5d0b, 5f6c, 5w5i, 6cf2, 5eao, 5ah5, 3g6e, 4iqs, 2p7d, 6c8n, 4o8j, 4v9n, 3gx5, 2zue, 3skl, 3sux, 2o3w, 5jch, 4u25, 3mij, 6cy2, 4v9c, 4g9z, 4yvj, 354d, 5fke, 1o9m, 1h4s, 4aqy, 2eeu, 2oj0, 4nh3, 6dpc, 4wfm, 4qlm, 4zdp, 4wc7, 2gpm, 1qa6, 4qvc, 3ptx, 4m7a, 6dp5, 1y99, 3ccr, 5gip, 2cv0, 5dm6, 4oi0, 2ab4, 5zw4, 5swe, 1yyk, 5y58, 1j7t, 5emf, 3bwp, 4nh6, 6dpb, 4f8u, 1wmq, 4k4t, 3cma, 5i9g, 2pwt, 5xow, 1nlc, 5do4, 4v8c, 3oww, 3t3n, 1fix, 4j7l, 1y90, 4wzq, 4erj, 3e5f, 2fcy,

4x66, 1yxp, 4ngf, 2nz4, 5dto, 4w2i, 4y7n, 4rj1, 2jly, 5en1, 4ari, 4l8h, 4gha, 3t1h, 5on6, 5k36, 4tzy, 1f7u, 5y87, 4v7v, 1m1k, 6dn2, 3dw5, 2gcv, 3b4c, 2otj, 4wpo, 1j5a, 1yrj, 2x1f, 2rd2, 4cs1, 6bfb, 3hjf, 6dta, 4qik, 6aso, 2zi0, 4nku, 5f8l, 3o6e, 2dqq, 3qsu, 3b4b, 3pu0, 3diy, 4enc, 4v8e, 1vql, 5kvj, 4tz0, 5wnq, 5dcv, 1xpo, 5a0v, 4wsb, 2dr9, 2bx2, 4y91, 1yz9, 5tf6, 5nex, 4u4r, 3cul, 4nyd, 6hu6, 5uz6, 413d, 2y8w, 1jid, 5dge, 4wfb, 3egz, 4ts2, 4y52, 1y3o, 5tbw, 5hp3, 6cy0, 4e48, 1euy, 6ar5, 3knc, 4e8n, 3mdg, 6d12, 3zd6, 4mce, 4pmi, 4v52, 6d1v, 2yie, 2az0, 3qgc, 5k78, 4j7m, 3pio, 6doz, 5vci, 4wte, 2der, 1aq3, 2jlv, 3diq, 3fo4, 4v4q, 1g2e, 3e5c, 1q9a, 4e8t, 5www, 3jxr, 2ydh, 1mme, 2qkk, 4x62, 3oxm, 4wqr, 4f3t, 479d, 4jzv, 469d, 1q81, 1msw, 299d, 5c7w, 2pxv, 4p43, 3b0u, 4xw7, 7msf, 6dp7, 1jzy, 2hw8, 4w5q, 2dlc, 5vgw, 377d, 3cc7, 2gtt, 4v7w, 2dqp, 2r7v, 2g8k, 1yvp, 5br8, 2h0z, 5ho4, 3czw, 5fj0, 1mhk, 4e8p, 4qka, 3am1, 1nwx, 2fqn, 4p9r, 3i2q, 3ski, 4v9a, 4w5n, 3zjv, 3g8s, 5o1z, 4wti, 3po3, 5xtm, 2a04, 3add, 3zgz, 4p3s, 1g1x, 5wws, 2xgj, 2xzo, 5hnq, 1si3, 3s8u, 2qex, 6aay, 5f9r, 3adc, 4oji, 6c63, 1n38, 5w4k, 3wbm, 3q50, 2yjy, 4kr6, 3i62, 5m0h, 6fpq, 4u3l, 5u0q, 3cr1, 6dpd, 6du5, 4fep, 5det, 3iab, 4qoz, 1sj3, 1zho, 2gis, 5fj1, 3e5e, 4yzv, 4znp, 5udj, 2d3o, 1asy, 4v87, 5fq5, 1rxa, 3i2s, 1k8a, 3boy, 3gog, 3umy, 5tdk, 1m5p, 6i1l, 2awe, 4n0t, 5u34, 404d, 4pwd, 5x21, 2uub, 3f2y, 3hl2, 1kqs, 5ay3, 4feo, 2ake, 5di2, 4yn6, 3q0p, 470d, 6b0b, 4lx6, 3bnr, 3w3s, 5wis, 5eex, 4jgn, 3igi, 4qjd, 2bbv, 1jbs, 3far, 5ccb, 4hor, 5b2r, 1zfv, 5hby, 4aob, 4qei, 2gun, 2c50, 1f27, 4ioa, 4pdb, 2qux, 5voe, 3s1r, 4boc, 5on2, 4v8d, 5v8i, 5hab, 1yhq, 1ser, 1u63, 438d, 6doj, 3d2x, 1jzv, 2il9, 1b2m, 4iqx, 4nxh, 1q29, 1zbh, 1qln, 3mxh, 3k0j, 1xpr, 1kd3, 4rzd, 3bns, 5de5, 5ef3, 4jf2, 3dj2, 3p4c, 3wru, 5ml7, 3ovb, 5wtk, 435d, 5f98, 6qic, 6c64, 1dqh, 2izn, 4gpy, 5nz6, 2w2h, 4wt1, 5kx9, 2vqe, 4z4c, 2az2, 6dvc, 3pdr, 4wtl, 4plx, 6c8i, 5xbl, 4h5p, 3nmr, 4ig8, 5c0y, 2zh2, 3m85, 5f8k, 6c8j, 1zdj, 3rkf, 1f8v, 3q2t, 4rge, 3r9w, 5ch0, 2g8f, 4u4q, 4ht9, 4rum, 3f2x, 1fjg, 4tyy, 1ffz, 2wna, 6bsg, 4k50, 3af6, 5jc3, 2e2i, 2npz, 2cv1, 2bee, 1kxk, 1yj9, 4v57, 5ib8, 5nz3, 2oiy, 4w2e, 3b58, 2qus, 3sqw, 5w0o, 3g9y, 4uyk, 6d8a, 1dqf, 4ts0, 2iy5, 1pvo, 6cc1, 5v3i, 1zbi, 2xli, 5zx2, 6dpa, 3cjz, 3b5f, 2gje, 4v5l, 3mei, 4wra, 4u51, 3o3i, 5hcr, 5fw1, 3ftf, 5oc6, 3cc4, 4pjo, 4ena, 5z1h, 5f8j, 5jrc, 5lr3, 1osu, 3bsx, 6bjg, 5axw, 4py5, 2bj6, 3g4s, 1kd1, 1m8y, 2esi, 4csf, 3siv, 5ed2, 4v5r, 2dra, 1s03, 4nfq, 4a3l, 5w1h, 2gq6, 4jiy, 1ond, 5g4t, 4rwn, 4v9e, 4db2, 4y1i, 5u31, 3epk, 5gxh, 2v3c, 4qvi, 3rzd, 5xj2, 4qk9, 6e4p, 6ar1, 6dpk, 5vpo, 3bbi, 6fz0, 3k49, 6d8o, 4dv6, 1rc7, 4hkq, 1msy, 1u6b, 4erd, 3o8c, 5zse, 4y4p, 3v74, 3vnv, 1ibk, 5di4, 1hnw, 5jjk, 4lx5, 4y1j, 4rbq, 3l0u, 3gs8, 4faq, 4j50, 4jzu, 333d, 4rby, 3nvk, 4kji, 1zci, 4oo1, 3dio, 2pxl, 4meg, 1ffy, 4nfo, 6dp9, 3lqx, 1efo, 5v0o, 4pmw, 5vof, 4wq1, 5amq, 1qvg, 3q0q, 2vpl, 6cao, 6buw, 5zc9, 1vbx, 6dpo, 5jjl, 6dcb, 4far, 4ybb, 4lck, 3rzo, 3lwp, 2q1o, 1bmv, 3s17, 6ck5, 3b31, 2tra, 2e5l, 3ts2, 5zq1, 4khp, 3b4a, 3epj, 6bjy, 2otl, 5nrg, 437d, 3nnh, 2bh2, 1n32, 5y6z, 5ueg, 2bu1, 4e8m, 4c40, 5emo, 6hhq, 4k4v, 3bbk, 2nqp, 1vby, 5ew7, 1dul, 3u56, 4kz2, 3d2v, 2nvt, 5lyb, 3dw7, 2wj8, 1xpe, 1r9t, 4wsm, 6nod, 4v9s, 3ccq, 1k9w, 2jlx, 6do8, 5swd, 5wt3, 4v7t, 3zla, 2pxq, 1kd4, 5elh, 4fau, 4hos, 4j1g, 4rwo, 2ao5, 2a43, 5xpg, 3td1, 2zh4, 3dd2, 1l2x, 5j4b, 6cbd, 3hhn, 5v0j, 4xk0, 5h1k, 1zzn, 3bnt, 6doy, 1z79, 2uuc, 4ang, 1sm1, 3dj0, 1hmh, 3ov7, 5fkg, 5ki6, 3t3o, 4wc5, 5o58, 4wc3, 4oo8, 5c9h, 3ger, 1n77, 1f7v, 5l2l, 2dr2, 6fql, 2g8h, 4kr2, 3k64, 2v0g, 4u38, 4oq8, 3sn2, 5new, 3nvi, 3d0x, 2jlu, 4fts, 5lyv, 402d, 2o3v, 1uvl, 2hok, 6dlt, 4u6m, 4cqn, 4i67, 3ndb, 4b3g, 2uxd, 3ie1, 4gkj, 1urn, 2r7t, 1nb7, 4fel, 1s76, 6a4e, 6cfj, 4qk8, 6gd2, 6gx6, 1g2j, 3avv, 3i2u, 5dhb, 2vnu, 1h4q, 3bso, 2g4b, 6hbt, 1vq9, 2du4, 3mj3, 4kzd, 6bsh, 5vp2, 6e0o, 5el4, 4q9r, 3s1n, 3ts0, 5ny8, 2bny, 6db9, 1rxb, 4w90, 4nyg, 3fu2, 2oj3, 5tga, 4fej, 468d, 5xuu, 5o6u, 4z4f, 4kze, 6cu1, 4ji0, 488d, 3qrr, 5c7u, 6ck4, 6caq, 1l8v, 3al0, 1y27, 2zjq, 1t0e, 6fhh, 3ivk, 4v7m, 4fte, 5obm, 3r2c, 5xh7, 4v9k, 5vjb, 4zt0, 5vm9, 4z4i, 1fka, 5f8m, 5ddq, 6cyt, 4qyz, 5neq, 5sze, 5jc7, 4meh, 5ndv, 5fk1, 5g4u, 5x6b, 4v63, 5c5w, 1cwp, 2x1a, 2dqo, 3ivn, 5ot2, 4afy, 4f8v, 3t5n, 6d95, 1p6v, 3hov, 5dat, 4zdo, 3nmu, 4v84, 1zfx, 1vq5, 464d, 4qpx, 2qkb, 5axm, 3h5x, 5ws2, 6dp1, 4al7, 1t0d, 5bzv, 4o41, 3gx7, 3bx3, 4tv0, 4am3, 4qvd, 5m0j, 5j30, 3adi, 6d9l, 5gin, 6dot, 6don, 3dir,

4y1n, 2fgp, 5e7k, 4g6s, 1lng, 5z4d, 6cmn, 5h9f, 5fkh, 4p20, 4q9q, 4gpx, 1et4, 4jk0, 3r1d, 1lnt, 5wzh, 1gtf, 3ahu, 3r4f, 4v8b, 4al6, 2zxu, 3ex7, 2qbz, 4v9d, 5mwi, 1jj2, 2oiu, 5i9d, 4tzx, 2xdb, 5u30, 3cd6, 3ol7, 4p95, 6bz6, 1knz, 4nlf, 4j5v, 6msf, 3sqx, 4z4d, 3r1l, 5jaj, 2ho6, 255d, 1qru, 2rfk, 2r8s, 4oe1, 5dgf, 5x70, 4pkd, 4y1o, 1t0k, 2csx, 3hm9, 6bjh, 5uza, 3dh3, 1hq1, 1zbl, 2hop, 5j91, 6c8m, 4y1m, 4qg3, 5f9f, 4u7u, 1n8r, 4pei, 4v54, 2c4q, 5f9h, 5hnj, 3bo2, 2zko, 2d2k, 3iqn, 5m3j, 4g6r, 422d, 3vyy, 439d, 2nok, 2zni, 6dp0, 4p3e, 4wf9, 4m30, 4m6d, 3htx, 5wnt, 4lg2, 1jbr, 3pkm, 6dox, 5t3k, 6c8o, 3twh, 1yfg, 3zju, 2a8v, 5b2s, 6i0u, 5m0i, 6bbo, 4mdx, 4u8t, 1vy6, 5mjv, 4x4t, 5eim, 2b8r, 1m5k, 1y95, 1zft, 1nuv, 3rg5, 2y8y, 3g4m, 6gc5, 6dmd, 4msb, 3nd3, 4knq, 4jrd, 3dll, 6gpg, 4x4o, 2xc6, 2g91, 2r21, 5udz, 5aox, 6i0t, 3ks8, 2vum, 1l3d, 2val, 2y9h, 1g59, 3qg9, 4ypb, 3r2d, 1vy7, 5x2h, 3b5a, 4wsa, 4u1u, 6dov, 1br3, 5lr5, 4uyj, 5v2h, 2yu9, 5btp, 4u3r, 4yvk, 4wan, 1sjf, 398d, 6d8f, 4z3s, 4c8z, 2h0x, 4zld, 2i82, 4u53, 1h38, 5ex7, 3p59, 4d26, 5j7l, 3nl0, 1q7y, 5vw1, 3gx3, 4v90, 6i7v, 4ba2, 1zdk, 4pco, 3adb, 4by7, 6f3h, 4x4q, 4b3r, 2fk6, 2vuq, 6dme, 6hct, 3r1c, 2bte, 5ddo, 2xbm, 2d2l, 5kla, 6dcc, 4ejt, 1vc0, 2voo, 3suy, 1uvm, 4peh, 3ucz, 1ivs, 1i5l, 2hol, 6cb3, 4oav, 3oxd, 5da6, 429d, 3i55, 4oog, 4ftb, 5cd4, 5v0h, 6d06, 4c8y, 4a3b, 1xpf, 2hom, 3v71, 6dph, 2gq5, 2h1m, 2asb, 4v7l, 3nj6, 5hp2, 4v7y, 2aar, 5yyn, 4wj4, 5bjp, 5neo, 3g0h, 1ob2, 5ib7, 3dig, 4v9p, 4x65, 1nyi, 3lwv, 4z7l, 4v5k, 4u78, 1vvj, 2uua, 4v9r, 1qf6, 5gmf, 4rbz, 2i91, 2jlz, 5uee, 2ct8, 6dpe, 5b43, 1k8w, 2ez6, 2c4z, 6nd5, 3ssf, 5lys, 6hau, 6dom, 4m59, 3g78, 1s0v, 2eev, 2a0p, 5vaj, 3ice, 1o0c, 1xnr, 6dp4, 4n2q, 6bjv, 6blo, 6ai6, 5d8t, 5d8h, 5el7, 6dop, 5z98, 3pla, 5f8n, 1gid, 3eoh, 5z4j, 4qln, 3dix, 5hk0, 3ud3, 5msf, 5lqo, 5j3c, 1u0b, 480d, 4lf6, 4v9i, 1nuj, 4l47, 1i6u, 1j2b, 3hsb, 4v6g, 2iz8, 5uk4, 1nkw, 5fcj, 310d, 5xus, 4v8i, 2xsl, 5dm7, 4ii9, 361d, 4lf9, 3gm7, 4b3m, 3zd7, 4ms9, 4r8i, 6bsi, 4u4n, 3hhz, 6gd3, 4lvx, 5h1l, 5j02, 1hnx, 4msr, 6is0, 4u3n, 4lq3, 1eiy, 4nl3, 4v7z, 4jvy, 5hjz, 3oin, 3u4m, 2pxu, 5ddr, 2gic, 1pjo, 4e5c, 3avt, 5cnr, 4jnx, 3eqt, 3suh, 1ykq, 5i4l, 4pgy, 6tna, 2zy6, 5zej, 2fmt, 4mcf, 1ob5, 2re8, 3got, 4v6a, 5ux3, 4s2y, 5wnp, 6doi, 2d6f, 3lwr, 4o26, 1o3z, 6doe, 1sdr, 3og8, 2c0b, 5v1k, 6d8p, 5wqe, 3f4h, 4v5e, 6dt8, 2nuf, 4arc, 4tyw, 5dhc, 6cae, 4yhw, 5js2, 471d, 3cc2, 5ux0, 5b2t, 5ew4, 4gv3, 5o3j, 4v5d, 4m4o, 3gtj, 5yze, 4wc6, 4u52, 3vnu, 2j0q, 3rw6, 4x9e, 4lmz, 4yco, 4xnr, 3k5z, 6dpn, 6dtd, 4u6l, 1efw, 1jb8, 1n33, 3mut, 1h2d, 1i9v, 2fz2, 2nvq, 359d, 4x4u, 6do9, 2r1s, 3g9c, 3tup, 5els, 2po1, 4ji3, 5el5, 5x2g, 1laj, 5h9e, 4tu0, 1mwl, 5hc9, 2e9z, 3ovs, 2det, 5lyu, 4qu7, 2ann, 1q96, 2du5, 3iqp, 5hau, 4k27, 1jzx, 5vzj, 5nep, 2pxe, 3pf5, 5el6, 3amt, 5fk3, 4lw0, 4lf4, 4v8n, 3bnl, 5sup, 3cme, 4jab, 4d25, 4yhh, 5doy, 5ef1, 5wea, 4z8c, 3eog, 4olb, 379d, 5vcf, 3f2t, 6dou, 1o0b, 4v5a, 4a3c, 1nwy, 4k32, 5tsn, 3muv, 4v99

**Table S2.** Percent occurrence frequency of three most dominant topologies of purine||purine stacks.

| **Topology** | **Percent occurrence** |
|---|---|
| 5‖5, 6‖56 α‖β cis | 24.2% |
| 5‖6 α‖β cis | 17.5% |
| 6‖6 β‖β cis | 15.8% |

**Table S3.** List of purine||purine stacking topologies that can occur only at extreme σ values (close to 90°).

| Face | Topology | Glycosidic Orientation | Base | consecutive/non-consecutive |
|---|---|---|---|---|
| α\|\|α | 5\|\|5, 6\|\|6 | *cis* | A\|\|A | consecutive |
| | | | | non consecutive |
| | | | A\|\|G[a] | non consecutive |
| | | | G\|\|G[a] | non consecutive |
| | 5\|\|6, 6\|\|5 | *trans* | A\|\|A | consecutive |
| | | | | non consecutive |
| | | | A\|\|G | consecutive |
| | | | | non consecutive |
| | | | G\|\|G | consecutive |
| | | | | non consecutive |
| α\|\|β | 5\|\|6, 6\|\|5 | *cis* | A\|\|A | consecutive |
| | | | | non consecutive |
| | | | A\|\|G | consecutive |
| | | | | non consecutive |
| | | | G\|\|G | consecutive |
| | | | | non consecutive |
| | 5\|\|5,6\|\|6 | *trans* | A\|\|A | consecutive |
| | | | | non consecutive |
| | | | A\|\|G | consecutive |
| | | | | non consecutive |
| | | | G\|\|G | consecutive |
| | | | | non consecutive |
| β\|\|α | 5\|\|6, 6\|\|5 | *cis* | A\|\|G | consecutive |
| | | | | non consecutive |
| | 5\|\|5, 6\|\|6 | *trans* | A\|\|G | consecutive |
| | | | | non consecutive |
| β\|\|β | 5\|\|5,6\|\|6 | *cis* | A\|\|A | consecutive |
| | | | | non consecutive |
| | | | A\|\|G | consecutive |
| | | | | non consecutive |
| | | | G\|\|G | consecutive |
| | | | | non consecutive |
| | 5\|\|6, 6\|\|5 | *trans* | A\|\|A | consecutive |
| | | | | non consecutive |
| | | | A\|\|G | consecutive |
| | | | | non consecutive |
| | | | G\|\|G | consecutive |
| | | | | non consecutive |

[a]Of all the stacking types listed in this table, one example each of these two stacks were found in RNA crystal structures, albeit at high σ values (83.5° (A||G) and 85.3° (G||G)) See Table S12 for examples of these two topologies.

**Table S4.** List of purine||purine stacking topologies that are possible at reasonable σ values, but not found in RNA crystal structures.

| Face | Topology | Glycosidic Orientation | Base | consecutive/non-consecutive |
|---|---|---|---|---|
| α‖α | 5‖56, 6‖5 | *trans* | A‖A | consecutive |
| | | | | non consecutive |
| | 5‖56 | *trans* | A‖A | consecutive |
| | | | | non consecutive |
| α‖β | 5‖56, 6‖6 | *cis* | A‖A | consecutive |
| | | | | non-consecutive |
| | | | G‖G | consecutive |
| | | | | non-consecutive |
| | | *trans* | A‖A | consecutive |
| | | | | non-consecutive |
| | | | G‖G | consecutive |
| | | | | non-consecutive |
| | 6‖56 | *cis* | A‖A | consecutive |
| | | | | non-consecutive |
| | | | G‖G | consecutive |
| | | | | non-consecutive |
| | | *trans* | A‖A | consecutive |
| | | | | non-consecutive |
| | | | G‖G | consecutive |
| | | | | non-consecutive |
| | 5‖5, 6‖5 | *cis* | A‖A | consecutive |
| | | | | non-consecutive |
| | | | G‖G | consecutive |
| | | | | non-consecutive |
| | | *trans* | A‖A | consecutive |
| | | | | non-consecutive |
| | | | G‖G | consecutive |
| | | | | non-consecutive |
| | 5‖6 | *cis* | A‖A | consecutive |
| | | | | non-consecutive |
| | | | G‖G | consecutive |
| | | | | non-consecutive |
| | | *trans* | A‖A | consecutive |
| | | | | non-consecutive |
| | | | G‖G | consecutive |
| | | | | non-consecutive |
| | 5‖56 | *trans* | G‖G | consecutive |
| | | | | non consecutive |
| β‖β | 5‖56,6‖5 | *trans* | G‖G | consecutive |
| | | | | non consecutive |

**Table S5**. Examples (i.e. identities of bases involved in stacking and corresponding PDB codes) of consecutive (black) and non-consecutive (grey) 5||56, 6||56 α||α *cis* purine||purine stacks. The geometrical parameters ($\vec{d}_{ab}$, $\theta_{ab}$, $\tau_a$, $\tau_b$ and $\sigma_{ab}$) averaged for each ring:ring stacking contact (i.e. average parameters from 5||5, 5||6, 6||5 and 6||6 ring:ring contacts) for each example stack are provided in parentheses respectively.

| | | α face (56), (56) | |
|---|---|---|---|
| | | A | G |
| α face (5), (6) | A | A16(c)\|\|A17(c); 4w90<br>(3.8 Å, 15.4°, 22.4°, 20.1°,9.1°) | A900(A)\|\|G901(A); 6hhq<br>(3.8 Å, 13.8°, 28.9°, 20.1°, 15.8°) |
| | | A26(A)\|\|A28(A); 6d3p<br>(3.9 Å, 4.9°, 24.1°, 25.3°, 29.1°) | G31(D)\|\|A32(D); 3nmu<br>(3.9 Å, 20.0°, 27.2°, 16.9°, 4.9°) |
| | G | | G165(a)\|\|G164(a); 5j8b<br>(4.0 Å, 21.9°, 25.6°, 17.4°, 8.8°) |
| | | | G6(A)\|\|G36(A); 4lvv)<br>(3.8 Å, 4.0°, 22.2°, 22.2°, 0.5°) |

**Table S6**. Examples (i.e. identities of bases involved in stacking and corresponding PDB codes) of consecutive (black) and non-consecutive (grey) 5||6, 6||56 α||α *cis* purine||purine stacks. The geometrical parameters ($\vec{d}_{ab}$, $\theta_{ab}$, $\tau_a$, $\tau_b$ and $\sigma_{ab}$) averaged for each ring:ring stacking contact (i.e. average parameters from 5||6, 6||5 and 6||6 ring:ring contacts) for each example stack are provided in parentheses respectively.

| | | α face (6), (56) | |
|---|---|---|---|
| | | A | G |
| α face (5), (6) | A | A17(w)\|\|A16(w); 2gtt<br>(3.7 Å, 12.1°, 15.6°, 21.5°,3.3°) | A1287(DA)\|\|G1288(DA); 4u24<br>(3.8 Å, 15.6°, 21.5°, 16.6°,7.0°) |
| | | A1127(DB)\|\|A2518(DB); 4v54<br>(3.9 Å, 10.0°, 23.1°, 18.4°,42.5°) | A933(ya)\|\|G931(ya); 4w4g<br>(3.8 Å, 6.4°, 22.7°, 23.6°, 21.4°) |
| | G | | G390(2)\|\|G389(2); 4u6f<br>(3.7 Å, 11.7°, 18.1°, 19.2°, 4.1°) |
| | | | G1483(a)\|\|G904(a); 4k0k<br>(3.6 Å, 7.6°, 16.1°, 17.1°, 7.6°) |

**Table S7**. Examples (i.e. identities of bases involved in stacking and corresponding PDB codes) of consecutive (black) and non-consecutive (grey) 5||5, 6||56 α||α *cis* purine||purine stacks. The geometrical parameters ($\vec{d}_{ab}$, $\theta_{ab}$, $\tau_a$, $\tau_b$ and $\sigma_{ab}$) averaged for each ring:ring stacking contact (i.e. average parameters from 5||5, 6||5 and 6||6 ring:ring contacts) for each example stack are provided in parentheses respectively

| | | α face (5), (56) | |
|---|---|---|---|
| | | A | G |
| α face (5), (6) | A | A1085(DA)\|\|A1086(DA); 4v63 (4.1 Å, 23.0°, 22.2°, 34.2°, 2.8°) | G973(AA)\|\|A974(AA); 6i7v (4.4 Å, 22.5°, 24.5°, 22.1, 77.7°) |
| | | A-5(D)\|\|A-7(D); 2vod (3.9 Å, 12.0°, 23.9°, 19.7°, 65.9°) | |
| | G | | G1024(CA)\|\|G1025(CA); 6i7v (4.6 Å, 17.3°,21.3°, 29.8°, 75.8°) |

**Table S8**. Examples (i.e. identities of bases involved in stacking and corresponding PDB codes) of consecutive (black) and non-consecutive (grey) 5||56, 6||6 α||α *cis* purine||purine stacks. The geometrical parameters ($\vec{d}_{ab}$, $\theta_{ab}$, $\tau_a$, $\tau_b$ and $\sigma_{ab}$) averaged for each ring:ring stacking contact (i.e. average parameters from 5||5, 5||6 and 6||6 ring:ring contacts) for each example stack are provided in parentheses respectively

| | | α face (56), (6) | |
|---|---|---|---|
| | | A | G |
| α face (5), (6) | A | | G1806(X)\|\|A1807(X); 4io9 (4.2 Å, 14.2°, 25.8°, 77.7°) |
| | | | A1050(1H)\|\|G2751(1H);5e7k (3.5 Å, 12.2°, 26.7°, 21.2°, 75.6°) |
| | G | | |

.

**Table S9**. Examples (i.e. identities of bases involved in stacking and corresponding PDB codes) of consecutive (black) and non-consecutive (grey) 5||56, 6||5 α||α *cis* purine||purine stacks. The geometrical parameters ($\vec{d}_{ab}$, $\theta_{ab}$, $\tau_a$, $\tau_b$ and $\sigma_{ab}$) averaged for each ring:ring stacking contact (i.e. average parameters from 5||5, 5||6 and 6||5 ring:ring contacts) for each example stack are provided in parentheses respectively.

<table>
<tr><td colspan="2" rowspan="2"></td><td colspan="2">α face (56), (6)</td></tr>
<tr><td>A</td><td>G</td></tr>
<tr><td rowspan="4">α face (5), (6)</td><td rowspan="2">A</td><td>A806(1)||A807(1);5fci<br>(4.0 Å, 22.4°, 14.4°, 24.2°,1.9°)</td><td>A2841(0)||G2842(0);3i55<br>(4.1 Å, 20.4°, 25.0°, 35.4°,43.1°)</td></tr>
<tr><td>A1492(AA)||A1913(BB); 4v50<br>(3.7 Å, 21.4°, 16.2°, 30.9°, 60.0°)</td><td></td></tr>
<tr><td rowspan="2">G</td><td rowspan="2"></td><td>G1647(1G)||G1648(1G);5ndk<br>(4.0 Å, 22.2°, 21.8°, 13.0°, 30.7°)</td></tr>
<tr><td>G22(B)||G54(B);5ddq<br>(3.8 Å, 11.1°, 21.5°, 26.5°, 50.5°)</td></tr>
</table>

**Table S10**. Examples (i.e. identities of bases involved in stacking and corresponding PDB codes) of consecutive (black) and non-consecutive (grey) 5||6, 6||6 α||α *cis* purine||purine stacks. The geometrical parameters ($\vec{d}_{ab}$, $\theta_{ab}$, $\tau_a$, $\tau_b$ and $\sigma_{ab}$) averaged for each ring:ring stacking contact (i.e. average parameters from 5||6 and 6||6 ring:ring contacts) for each example stack are provided in parentheses respectively.

<table>
<tr><td colspan="2" rowspan="2"></td><td colspan="2">α face (6), (6)</td></tr>
<tr><td>A</td><td>G</td></tr>
<tr><td rowspan="4">α face (5), (6)</td><td rowspan="2">A</td><td>A1433(DA)||A1434(DA); 4v7u (3.9 Å, 19.6°, 23.7°, 26.4°, 26.0°)</td><td>A7(C)||G8(C); 4z0c<br>(4.1 Å, 18.3°, 23.7°, 37.6°, 34.5°)</td></tr>
<tr><td>A115(1)||A265(1); 5fci<br>(3.7 Å, 10.8°, 27.7°, 19.4°,7.5° )</td><td>A909(C)||G945(C); 1il2<br>(4.1 Å, 19.4°, 14.4°, 22.8°, 2.0°)</td></tr>
<tr><td rowspan="2">G</td><td rowspan="2"></td><td>G176(BA)||G177(BA); 4v51<br>(4.1 Å, 22.4°, 17.8°, 29.1°, 9.7°)</td></tr>
<tr><td>G1417(AA)||G1482(AA); 4u24<br>(3.7 Å, 20.2°, 20.4°, 26.7°, 25.3°)</td></tr>
</table>

.

**Table S11.** Examples (i.e. identities of bases involved in stacking and corresponding PDB codes) of consecutive (black) and non-consecutive (grey) 6||56 α||α *cis* purine||purine stacks. The geometrical parameters ($\vec{d}_{ab}$, $\theta_{ab}$, $\tau_a$, $\tau_b$ and $\sigma_{ab}$) averaged for each ring:ring stacking contact (i.e. average parameters from 6||5 and 6||6 ring:ring contacts) for each example stack are provided in parentheses respectively.

| | | α face (56) | |
|---|---|---|---|
| | | A | G |
| α face (6) | A | | A1287(DA)\|\|G1288(DA); 4v7t (4.2 Å, 21.0°, 27.7°, 18.9°, 16.6°) |
| | | | A9(CV)\|\|G44(CV); 4v9c (4.0 Å, 15.5°, 15.4°, 15.2°, 23.4°) |
| | G | | |

**Table S12**. Examples (i.e. identities of bases involved in stacking and corresponding PDB codes) of consecutive (black) and non-consecutive (grey) 5||5, 6||6 α||α *cis* purine||purine stacks. The geometrical parameters ($\vec{d}_{ab}$, $\theta_{ab}$, $\tau_a$, $\tau_b$ and $\sigma_{ab}$) averaged for each ring:ring stacking contact (i.e. average parameters from 5||5 and 6||6 ring:ring contacts) for each example stack are provided in parentheses respectively

| | | α face (5), (6) | |
|---|---|---|---|
| | | A | G |
| α face (5), (6) | A | | G1814(GA)\|\|A1815(GA);4v9o (4.4 Å, 16.6°,24.5°,16.5°, 83.5°) |
| | G | | G1024(AA)\|\|G1025(AA); 4v9o (4.2 Å, 11.0°, 28.4°, 22.9°, 85.3°) |

**Table S13**. Examples (i.e. identities of bases involved in stacking and corresponding PDB codes) of consecutive (black) and non-consecutive (grey) 5||6, 6||5 α||α *cis* purine||purine stacks. The geometrical parameters ($\vec{d}_{ab}$, $\theta_{ab}$, $\tau_a$, $\tau_b$ and $\sigma_{ab}$) averaged for each ring:ring stacking contact (i.e. average parameters from 6||5 and 6||5 ring:ring contacts) for each example stack are provided in parentheses respectively

<table>
<tr><td rowspan="2" colspan="2"></td><td colspan="2">α face (6), (5)</td></tr>
<tr><td>A</td><td>G</td></tr>
<tr><td rowspan="4">α face (5) ,(6)</td><td rowspan="2">A</td><td>A1085(RA)||A1086(RA);4www<br>(3.7 Å, 21.0°, 20.1°, 33.8°,12.8°)</td><td>A427(x)||G428(x);4wfa<br>(3.9 Å, 22.1°, 17.8°, 17.5°,24.0°)</td></tr>
<tr><td>A556(x)||A1233(x);5dm7<br>(4.1 Å, 20.9°, 20.1°, 35.5°,27.1°)</td><td>G976 (FA)|| A1362 (FA);4v9o<br>(4.2 Å, 15.1°, 26.6°, 13.8°, 70.5°)</td></tr>
<tr><td rowspan="2">G</td><td rowspan="2"></td><td>G1356(x)||G1357(x);4wfb<br>(3.8 Å, 22.4°, 31.4°, 14.0°, 1.7°)</td></tr>
<tr><td>G2133(BA)||G2157(BA); 4v8i<br>(4.1 Å, 21.2°,10.8°, 18.4°,15.8°)</td></tr>
</table>

**Table S14**. Examples (i.e. identities of bases involved in stacking and corresponding PDB codes) of consecutive (black) and non-consecutive (grey) 5||56 α||α *cis* purine||purine stacks. The geometrical parameters ($\vec{d}_{ab}$, $\theta_{ab}$, $\tau_a$, $\tau_b$ and $\sigma_{ab}$) averaged for each ring:ring stacking contact (i.e. average parameters from 5||5 and 5||6 ring:ring contacts) for each example stack are provided in parentheses respectively.

<table>
<tr><td rowspan="2" colspan="2"></td><td colspan="2">α face (56)</td></tr>
<tr><td>A</td><td>G</td></tr>
<tr><td rowspan="4">α face (5)</td><td rowspan="2">A</td><td>A806(1)||A807(1); 4u4n<br>(4.0 Å, 22.8°,14.5°, 26.5°,1.0°)</td><td>A1260(5)||G1261(5); 4u55<br>(4.3 Å, 22.5°, 31.6°,16.1°,12.2°)</td></tr>
<tr><td>A1492(CA)||A1913(DA); 4v84<br>(4.1 Å, 12.3°, 19.8°, 25.7°,88.8°)</td><td>A1050(BA)||G2751(BA); 4v8b<br>(3.8 Å, 19.7°, 22.1°, 37.7°,77.0°)</td></tr>
<tr><td rowspan="2">G</td><td rowspan="2"></td><td>G1492(1)||G1493(1); 4u51<br>(4.3 Å, 22.1°, 26.7°, 13.7°,44.4°)</td></tr>
<tr><td>G22(B)||G4(B); 5ddr<br>(3.6 Å, 22.8°, 17.0°, 25.5°, 50.9°)</td></tr>
</table>

**Table S15.**Examples (i.e. identities of bases involved in stacking and corresponding PDB codes) of consecutive (black) and non-consecutive (grey) 5||5, 6||5 α||α *cis* purine||purine stacks. The geometrical parameters ($\vec{d}_{ab}$, $\theta_{ab}$, $\tau_a$, $\tau_b$ and $\sigma_{ab}$) averaged for each ring:ring stacking contact (i.e. average parameters from 6||5 and 6||6 ring:ring contacts) for each example stack are provided in parentheses respectively.

| | | α face (5), (5) | |
|---|---|---|---|
| | | A | G |
| α face (5), (6) | A | | A2841(0)\|\|G2842(0); 1vq5 (4.1 Å, 19.4°, 23.7°, 33.6°,46.1°) |
| | | | G976(AA)\|\|A1362(AA); 5j8a (4.1 Å, 22.3°, 24.3°,21.4°, 64.1°) |
| | G | | |

**Table S16**. Examples (i.e. identities of bases involved in stacking and corresponding PDB codes) of consecutive (black) and non-consecutive (grey) 6||6 α||α *cis* purine||purine stacks. The geometrical parameters ($\vec{d}_{ab}$, $\theta_{ab}$, $\tau_a$, $\tau_b$ and $\sigma_{ab}$) for each example stack are provided in parentheses respectively.

| | | α face (6) | |
|---|---|---|---|
| | | A | G |
| α face (6) | A | A3056(9)\|\|A3057(9);1vqp (3.8 Å, 3.9°, 22.1°, 18.2°, 1.8°) | G1348(da)\|\|A1349(da);4v5e (3.8 Å, 12.3°, 12.2°, 16.2°, 6.3°) |
| | | A173(a)\|\|A126(a);1u9s (3.4 Å, 10.9°, 19.4°, 23.3°,32.2°) | G8(e)\|\|A12(d);1m5p (3.8 Å, 6.0°, 23.3°, 28.9°, 60.8°) |
| | G | | G2781(x)\|\|G2782(x);3dll (4.4 Å, 1.0°, 29.5°, 28.5°, 13.1°) |
| | | | G32(d)\|\|G43(d);1e7k (4.2 Å, 16.0°, 14.4°, 25.4°, 46.6°) |

**Table S17**. Examples (i.e. identities of bases involved in stacking and corresponding PDB codes) of consecutive (black) and non-consecutive (grey) 5||6 α||α *cis* purine||purine stacks. The geometrical parameters ($\vec{d}_{ab}$, $\theta_{ab}$, $\tau_a$, $\tau_b$ and $\sigma_{ab}$) for each example stack are provided in parentheses respectively.

| | | α face (6) | |
|---|---|---|---|
| | | A | G |
| α face (5) | A | | A2841(0)\|\|G2842(0); 3ccj<br>(4.2 Å, 22.3°, 27.7°, 38.5°, 40.8°) |
| | | | A19(A)\|\|G35(A); 2ygh<br>(4.2 Å, 19.5°, 35.0°, 34.8°,64.3°) |
| | G | | |

**Table S18**. Examples (i.e. identities of bases involved in stacking and corresponding PDB codes) of consecutive (black) and non-consecutive (grey) 6||5 α||α *cis* purine||purine stacks. The geometrical parameters ($\vec{d}_{ab}$, $\theta_{ab}$, $\tau_a$, $\tau_b$ and $\sigma_{ab}$) for each example stack are provided in parentheses respectively.

| | | α face (5) | |
|---|---|---|---|
| | | A | G |
| α face (6) | A | A78(X)\|\|A79(X); 2gtt<br>(4.0 Å, 11.4°, 24.2°, 35.8°, 0.1°) | A1287(CA)\|\|G1288(CA); 4v9p<br>(3.6 Å, 22.8°, 10.5°, 23.9°,3.1°) |
| | | A644(14)\|\|A646(14); 5ib8<br>(3.5 Å, 23.0°, 33.4°,10.5°, 44.0°) | A9(A)\|\|G44(A); 3wfs<br>(4.0 Å, 18.5°, 20.3°, 7.9°, 58.3°) |
| | G | | G107(CA)\|\|G108(CA); 4v8b<br>(4.4 Å, 15.9°, 35.8°, 21.6°, 29.4°) |
| | | | G1347(A)\|\|G1373(A); 5wns<br>(4.0 Å, 7.0°, 23.5°, 16.5°, 75.2°) |

**Table S19**. Examples (i.e. identities of bases involved in stacking and corresponding PDB codes) of consecutive (black) and non-consecutive (grey) 5||5 α||α *cis* purine||purine stacks. The geometrical parameters ($\vec{d}_{ab}$, $\theta_{ab}$, $\tau_a$, $\tau_b$ and $\sigma_{ab}$) for each example stack are provided in parentheses respectively.

| | | α face (5) | |
|---|---|---|---|
| | | A | G |
| α face (5) | A | A1096(X)\|\|A1097(X);3cf5 (3.9 Å, 20.2°, 12.5°, 27.5°, 20.2°) | A2841(A)\|\|G2842(A); 1nji (3.8 Å, 21.6°, 13.6°, 30.9°, 46.8°) |
| | | | A411 (A)\|\| G413 (A);4ji0 (3.8 Å, 16.3°, 32.4°, 36.7°,17.2° ) |
| | G | | G25(C)\|\|G26(C);5de5 (4.1 Å, 12.7°, 36.9°, 29.6°, 58.6°) |
| | | | G9(A)\|\|G12(A);5de8 (3.9 Å, 11.3°, 18.9°, 29.9°, 71.5°) |

**Table S20**. Examples (i.e. identities of bases involved in stacking and corresponding PDB codes) of consecutive (black) and non-consecutive (grey) 5||56, 6||56 α||α *trans* purine||purine stacks. The geometrical parameters ($\vec{d}_{ab}$, $\theta_{ab}$, $\tau_a$, $\tau_b$ and $\sigma_{ab}$) averaged for each ring:ring stacking contact (i.e. average parameters from 5||5, 5||6, 6||5 and 6||6 ring:ring contacts) for each example stack are provided in parentheses respectively.

| | | α face (56), (56) | |
|---|---|---|---|
| | | A | G |
| α face (5), (6) | A | A972(BA)\|\|A973(BA);4v85 (4.2 Å, 20.3°, 24.3°, 22.1°, 90.9°) | A1070(0)\|\|G1071(0);1yj9 (4.1 Å, 13.5°, 22.0°, 24.6°,94.4°) |
| | | A1390(1)\|\|A1418(1);4u56 (3.8 Å, 8.8°, 27.0°, 23.1°, 175.5°) | G795 (1A)\|\| A797 (1A);5wis (3.8 Å, 13.7°,25.6°, 19.0°,137.6°) |
| | G | | G1035(X)\|\|G1036(X); 2zjr (3.9 Å, 3.9°, 30.1°, 30.7°, 95.7°) |
| | | | G1198(6)\|\|G1200(6);5dgv (3.9 Å, 8.6°, 27.0°, 27.7°, 111.5°) |

**Table S21**. Examples (i.e. identities of bases involved in stacking and corresponding PDB codes) of consecutive (black) and non-consecutive (grey) 5||6, 6||56 α||α *trans* purine||purine stacks. The geometrical parameters ($\vec{d}_{ab}$, $\theta_{ab}$, $\tau_a$, $\tau_b$ and $\sigma_{ab}$) averaged for each ring:ring stacking contact (i.e. average parameters from 5||6, 6||5 and 6||6 ring:ring contacts) for each example stack are provided in parentheses respectively.

| | | α face (6), (56) | |
|---|---|---|---|
| | | A | G |
| α face (5), (6) | A | | |
| | | A2078(X)\|\|A2641(X);4wfa (4.0, Å, 4.0°, 26.7°, 29.5°, 141.5°) | A21(1x)\|\|G46(1x); 5hcq (3.9 Å, 6.7°, 26.1°, 24.3°, 138.2°) |
| | G | | |
| | | | G2437(0)\|\|G2469(0);2ogm (3.6 Å, 11.3°,26.5°, 22.2°, 115.1°) |

**Table S22**. Examples (i.e. identities of bases involved in stacking and corresponding PDB codes) of consecutive (black) and non-consecutive (grey) 5||5, 6||56 α||α *trans* purine||purine stacks. The geometrical parameters ($\vec{d}_{ab}$, $\theta_{ab}$, $\tau_a$, $\tau_b$ and $\sigma_{ab}$) averaged for each ring:ring stacking contact (i.e. average parameters from 5||5, 6||5 and 6||6 ring:ring contacts) for each example stack are provided in parentheses respectively.

| | | α face (5), (56) | |
|---|---|---|---|
| | | A | G |
| α face (5), (6) | A | A1775(0)\|\|A1776(0); 3ccu (4.1 Å, 17.9°, 26.8°, 27.6°, 90.9°) | G75(3)\|\|A76(3); 5dgf (4.0 Å, 19.5°, 17.5°, 21.0°, 91.9°) |
| | | A713(X)\|\|A715(X); 5nrg (3.5 Å, 4.4°, 16.5°, 18.0°, 158.1°) | G56(0)\|\|A59(0); 3ccs (3.9 Å, 8.7°, 28.1°, 28.3°, 126.7°) |
| | G | | G1024(EA)\|\|G1025(EA); 4v9o (3.9 Å, 18.0°, 26.9°, 19.2°, 98.9°) |
| | | | G1198(6)\|\|1200(6); 4u50 (3.8 Å, 17.1°, 21.3°, 24.0°, 116.0°) |

.

**Table S23**. Examples (i.e. identities of bases involved in stacking and corresponding PDB codes) of consecutive (black) and non-consecutive (grey) 5||56, 6||6 α||α *trans* purine||purine stacks. The geometrical parameters ($\vec{d}_{ab}$, $\theta_{ab}$, $\tau_a$, $\tau_b$ and $\sigma_{ab}$) averaged for each ring:ring stacking contact (i.e. average parameters from 5||5, 5||6 and 6||6 ring:ring contacts) for each example stack are provided in parentheses respectively.

| | | α face (56), (6) | |
|---|---|---|---|
| | | A | G |
| α face (5), (6) | A | | G2308(BA)\|\|A2309(BA); 4v7x (3.9 Å, 18.1°, 22.7°, 13.9°, 119.8°) |
| | | | A21(C)\|\|G46(C); 5wwr (3.6 Å, 7.2°, 20.9°, 25.3°, 164.1°) |
| | G | | |

.

.

**Table S24**. Examples (i.e. identities of bases involved in stacking and corresponding PDB codes) of consecutive (black) and non-consecutive (grey) 5||56, 6||5 α||α *trans* purine||purine stacks. The geometrical parameters ($\vec{d}_{ab}$, $\theta_{ab}$, $\tau_a$, $\tau_b$ and $\sigma_{ab}$) averaged for each ring:ring stacking contact (i.e. average parameters from 5||5, 5||6 and 6||5 ring:ring contacts) for each example stack are provided in parentheses respectively.

| | | α face (56), (6) | |
|---|---|---|---|
| | | A | G |
| α face (5), (6) | A | | |
| | | | A21(IB)\|\|G46(IB); 5j4d (4.2 Å, 13.1°, 19.4°, 17.6°, 97.3°) |
| | G | | |
| | | | G1347(ca)\|\|G1373(ca);4v8h (3.9 Å, 19.6°, 26.8°, 17.9°, 96.9°) |

.

**Table S25**. Examples (i.e. identities of bases involved in stacking and corresponding PDB codes) of consecutive (black) and non-consecutive (grey) 5||6, 6||6 α||α *trans* purine||purine stacks. The geometrical parameters ($\vec{d}_{ab}$, $\theta_{ab}$, $\tau_a$, $\tau_b$ and $\sigma_{ab}$) averaged for each ring:ring stacking contact (i.e. average parameters from 5||6 and 6||6 ring:ring contacts) for each example stack are provided in parentheses respectively.

| | | α face (6), (6) | |
|---|---|---|---|
| | | A | G |
| α face (5), (6) | A | A2034(X)\|\|A2593(X); 3pip (3.5 Å, 8.4°, 24.4°,25.5°, 130.0°) | A66(BB)\|\|G107(BB); 4u24 (3.7 Å, 4.1°, 20.2°,21.3 °,119.3°) |
| | G | | G10(A)\|\|G19(A); 2zy6 (4.0 Å, 10.8°, 25.9°, 24.6°, 133.7°) |

**Table S26**. Examples (i.e. identities of bases involved in stacking and corresponding PDB codes) of consecutive (black) and non-consecutive (grey) 6||56 α||α *trans* purine||purine stacks. The geometrical parameters ($\vec{d}_{ab}$, $\theta_{ab}$, $\tau_a$, $\tau_b$ and $\sigma_{ab}$) averaged for each ring:ring stacking contact (i.e. average parameters from 6||5 and 6||6 ring:ring contacts) for each example stack are provided in parentheses respectively.

| | | α face (56) | |
|---|---|---|---|
| | | A | G |
| α face (6) | A | | A34(DB)\|\|G44(DB); 4u27 (3.5 Å, 10.1°, 20.1°, 27.5°, 177.8°) |
| | G | | |

.

**Table S27**. Examples (i.e. identities of bases involved in stacking and corresponding PDB codes) of consecutive (black) and non-consecutive (grey) 5||5, 6||6 α||α *trans* purine||purine stacks. The geometrical parameters ($\vec{d}_{ab}$, $\theta_{ab}$, $\tau_a$, $\tau_b$ and $\sigma_{ab}$) averaged for each ring:ring stacking contact (i.e. average parameters from 5||5 and 6||6 ring:ring contacts) for each example stack are provided in parentheses respectively.

| | | α face (5), (6) | |
|---|---|---|---|
| | | A | G |
| α face (5), (6) | A | | G1019(14)‖A1020(14);5ndk (4.3 Å, 11.8°, 19.3°, 21.3°, 90.5°) |
| | | A246(ca)‖A279(ca);1vy4 (3.8 Å, 1.5°, 30.9°, 31.6°, 173.7°) | G761 (0)‖ A763 (0);2ogm (3.7 Å, 16.0°, 25.7°, 26.4°, 143.0°) |
| | G | | G45 (ya)‖ G215 (ya);4www (4.3 Å, 4.1°, 13.0°, 13.0°, 148.2°) |

**Table S28**. Examples (i.e. identities of bases involved in stacking and corresponding PDB codes) of consecutive (black) and non-consecutive (grey) 5||6, 6||5 α||α *trans* purine||purine stacks. The geometrical parameters ($\vec{d}_{ab}$, $\theta_{ab}$, $\tau_a$, $\tau_b$ and $\sigma_{ab}$) averaged for each ring:ring stacking contact (i.e. average parameters from 5||6 and 6||5 ring:ring contacts) for each example stack are provided in parentheses respectively.

| | | α face (6), (5) | |
|---|---|---|---|
| | | A | G |
| α face (5), (6) | A | | |
| | G | | |

**Table S29**. Examples (i.e. identities of bases involved in stacking and corresponding PDB codes) of consecutive (black) and non-consecutive (grey) 5||56 α||α *trans* purine||purine stacks. The geometrical parameters ($\vec{d}_{ab}$, $\theta_{ab}$, $\tau_a$, $\tau_b$ and $\sigma_{ab}$) averaged for each ring:ring stacking contact (i.e. average parameters from 5||5 and 5||6 ring:ring contacts) for each example stack are provided in parentheses respectively.

| | | α face (56) | |
|---|---|---|---|
| | | A | G |
| α face (5) | A | A1069 (2A)\|\| A1073 (2A); 6fkr (4.2 Å, 21.9°, 14.7°, 20.1°, 97.9°) | G60 (A)\|\| A61 (A); 4nya (3.7 Å, 16.6°, 22.4°, 20.0°, 98.2°) |
| | | | A589(5)\|\|G610(5); 4u55 (3.4 Å, 11.7°, 18.4°, 22.9°, 167.1°) |
| | G | | |
| | | | G447(CA)\|\|G485(CA); 4v9k (4.3 Å, 11.0°, 23.1°, 15.4°, 108.4°) |

**Table S30**. Examples (i.e. identities of bases involved in stacking and corresponding PDB codes) of consecutive (black) and non-consecutive (grey) 5||5, 6||5 α||α *trans* purine||purine stacks. The geometrical parameters ($\vec{d}_{ab}$, $\theta_{ab}$, $\tau_a$, $\tau_b$ and $\sigma_{ab}$) averaged for each ring:ring stacking contact (i.e. average parameters from 5||5 and 6||5 ring:ring contacts) for each example stack are provided in parentheses respectively.

| | | α face (5), (5) | |
|---|---|---|---|
| | | A | G |
| α face (5), (6) | A | | G90(X)\|\|A91 (X); 5dm6 (4.0 Å, 19.0°, 25.4°, 12.5°, 98.6°) |
| | | | A34(GB)\|\|G44(GB); 4v9p (3.9 Å, 11.8°, 26.8°, 26.0°, 153.8°) |
| | G | | |

**Table S31**. Examples (i.e. identities of bases involved in stacking and corresponding PDB codes) of consecutive (black) and non-consecutive (grey) 6||6 α||α *trans* purine||purine stacks. The geometrical parameters ($\vec{d}_{ab}$, $\theta_{ab}$, $\tau_a$, $\tau_b$ and $\sigma_{ab}$) for each example stack are provided in parentheses respectively.

<table>
<tr><td colspan="2" rowspan="2"></td><td colspan="2">α face (6)</td></tr>
<tr><td>A</td><td>G</td></tr>
<tr><td rowspan="4">α face (6)</td><td rowspan="2">A</td><td>A2749 (BA)|| A2750 (BA);4v9l<br>(4.4 Å, 19.8°, 23.9°,5.7°, 92.7°)</td><td></td></tr>
<tr><td>A20(A)||A33(A);4kqy<br>(4.2 Å, 4.8°, 33.1°, 33.5°,142.1°)</td><td>G720 (2A)|| A850 (2A);4z8c<br>(3.7 Å, 12.7°, 20.9°, 11.0°, 110.7°)</td></tr>
<tr><td rowspan="2">G</td><td rowspan="2"></td><td></td></tr>
<tr><td>G1300(CA)||G1334(CA);1vy6<br>(3.7 Å, 14.4°, 25.2°, 21.0°, 150.0°)</td></tr>
</table>

**Table S32**. Examples (i.e. identities of bases involved in stacking and corresponding PDB codes) of consecutive (black) and non-consecutive (grey) 5||6 α||α *trans* purine||purine stacks. The geometrical parameters ($\vec{d}_{ab}$, $\theta_{ab}$, $\tau_a$, $\tau_b$ and $\sigma_{ab}$) for each example stack are provided in parentheses respectively.

<table>
<tr><td colspan="2" rowspan="2"></td><td colspan="2">α face (6)</td></tr>
<tr><td>A</td><td>G</td></tr>
<tr><td rowspan="3">α face (5)</td><td rowspan="2">A</td><td rowspan="2"></td><td></td></tr>
<tr><td>A1028(DA)||G1125(DA); 4u27<br>(4.5 Å, 3.2°, 32.7°, 33.4°, 113.9°)</td></tr>
<tr><td>G</td><td></td><td></td></tr>
</table>

**Table S33**. Examples (i.e. identities of bases involved in stacking and corresponding PDB codes) of consecutive (black) and non-consecutive (grey) 6||5 α||α *trans* purine||purine stacks. The geometrical parameters ($\vec{d}_{ab}$, $\theta_{ab}$, $\tau_a$, $\tau_b$ and $\sigma_{ab}$) for each example stack are provided in parentheses respectively.

| | | α face (5) | |
|---|---|---|---|
| | | A | G |
| α face (6) | A | | |
| | | A668(DA)\|\|A670(DA); 4u24 (3.1 Å, 15.4°, 8.0°, 13.7°, 162.3°) | A22(CD)\|\|G47(CD); 4v8b (3.3 Å, 16.9°, 28.2°, 15.1°, 105.0°) |
| | G | | |
| | | | G372(BA)\|\|G400(BA); 4u24 (3.9 Å, 7.1°, 26.4°, 27.7°, 106.6°) |

**Table S34**. Examples (i.e. identities of bases involved in stacking and corresponding PDB codes) of consecutive (black) and non-consecutive (grey) 5||5 α||α *trans* purine||purine stacks. The geometrical parameters ($\vec{d}_{ab}$, $\theta_{ab}$, $\tau_a$, $\tau_b$ and $\sigma_{ab}$) for each example stack are provided in parentheses respectively.

| | | α face (5) | |
|---|---|---|---|
| | | A | G |
| α face (5) | A | A203(da)\|\|A204 (da);4u24 (3.8 Å, 11.1°, 18.7°, 7.7°, 90.1°) | A889 (CA)\|\|G888 (CA); 4v67 (4.0 Å, 12.2°, 11.0°, 19.9°, 93.0°) |
| | | A681(0)\|\|A683(0);1ond (3.5 Å, 22.6°, 18.9°, 35.8°, 165.4°) | G2307(1H)\|\|A2311(1H);4wzd (4.1 Å, 15.2°, 32.5°, 18.0°, 116.0°) |
| | G | | G1035(X)\|\|G1036(X);2zjq (3.3 Å, 5.2°, 17.6°, 19.7°, 95.9°) |
| | | | G45 (da)\|\| G215 (da);4u27 (4.4 Å, 16.6°, 15.3°, 4.1°, 157.5°) |

**Table S35**. Examples (i.e. identities of bases involved in stacking and corresponding PDB codes) of consecutive (black) and non-consecutive (grey) 5||56, 6||56 α||β *cis* purine||purine stacks. The geometrical parameters ($\vec{d}_{ab}$, $\theta_{ab}$, $\tau_a$, $\tau_b$ and $\sigma_{ab}$) averaged for each ring:ring stacking contact (i.e. average parameters from 5||5, 5||6, 6||5 and 6||6 ring:ring contacts) for each example stack are provided in parentheses respectively.

| | | β face (56), (56) | |
|---|---|---|---|
| | | A | G |
| α face (5), (6) | A | A511(X)\|\|A512(X);3pip<br>(3.9 Å, 22.0°, 25.6°, 19.6°34.2°) | A5(A)\|\|G6(A);4lvv<br>(3.9 Å, 19.8°, 19.6°, 23.4°, 47.5°) |
| | | A716(5)\|\|A720(5); 4u3m<br>(3.8 Å, 2.9°, 25.3°, 24.4°, 70.1°) | A225(x)\|\|G227(x);4wf9<br>(3.9 Å, 10.2°, 26.1°, 28.2°, 58.0°) |
| | G | | G557(A)\|\|G5581(A);1xnr<br>(4.0 Å, 18.4°, 24.8°, 19.4°, 38.9°) |
| | | | G718(1h)\|\|G850(1h); 5ndk<br>(3.8 Å, 16.5°, 21.6°, 24.4°, 49.4°) |

**Table S36**. Examples (i.e. identities of bases involved in stacking and corresponding PDB codes) of consecutive (black) and non-consecutive (grey) 5||6, 6||56 α||β *cis* purine||purine stacks. The geometrical parameters ($\vec{d}_{ab}$, $\theta_{ab}$, $\tau_a$, $\tau_b$ and $\sigma_{ab}$) averaged for each ring:ring stacking contact (i.e. average parameters from 5||6, 6||5 and 6||6 ring:ring contacts) for each example stack are provided in parentheses respectively.

| | | β face (6), (56) | |
|---|---|---|---|
| | | A | G |
| α face (5), (6) | A | A45(C)\|\|A46(C); 3v7e<br>(4.0 Å, 21.7°, 22.3°, 26.7°, 38.8°) | A495(0)\|\| G496(0); 1yit<br>(4.1 Å, 22.7°, 25.1°, 36.9°) |
| | | A33(A)\|\|A35(A); 2qwy<br>(3.7 Å, 13.2°, 22.1°, 25.3°, 78.7°) | A120(AA)\|\|G122(AA);4v50<br>(4.0 Å, 21.8°, 20.8°, 27.7°, 78.5°) |
| | G | | G1706(0)\|\|G1707(0); 1yj9<br>(3.9 Å, 22.3°, 27.8°, 20.9°, 33.1°) |

**Table S37**. Examples (i.e. identities of bases involved in stacking and corresponding PDB codes) of consecutive (black) and non-consecutive (grey) 5||5, 6||56 α||β *cis* purine||purine stacks. The geometrical parameters ($\vec{d}_{ab}$, $\theta_{ab}$, $\tau_a$, $\tau_b$ and $\sigma_{ab}$) averaged for each ring:ring stacking contact (i.e. average parameters from 5||5, 5||6, 6||5 and 6||6 ring:ring contacts) for each example stack are provided in parentheses respectively.

| | | β face (5),(56) | |
|---|---|---|---|
| | | A | G |
| 9α face (5), (6) | A | A31(A)\|\|A32(A); 3vrs (3.7 Å,14.0°, 18.9°, 18.1°, 44.8°) | A41(B)\|\|G42(B); 2dr2 (3.9 Å, 12.9°, 18.2°, 23.6°, 19.0°) |
| | | A327(CA)\|\|A329(CA); 4u24 (3.4 Å, 10.1°, 21.3°, 19.1°, 32.1°) | A503(BA)\|\|G506(BA); 4u24 (3.8 Å, 13.0°, 25.1°, 19.5°, 60.7°) |
| | G | | G2(A)\|\|G3(A); 3vrs (3.8 Å, 3.0°, 27.8°, 27.2°, 25.7°) |
| | | | G1731(BA)\|\|G1733(BA); 4u24 (3.2 Å, 6.5°, 22.9°, 23.7°, 12.5°) |

**Table S38**. Examples (i.e. identities of bases involved in stacking and corresponding PDB codes) of consecutive (black) and non-consecutive (grey) 5||56, 6||6 α||β *cis* purine||purine stacks. The geometrical parameters ($\vec{d}_{ab}$, $\theta_{ab}$, $\tau_a$, $\tau_b$ and $\sigma_{ab}$) averaged for each ring:ring stacking contact (i.e. average parameters from 5||5, 5||6 and 6||6 ring:ring contacts) for each example stack are provided in parentheses respectively.

| | | β face (56), (6) | |
|---|---|---|---|
| | | A | G |
| α face (5), (6) | A | | A1632(DA)\|\|G1633(DA); 4u24 (4.0 Å, 22.9°, 20.5°, 21.5°, 35.9°) |
| | | | A404(BA)\|\|G406(BA); 4u24 (3.6 Å, 15.4°, 21.9°, 22.0°, 75.8° ) |
| | G | | |

**Table S39**. Examples (i.e. identities of bases involved in stacking and corresponding PDB codes) of consecutive (black) and non-consecutive (grey) 5||56, 6||5 α||β *cis* purine||purine stacks. The geometrical parameters ($\vec{d}_{ab}$, $\theta_{ab}$, $\tau_a$, $\tau_b$ and $\sigma_{ab}$) averaged for each ring:ring stacking contact (i.e. average parameters from 5||5, 5||6 and 6||5 ring:ring contacts) for each example stack are provided in parentheses respectively.

| | | β face (56), (5) | |
|---|---|---|---|
| | | A | G |
| α face (5), (6) | A | A1268(14)\|\|A1269(14);5ib8 (3.0 Å, 3.3°, 27.7°, 25.8°, 64.0°) | A583(1G)\|\|G584(1G);4wt1 (4.1 Å, 5.3°, 28.9°, 25.2°, 63.8°) |
| | | | A222(1H)\|\|A224(1H);5ib8 (4.1 Å, 5.1°, 29.6°, 29.4°, 56.3°) |
| | G | | G1142(A)\|\|G1143(A);4b3m (4.3 Å, 4.6°, 27.2°, 27.1°, 66.3°) |
| | | | G5(R)\|\|G7(R); 3pu1 (3.8 Å, 7.6°, 26.0°, 27.2°, 89.8°) |

**Table S40**. Examples (i.e. identities of bases involved in stacking and corresponding PDB codes) of consecutive (black) and non-consecutive (grey) 5||6, 6||6 α||β *cis* purine||purine stacks. The geometrical parameters ($\vec{d}_{ab}$, $\theta_{ab}$, $\tau_a$, $\tau_b$ and $\sigma_{ab}$) averaged for each ring:ring stacking contact (i.e. average parameters from 5||6 and 6||6 ring:ring contacts) for each example stack are provided in parentheses respectively.

| | | β face (6), (6) | |
|---|---|---|---|
| | | A | G |
| α face (5), (6) | A | A509(A)\|\|A510(A); 5wns (4.0 Å, 22.3°, 23.8°, 19.4°, 35.2°) | A1433(5)\|\|G1434(5); 4u51 (3.9 Å, 18.2°, 16.0°, 29.9°, 35.1°) |
| | | A15(DB)\|\|A109(DB); 4u24 (3.8 Å, 7.7°, 18.5°, 24.1°, 71.3°) | A9(1K)\|\|G46(1K); 6gsj (4.2 Å, 16.3°, 24.4°, 16.7°, 5.9°) |
| | G | | G1181(CA)\|\|G1182(CA); 4v64 (4.0 Å, 19.0°, 37.5°, 24.1°, 65.9°) |
| | | | G974(BB)\|\|G1186(BB); 4v64 (3.7 Å, 6.3°, 22.5°, 26.6°, 42.7°) |

**Table S41**. Examples (i.e. identities of bases involved in stacking and corresponding PDB codes) of consecutive (black) and non-consecutive (grey) 6||56 α||β *cis* purine||purine stacks. The geometrical parameters ($\vec{d}_{ab}$, $\theta_{ab}$, $\tau_a$, $\tau_b$ and $\sigma_{ab}$) averaged for each ring:ring stacking contact (i.e. average parameters from 6||5 and 6||6 ring:ring contacts) for each example stack are provided in parentheses respectively.

| | | β face (56) | |
|---|---|---|---|
| | | A | G |
| α face (6) | A | | A9(A)\|\|G10(A); 3vrs (3.9 Å, 9.1°, 26.2°, 30.9°, 20.3°) |
| | | | A503(DA)\|\|G506(DA); 4u24 (3.2 Å, 8.2°, 25.8°, 27.0°, 63.6°) |
| | G | | |

**Table S42**. Examples (i.e. identities of bases involved in stacking and corresponding PDB codes) of consecutive (black) and non-consecutive (grey) 5||5, 6||6 α||β *cis* purine||purine stacks. The geometrical parameters ($\vec{d}_{ab}$, $\theta_{ab}$, $\tau_a$, $\tau_b$ and $\sigma_{ab}$) averaged for each ring:ring stacking contact (i.e. average parameters from 5||5 and 6||6 ring:ring contacts) for each example stack are provided in parentheses respectively.

| | | β face (5), (6) | |
|---|---|---|---|
| | | A | G |
| α face (5), (6) | A | A2014(BA)\|\|A2015(BA); 4u24 (4.1 Å, 16.6°, 20.4°, 30.6°, 18.7°) | A1042(CA)\|\|A1043(CA); 4u24 (4.0 Å, 14.3°, 19.5°, 5.4°, 0.1°) |
| | | A484(0)\|\|A486(0);1ffk (4.1 Å, 22.6°, 28.9°, 30.5°, 50.6°) | A890(GB)\|\|G892(GB);6boh (4.5 Å, 20.6°, 19.8°, 25.3°, 59.8°) |
| | G | | G2168(d1)\|\|G2169(d1);4wt8 (4.3 Å, 5.8°, 35.5°, 34.5°, 19.8°) |
| | | | G945(CA)\|\|G1337 (CA);4u24 (5\|\|5)(4.2 Å, 9.4°, 28.7°, 37.0°, 5.7°) |

**Table S43**. Examples (i.e. identities of bases involved in stacking and corresponding PDB codes) of consecutive (black) and non-consecutive (grey) 5||6, 6||5 α||β *cis* purine||purine stacks. The geometrical parameters ($\vec{d}_{ab}$, $\theta_{ab}$, $\tau_a$, $\tau_b$ and $\sigma_{ab}$) averaged for each ring:ring stacking contact (i.e. average parameters from 5||6 and 6||5 ring:ring contacts) for each example stack are provided in parentheses respectively.

| | | β face (6), (5) | |
|---|---|---|---|
| | | A | G |
| α face (5), (6) | A | | |
| | G | | |

**Table S44**. Examples (i.e. identities of bases involved in stacking and corresponding PDB codes) of consecutive (black) and non-consecutive (grey) 5||56 α||β *cis* purine||purine stacks. The geometrical parameters ($\vec{d}_{ab}$, $\theta_{ab}$, $\tau_a$, $\tau_b$ and $\sigma_{ab}$) averaged for each ring:ring stacking contact (i.e. average parameters from 5||5 and 5||6 ring:ring contacts) for each example stack are provided in parentheses respectively.

| | | β face (56) | |
|---|---|---|---|
| | | A | G |
| α face (5) | A | A64(A)\|\|A65(A);2b57<br>(4.1 Å, 20.9°, 19.9°, 37.3°, 13.2°) | A14(CW)\|\|G15(CW); 4v8n<br>(4.0 Å, 21.3°, 21.3°, 18.7°, 50.5°) |
| | | A192(0)\|\|A194(0); 3ccu<br>(3.8 Å, 8.7°, 26.2°, 26.3°,55.5°) | A13(Bx)\|\|G963(Ba); 4v7j<br>(3.8 Å, 15.7°, 30.4°, 27.8°,37.7°) |
| | G | | G1138(BB)\|\|G1139(BB); 4v64<br>(4.3 Å, 21.7°, 22.2°, 33.1°, 69.4°) |
| | | | G1449(0)\|\|G1672(0); 3ccr<br>(4.1 Å, 7.4°, 30.8°, 25.2°, 6.2°) |

**Table S45**. Examples (i.e. identities of bases involved in stacking and corresponding PDB codes) of consecutive (black) and non-consecutive (grey) 5||5, 6||5 α||β *cis* purine||purine stacks The geometrical parameters ($\vec{d}_{ab}$, $\theta_{ab}$, $\tau_a$, $\tau_b$ and $\sigma_{ab}$) averaged for each ring:ring stacking contact (i.e. average parameters from 5||5 and 6||5 ring:ring contacts) for each example stack are provided in parentheses respectively.

| | | β face (5), (5) | |
|---|---|---|---|
| | | A | G |
| α face (5), (6) | A | | A20(C)\|\|G21(C); 2d2l (3.8 Å, 11.8°, 22.2°, 16.1°, 40.8°) |
| | | | A2799(DA)\|\|G2801(DA); 4u24 (3.5 Å, 6.9°, 36.4°, 33.3°, 52.0°) |
| | G | | |

**Table S46**. Examples (i.e. identities of bases involved in stacking and corresponding PDB codes) of consecutive (black) and non-consecutive (grey) 6||6 α||β *cis* purine||purine stacks. The geometrical parameters ($\vec{d}_{ab}$, $\theta_{ab}$, $\tau_a$, $\tau_b$ and $\sigma_{ab}$) for each example stack are provided in parentheses respectively.

| | | β face (6) | |
|---|---|---|---|
| | | A | G |
| α face (6) | A | A1693(0)\|\|A1694(0);2aar (3.6 Å, 21.6°, 31.2°, 11.8°, 33.2°) | A20(AX)\|\|G21(AX);4v5a (4.3 Å, 22.8°, 2.0°, 23.4°, 24.0°) |
| | | A484(0)\|\|A486(0);1yit (4.0 Å, 22.1°, 34.8°, 25.3°, 48.3°) | A1153(0)\|\|G1155(0);1y69 (3.7 Å, 8.6°, 38.3°, 29.7°, 53.6°) |
| | G | | G816(0)\|\|G817(0);1ffk (3.5 Å, 22.2°, 19.9°, 21.3°, 36.9°) |
| | | | G12(J)\|\|G16(J);2y9h (3.9 Å, 8.6°, 32.8°, 39.2°,49.9°) |

**Table S47**. Examples (i.e. identities of bases involved in stacking and corresponding PDB codes) of consecutive (black) and non-consecutive (grey) 5||6 α||β *cis* purine||purine stacks. The geometrical parameters ($\vec{d}_{ab}$, $\theta_{ab}$, $\tau_a$, $\tau_b$ and $\sigma_{ab}$) for each example stack are provided in parentheses respectively.

| | | β face (6) | |
|---|---|---|---|
| | | A | G |
| α face (5) | A | | A1433(5)\|\|G1434(5); 4u55<br>(3.6 Å, 20.1°, 3.2°, 22.6°, 39.4°) |
| | | | A665(CA)\|\|G725(CA); 4u24<br>(4.1 Å, 8.7°, 37.5°, 31.1°, 36.4°) |
| | G | | |

**Table S48**. Examples (i.e. identities of bases involved in stacking and corresponding PDB codes) of consecutive (black) and non-consecutive (grey) 6||5 α||β *cis* purine||purine stacks. The geometrical parameters ($\vec{d}_{ab}$, $\theta_{ab}$, $\tau_a$, $\tau_b$ and $\sigma_{ab}$) for each example stack are provided in parentheses respectively.

| | | β face (5) | |
|---|---|---|---|
| | | A | G |
| α face (6) | A | A347(BA)\|\|A348(BA); 4u24<br>(3.7 Å, 16.5°, 20.3°, 24.4°, 15.2°) | A18(C)\|\|G19(C); 2d2l<br>(3.4 Å, 5.9°, 18.7°, 13.2°, 17.6°) |
| | | A51(CA)\|\|A116(CA); 4u24<br>(4.0 Å, 5.1°, 30.3°, 27.5°, 2.1°) | A2274(DA)\|\|G2276(DA); 4u24<br>(4.1 Å, 3.2°, 38.7°, 37.8°, 59.8°) |
| | G | | G67(B)\|\|G68(B); 2dr2<br>(3.5 Å, 15.8°, 23.7°, 39.3°, 8.8°) |
| | | | G7(B)\|\|G49(B); 2dr2<br>(3.9 Å, 12.9°, 33.1°, 26.2°, 9.1°) |

**Table S49**. Examples (i.e. identities of bases involved in stacking and corresponding PDB codes) of consecutive (black) and non-consecutive (grey) 5||5 α||β *cis* purine||purine stacks. The geometrical parameters ($\vec{d}_{ab}$, $\theta_{ab}$, $\tau_a$, $\tau_b$ and $\sigma_{ab}$) for each example stack are provided in parentheses respectively.

| | | β face (5) | |
|---|---|---|---|
| | | A | G |
| α face (5) | A | A71(6)\|\|A7(6);5fcj<br>(4.5 Å, 19.5°, 39.9°, 33.1°, 7.0°) | A8(A)\|\|G9(A);4lvw<br>(4.1 Å, 6.8°, 33.3°, 29.0°,64.3°) |
| | | A1921(2A)\|\|A1991(A);5hcq<br>(4.3 Å, 23.0°, 17.8°, 34.5°, 3.8°) | A109(B)\|\|G111(B);4y1o<br>(4.0 Å, 5.3°, 36.4°, 31.8°, 10.8°) |
| | G | | G1183(1A)\|\|G1184(1A);5fdu<br>(4.1 Å, 18.8°, 25.8°, 30.5°, 70.4°) |
| | | | G1133(1a)\|\|G1135(1a);5fdu<br>(3.8 Å, 7.8°, 33.2°, 27.5°, 42.7°) |

**Table S50**. Examples (i.e. identities of bases involved in stacking and corresponding PDB codes) of consecutive (black) and non-consecutive (grey) 5||56, 6||56 α||β *trans* purine||purine stacks. The geometrical parameters ($\vec{d}_{ab}$, $\theta_{ab}$, $\tau_a$, $\tau_b$ and $\sigma_{ab}$) averaged for each ring:ring stacking contact (i.e. average parameters from 5||5, 5||6, 6||5 and 6||6 ring:ring contacts) for each example stack are provided in parentheses respectively.

| | | β face (56), (56) | |
|---|---|---|---|
| | | A | G |
| α face (5), (6) | A | | |
| | | A236(0)\|\|A435(0);4v9f<br>(3.8 Å, 12.2°, 30.6°, 21.7°, 166.7°) | A2725(BA)\|\|G2727(BA);4v8a<br>(3.9 Å, 3.1°,28.4°, 26.4°, 108.9°) |
| | G | | G881(YA)\|\|G882(YA);6buw<br>(4.2 Å, 5.1°,23.1°, 21.9°, 100.0°) |
| | | | G1874(1)\|\|G2945(1);4u6f<br>(3.8 Å, 6.4°, 26.8°, 25.3°, 155.4°) |

**Table S51**. Examples (i.e. identities of bases involved in stacking and corresponding PDB codes) of consecutive (black) and non-consecutive (grey) 5||6, 6||56 α||β *trans* purine||purine stacks. The geometrical parameters ($\vec{d}_{ab}$, $\theta_{ab}$, $\tau_a$, $\tau_b$ and $\sigma_{ab}$) averaged for each ring:ring stacking contact (i.e. average parameters from 5||6, 6||5 and 6||6 ring:ring contacts) for each example stack are provided in parentheses respectively.

| | | β face (6), (56) | |
|---|---|---|---|
| | | A | G |
| α face (5), (6) | A | A9(B)\|\|A9(C); 2fcz (3.8 Å, 1.8°, 27.7°, 27.7°, 135.3°) | |
| | | A35(B)\|\|A37(B); 4meh (3.9 Å, 3.5°, 24.1°,22.3°, 110.8°) | A1069(DA)\|\|G1074(DA); 4u24 (3.9 Å, 21.9°, 18.9°, 18.8°, 158.6°) |
| | G | | |
| | | | G2874(1)\|\|2945(1); 5dge (4.0 Å, 19.3°, 26.1°, 22.2°, 153.7°) |

**Table S52**. Examples (i.e. identities of bases involved in stacking and corresponding PDB codes) of consecutive (black) and non-consecutive (grey) 5||5, 6||56 α||β *trans* purine||purine stacks. The geometrical parameters ($\vec{d}_{ab}$, $\theta_{ab}$, $\tau_a$, $\tau_b$ and $\sigma_{ab}$) averaged for each ring:ring stacking contact (i.e. average parameters from 5||5, 6||5 and 6||6 ring:ring contacts) for each example stack are provided in parentheses respectively.

| | | β face (5), (56) | |
|---|---|---|---|
| | | A | G |
| α face (5), (6) | A | | |
| | | A1069(BA)\|\|A1073(BA); 4v8b (3.9 Å, 21.3°, 22.4°, 30.0°, 108.3°) | A119(CA)\|\|G240(CA); 4v7s (4.1 Å, .3.3°, 28.8°, 30.6°,119.9°) |
| | G | | |
| | | | G1633(DA)\|\|G1635(DA); 4v5s (3.6 Å, 12.7°, 22.2°, 21.5°, 94.6°) |

**Table S53**. Examples (i.e. identities of bases involved in stacking and corresponding PDB codes) of consecutive (black) and non-consecutive (grey) 5||56, 6||6 α||β *trans* purine||purine stacks. The geometrical parameters ($\vec{d}_{ab}$, $\theta_{ab}$, $\tau_a$, $\tau_b$ and $\sigma_{ab}$) averaged for each ring:ring stacking contact (i.e. average parameters from 5||5, 5||6 and 6||6 ring:ring contacts) for each example stack are provided in parentheses respectively.

<table>
<tr><td colspan="2" rowspan="2"></td><td colspan="2">β face (56), (6)</td></tr>
<tr><td>A</td><td>G</td></tr>
<tr><td rowspan="3">α face<br>(5), (6)</td><td rowspan="2">A</td><td rowspan="2"></td><td></td></tr>
<tr><td>A2764(CA)||G2766(CA); 4wqf<br>(3.9 Å, 22.7°, 21.6°, 29.7°, 94.7°)</td></tr>
<tr><td>G</td><td></td><td></td></tr>
</table>

**Table S54**. Examples (i.e. identities of bases involved in stacking and corresponding PDB codes) of consecutive (black) and non-consecutive (grey) 5||56, 6||5 α||β *trans* purine||purine stacks. The geometrical parameters ($\vec{d}_{ab}$, $\theta_{ab}$, $\tau_a$, $\tau_b$ and $\sigma_{ab}$) averaged for each ring:ring stacking contact (i.e. average parameters from 5||5, 5||6 and 6||6 ring:ring contacts) for each example stack are provided in parentheses respectively.

<table>
<tr><td colspan="2" rowspan="2"></td><td colspan="2">β face (56), (5)</td></tr>
<tr><td>A</td><td>G</td></tr>
<tr><td rowspan="4">α face<br>(5), (6)</td><td rowspan="2">A</td><td colspan="2">A2134(DA)||A2135(DA); 1vy6<br>(3.7 Å, 17.9°, 22.6°, 23.4°, 91.9°)</td></tr>
<tr><td>A199 (1)|| A201 (1);5tga<br>(3.5 Å, 10.1°, 26.9°, 21.7°, 118.7°)</td><td>A2369(0)||G2371(0);3cc7<br>(3.5 Å, 9.5°, 23.6°, 21.3°, 135.2°)</td></tr>
<tr><td rowspan="2">G</td><td rowspan="2"></td><td></td></tr>
<tr><td>G595(1)||G609(1);5dat<br>(3.6 Å, 18.5°, 21.2°, 20.7°, 112.3°)</td></tr>
</table>

**Table S55**. Examples (i.e. identities of bases involved in stacking and corresponding PDB codes) of consecutive (black) and non-consecutive (grey) 5||6, 6||6 α||β *trans* purine||purine stacks. The geometrical parameters ($\vec{d}_{ab}$, $\theta_{ab}$, $\tau_a$, $\tau_b$ and $\sigma_{ab}$) averaged for each ring:ring stacking contact (i.e. average parameters from 5||6 and 6||6 ring:ring contacts) for each example stack are provided in parentheses respectively.

| | | β face (6), (6) | |
|---|---|---|---|
| | | A | G |
| α face (5), (6) | A | | |
| | | A71(DA)\|\|A73(DA); 4u24<br>(4.2 Å, 13.0°,29.9°, 36.2°, 110.4°) | A45(0)\|\|G147(0);1ffk<br>(3.7 Å, 8.3°, 28.3°, 33.1°,174.5°) |
| | G | | G1181(A)\|\|G1182(A); 4aqy<br>(4.1 Å, 20.3°, 15.9°, 28.4°, 90.5°) |
| | | | G1063(5)\|\|G1097(5); 4u55<br>(3.6 Å, 11.5°, 19.0°, 16.2°, 107.3°) |

**Table S56**. Examples (i.e. identities of bases involved in stacking and corresponding PDB codes) of consecutive (black) and non-consecutive (grey) 6||56 α||β *trans* purine||purine stacks. The geometrical parameters ($\vec{d}_{ab}$, $\theta_{ab}$, $\tau_a$, $\tau_b$ and $\sigma_{ab}$) averaged for each ring:ring stacking contact (i.e. average parameters from 6||5 and 6||6 ring:ring contacts) for each example stack are provided in parentheses respectively.

| | | β face (56) | |
|---|---|---|---|
| | | A | G |
| α face (6) | A | | |
| | | | A1069(BA)\|\|G1074(BA); 4u24<br>(3.9 Å, 18.8°, 36.6°, 25.5°, 155.6°) |
| | G | | |

**Table S57**. Examples (i.e. identities of bases involved in stacking and corresponding PDB codes) of consecutive (black) and non-consecutive (grey) 5||5, 6||6 α||β *trans* purine||purine stacks. The geometrical parameters ($\vec{d}_{ab}$, $\theta_{ab}$, $\tau_a$, $\tau_b$ and $\sigma_{ab}$) averaged for each ring:ring stacking contact (i.e. average parameters from 5||5 and 6||6 ring:ring contacts) for each example stack are provided in parentheses respectively.

| | | β face (5), (6) | |
|---|---|---|---|
| | | A | G |
| α face (5), (6) | A | | |
| | G | | |

**Table S58**. Examples (i.e. identities of bases involved in stacking and corresponding PDB codes) of consecutive (black) and non-consecutive (grey) 5||6, 6||5 α||β *trans* purine||purine stacks. The The geometrical parameters ($\vec{d}_{ab}$, $\theta_{ab}$, $\tau_a$, $\tau_b$ and $\sigma_{ab}$) averaged for each ring:ring stacking contact (i.e. average parameters from 5||6 and 6||5 ring:ring contacts) for each example stack are provided in parentheses respectively.

| | | β face (6), (5) | |
|---|---|---|---|
| | | A | G |
| α face (5), (6) | A | | |
| | | A236(0)\|\|A435(0);1w2b<br>(3.6 Å, 22.1°, 31.1°, 11.7°, 161.9°) | A451(CA)\|\|G481(CA);4v8f<br>(4.0 Å, 4.5°, 38.6°, 37.3°, 167.3°) |
| | G | | |
| | | | G2505(BA)\|\|G2576(BA); 4u24<br>(3.7 Å, 13.0°, 30.4°, 26.3°, 155.9°) |

**Table S59**. Examples (i.e. identities of bases involved in stacking and corresponding PDB codes) of consecutive (black) and non-consecutive (grey) 5||56 α||β *trans* purine||purine stacks. The geometrical parameters ($\vec{d}_{ab}$, $\theta_{ab}$, $\tau_a$, $\tau_b$ and $\sigma_{ab}$) averaged for each ring:ring stacking contact (i.e. average parameters from 5||5 and 5||6 ring:ring contacts) for each example stack are provided in parentheses respectively.

| | | β face (56) | |
|---|---|---|---|
| | | A | G |
| α face (5) | A | A2134(CA)\|\|A2135(CA); 4wqy<br>(3.7 Å, 22.7°, 16.4°,23.5°, 93.6°) | A1067(AA)\|\|G1068(AA); 4v87<br>(4.2 Å, 22.9°, 17.8°, 19.5°, 99.0°) |
| | | A147(0)\|\|A1409(0);1y69<br>(3.4 Å, 14.8°, 18.3°, 26.1°,139.4°) | A2469(14)\|\|G2482(14); 5el5<br>(3.6 Å, 19.1°, 31.1°, 18.0°, 141.6°) |
| | G | | |
| | | | G1906(YA)\|\|1929(YA);6bz6<br>(4.0 Å, 4.0°, 24.7°, 26.5° , 98.8°) |

**Table S60**. Examples (i.e. identities of bases involved in stacking and corresponding PDB codes) of consecutive (black) and non-consecutive (grey) 5||5, 6||5 α||β *trans* purine||purine stacks. The geometrical parameters ($\vec{d}_{ab}$, $\theta_{ab}$, $\tau_a$, $\tau_b$ and $\sigma_{ab}$) averaged for each ring:ring stacking contact (i.e. average parameters from 5||5 and 6||5 ring:ring contacts) for each example stack are provided in parentheses respectively.

| | | β face (5), (5) | |
|---|---|---|---|
| | | A | G |
| α face (5), (6) | A | | A15(AW)\|\|G15(AW); 4c5c<br>(3.6 Å, 4.7°, 25.5°, 28.4°, 106.0°) |
| | | | A1553(DA)\|\|G1555(DA); 4u24<br>(3.6 Å, 19.9°, 29.9°, 18.5°, 121.1° ) |
| | G | | |

**Table S61**. Examples (i.e. identities of bases involved in stacking and corresponding PDB codes) of consecutive (black) and non-consecutive (grey) 6||6 α||β *trans* purine||purine stacks. The geometrical parameters ($\vec{d}_{ab}$, $\theta_{ab}$, $\tau_a$, $\tau_b$ and $\sigma_{ab}$) for each example stack are provided in parentheses respectively.

| | | β face (6) | |
|---|---|---|---|
| | | A | G |
| α face (6) | A | A13 (Bx)\|\| A14(Bx); 4v7j (4.5 Å, 8.2°, 17.2°, 25.4°, 94.9°) | |
| | | A792(A)\|\|A794(A);4jv5 (3.6 Å, 18.2°, 26.2°, 17.2°, 103.1°) | A35(A)\|\|G62(A);3g4m (3.8 Å, 5.2°, 32.0°, 26.9°, 148.5°) |
| | G | | G1(D)\|\|G2(C);1ze2 (4.0 Å, 6.7°, 36.5°, 30.0°, 140.9°) |
| | | | G15(c)\|\|G59(c);4rdx (3.9 Å, 7.7°, 29.2°, 21.6°, 174.3°) |

**Table S62**. Examples (i.e. identities of bases involved in stacking and corresponding PDB codes) of consecutive (black) and non-consecutive (grey) 5||6 α||β *trans* purine||purine stacks. The geometrical parameters ($\vec{d}_{ab}$, $\theta_{ab}$, $\tau_a$, $\tau_b$ and $\sigma_{ab}$) for each example stack are provided in parentheses respectively.

| | | β face (6) | |
|---|---|---|---|
| | | A | G |
| α face (5) | A | | |
| | | | A451(AA)\|\|G481(AA); 4u27 (4.1 Å, 21.7°, 39.9°, 26.4°, 174.0°) |
| | G | | |

**Table S63**. Examples (i.e. identities of bases involved in stacking and corresponding PDB codes) of consecutive (black) and non-consecutive (grey) 6||5 α||β *trans* purine||purine stacks. The geometrical parameters ($\vec{d}_{ab}$, $\theta_{ab}$, $\tau_a$, $\tau_b$ and $\sigma_{ab}$) for each example stack are provided in parentheses respectively.

| | | β face (6) | |
|---|---|---|---|
| | | A | G |
| α face (6) | A | | |
| | | A265(DA)\|\|A283(DA); 4v8b (4.2 Å, 7.6°, 33.7°, 38.6°, 142.8°) | A221(BA)\|\|G266(BA); 4u24 (3.4 Å, 20.8°, 10.6°, 24.5°, 135.9°) |
| | G | | |
| | | | G2458(1H)\|\|G2490(1H); 4wq1 (3.5 Å, 12.9°, 14.7°, 27.3°, 157.8°) |

**Table S64.** Examples (i.e. identities of bases involved in stacking and corresponding PDB codes) of consecutive (black) and non-consecutive (grey) 5||5 α||β *trans* purine||purine stacks. The geometrical parameters ($\vec{d}_{ab}$, $\theta_{ab}$, $\tau_a$, $\tau_b$ and $\sigma_{ab}$) for each example stack are provided in parentheses respectively.

| | | β face (5) | |
|---|---|---|---|
| | | A | G |
| α face (5) | A | A72(IB)\|\|A73(IB);6b4v (4.1 Å, 12.3°, 23.0°, 10.9°, 114.8°) | A460(6)\|\|G461(6);5fci (3.9 Å, 7.1°, 24.9°, 21.1°, 95.3°) |
| | | A571(1H)\|\|A575(1H);4wr6 (3.7 Å, 1.9°, 33.6°, 30.8°, 153.7°) | A2335(14)\|\|G2337(14);6gsj (3.3 Å, 21.1°, 29.1°, 15.1°,130.1°) |
| | G | | G2124(14)\|\|G2125(14);4wzd (3.9 Å, 3.5°, 25.7°, 28.9°,120.2°) |
| | | | G775(da)\|\|G794(da);4v8x (3.5 Å, 22.0°, 13.4°, 31.7°, 134.9°) |

**Table S65**. Examples (i.e. identities of bases involved in stacking and corresponding PDB codes) of consecutive (black) and non-consecutive (grey) 5||56, 6||56 β||α *cis* purine||purine stacks. The geometrical parameters ($\vec{d}_{ab}$, $\theta_{ab}$, $\tau_a$, $\tau_b$ and $\sigma_{ab}$) averaged for each ring:ring stacking contact (i.e. average parameters from 5||5, 5||6, 6||5 and 6||6 ring:ring contacts) for each example stack are provided in parentheses respectively.

| | | α face (56), (56) | |
|---|---|---|---|
| | | A | G |
| β face (5), (6) | A | | G1525(0)\|\|A1526(0);3ccj (3.9 Å, 22.4°, 26.2°, 21.4°, 73.4°) |
| | | | A2513(A)\|\|G2564(A);1fg0 (3.8 Å, 9.8°, 23.3°, 26.4°, 73.4°) |
| | G | | |

**Table S66**. Examples (i.e. identities of bases involved in stacking and corresponding PDB codes) of consecutive (black) and non-consecutive (grey) 5||6, 6||56 β||α *cis* purine||purine stacks. The geometrical parameters ($\vec{d}_{ab}$, $\theta_{ab}$, $\tau_a$, $\tau_b$ and $\sigma_{ab}$) averaged for each ring:ring stacking contact (i.e. average parameters from 5||6, 6||5 and 6||6 ring:ring contacts) for each example stack are provided in parentheses respectively.

| | | α face (6), (56) | |
|---|---|---|---|
| | | A | G |
| β face (5), (6) | A | | |
| | | | A109(XA)\|\|G326(XA);4www (4.2 Å, 22.6°, 19.0°, 25.9°, 84.1°) |
| | G | | |

**Table S67**. Examples (i.e. identities of bases involved in stacking and corresponding PDB codes) of consecutive (black) and non-consecutive (grey) 5||5, 6||56 β||α *cis* purine||purine stacks. The geometrical parameters ($\vec{d}_{ab}$, $\theta_{ab}$, $\tau_a$, $\tau_b$ and $\sigma_{ab}$) averaged for each ring:ring stacking contact (i.e. average parameters from 5||5, 6||5 and 6||6 ring:ring contacts) for each example stack are provided in parentheses respectively.

<table>
<tr><td colspan="2" rowspan="2"></td><td colspan="2">α face (5), (56)</td></tr>
<tr><td>A</td><td>G</td></tr>
<tr><td rowspan="3">β face (5), (6)</td><td rowspan="2">A</td><td rowspan="2"></td><td>G1878(5)||1879(5); 5fcj<br>(4.0 Å, 14.4°, 19.1°, 26.1°, 29.0°)</td></tr>
<tr><td>A103(CA)||G1031(CA); 4v63<br>(4.3 Å, 18.4°, 16.9°, 25.2°, 48.4°)</td></tr>
<tr><td>G</td><td></td><td></td></tr>
</table>

**Table S68**. Examples (i.e. identities of bases involved in stacking and corresponding PDB codes) of consecutive (black) and non-consecutive (grey) 5||56, 6||6 β||α *cis* purine||purine stacks. The geometrical parameters ($\vec{d}_{ab}$, $\theta_{ab}$, $\tau_a$, $\tau_b$ and $\sigma_{ab}$) averaged for each ring:ring stacking contact (i.e. average parameters from 5||5, 5||6 and 6||6 ring:ring contacts) for each example stack are provided in parentheses respectively.

<table>
<tr><td colspan="2" rowspan="2"></td><td colspan="2">α face (56), (6)</td></tr>
<tr><td>A</td><td>G</td></tr>
<tr><td rowspan="3">β face (5), (6)</td><td rowspan="2">A</td><td rowspan="2"></td><td>A1(D)||G2(D); 6dcl<br>(4.0 Å, 14.2°, 13.5°, 18.4°, 15.7°)</td></tr>
<tr><td>A109(AA)||G326(AA); 4u24<br>(3.6 Å, 11.6°, 25.3°, 24.2°, 61.2°)</td></tr>
<tr><td>G</td><td></td><td></td></tr>
</table>

**Table S69**. Examples (i.e. identities of bases involved in stacking and corresponding PDB codes) of consecutive (black) and non-consecutive (grey) 5||56, 6||5 β||α *cis* purine||purine stacks. The geometrical parameters ($\vec{d}_{ab}$, $\theta_{ab}$, $\tau_a$, $\tau_b$ and $\sigma_{ab}$) averaged for each ring:ring stacking contact (i.e. average parameters from 5||5, 5||6, and 6||5 ring:ring contacts) for each example stack are provided in parentheses respectively.

| | | α face (56), (5) | |
|---|---|---|---|
| | | A | G |
| β face (5), (6) | A | | |
| | | | A2513(A)\|\|G2564(A); 1ffz (3.5 Å, 15.1°, 26.6°, 25.8°, 77.6°) |
| | G | | |

**Table S70**. Examples (i.e. identities of bases involved in stacking and corresponding PDB codes) of consecutive (black) and non-consecutive (grey) 5||6, 6||6 β||α *cis* purine||purine stacks. The geometrical parameters ($\vec{d}_{ab}$, $\theta_{ab}$, $\tau_a$, $\tau_b$ and $\sigma_{ab}$) averaged for each ring:ring stacking contact (i.e. average parameters from 5||6 and 6||6 ring:ring contacts) for each example stack are provided in parentheses respectively.

| | | α face (6), (6) | |
|---|---|---|---|
| | | A | G |
| β face (5), (6) | A | | A33(A)\|\|G34(A); 3d2g (4.1 Å, 8.8°, 26.0°, 33.4°, 65.8°) |
| | | | A109(A)\|\|G326(A); 5wns (3.7 Å, 14.9°, 21.8°, 18.6°, 62.2°) |
| | G | | |

**Table S71**. Examples (i.e. identities of bases involved in stacking and corresponding PDB codes) of consecutive (black) and non-consecutive (grey) 6||56 β||α *cis* purine||purine stacks. The geometrical parameters ($\vec{d}_{ab}$, $\theta_{ab}$, $\tau_a$, $\tau_b$ and $\sigma_{ab}$) averaged for each ring:ring stacking contact (i.e. average parameters from 6||5 and 6||6 ring:ring contacts) for each example stack are provided in parentheses respectively.

| | | α face (56) | |
|---|---|---|---|
| | | A | G |
| β face (6) | A | | A496(BA)\|\|G497(BA); 4ybb<br>(4.3 Å, 8.5°, 39.2°, 32.4°, 57.2°) |
| | | | A1154(0)\|\|G2786(0); 3ccj<br>(3.8 Å, 9.8°, 26.8°, 27.5°, 65.4°) |
| | G | | |

**Table S72**. Examples (i.e. identities of bases involved in stacking and corresponding PDB codes) of consecutive (black) and non-consecutive (grey) 5||5, 6||6 β||α *cis* purine||purine stacks. The geometrical parameters ($\vec{d}_{ab}$, $\theta_{ab}$, $\tau_a$, $\tau_b$ and $\sigma_{ab}$) averaged for each ring:ring stacking contact (i.e. average parameters from 5||5 and 6||6 ring:ring contacts) for each example stack are provided in parentheses respectively.

| | | α face (5), (6) | |
|---|---|---|---|
| | | A | G |
| β face (5), (5) | A | | G2334(14)\|\|A2335(14); 5ndk<br>(3.9 Å, 17.1°, 32.9°, 27.8°, 9.4°) |
| | | | G2550(X)\|\|A2792(X); 5nrg<br>(4.4 Å, 1.9°, 37.6°, 37.4°, 2.5°) |
| | G | | |

**Table S73**. Examples (i.e. identities of bases involved in stacking and corresponding PDB codes) of consecutive (black) and non-consecutive (grey) 5||6, 6||5 β||α *cis* purine||purine stacks. The geometrical parameters ($\vec{d}_{ab}$, $\theta_{ab}$, $\tau_a$, $\tau_b$ and $\sigma_{ab}$) averaged for each ring:ring stacking contact (i.e. average parameters from 5||6 and 6||5 ring:ring contacts) for each example stack are provided in parentheses respectively.

| | | α face (6), (5) | |
|---|---|---|---|
| | | A | G |
| β face (5), (6) | A | | |
| | G | | |

**Table S74**. Examples (i.e. identities of bases involved in stacking and corresponding PDB codes) of consecutive (black) and non-consecutive (grey) 5||56 β||α *cis* purine||purine stacks. The geometrical parameters ($\vec{d}_{ab}$, $\theta_{ab}$, $\tau_a$, $\tau_b$ and $\sigma_{ab}$) averaged for each ring:ring stacking contact (i.e. average parameters from 5||5 and 5||6 ring:ring contacts) for each example stack are provided in parentheses respectively.

| | | α face (56) | |
|---|---|---|---|
| | | A | G |
| β face (5) | A | | G2319(1H)\|\|A2320(1H); 5el4<br>(4.1 Å, 21.7°, 29.8°, 22.1°, 9.4°)<br>A13(B)\|\|G37(B); 2qus<br>(3.8 Å, 11.0°, 28.3°, 30.0°, 14.9°) |
| | G | | |

**Table S75**. Examples (i.e. identities of bases involved in stacking and corresponding PDB codes) of consecutive (black) and non-consecutive (grey) 5||5, 6||5 β||α *cis* purine||purine stacks. The geometrical parameters ($\vec{d}_{ab}$, $\theta_{ab}$, $\tau_a$, $\tau_b$ and $\sigma_{ab}$) averaged for each ring:ring stacking contact (i.e. average parameters from 5||5 and 6||5 ring:ring contacts) for each example stack are provided in parentheses respectively.

| | | α face (5), (5) | |
|---|---|---|---|
| | | A | G |
| β face (5), (6) | A | | |
| | | | A2478(RA)\|\|G2529(RA); 4lt8 (3.8Å, 22.9°,23.7°, 19.0°, 88.4°) |
| | G | | |

**Table S76**. Examples (i.e. identities of bases involved in stacking and corresponding PDB codes) of consecutive (black) and non-consecutive (grey) 6||6 β||α *cis* purine||purine stacks. The geometrical parameters ($\vec{d}_{ab}$, $\theta_{ab}$, $\tau_a$, $\tau_b$ and $\sigma_{ab}$) for each example stack are provided in parentheses respectively.

| | | α face (6) | |
|---|---|---|---|
| | | A | G |
| β face (6) | A | | A159(A)\|\|G160(A);4fax (3.9 Å, 13.5°, 37.0°, 23.5°, 22.8°) |
| | | | G2558(A)\|\|G2800(A);1k8a (4.3 Å, 3.2°, 36.5°, 39.5°, 6.4°) |
| | G | | |

**Table S77**. Examples (i.e. identities of bases involved in stacking and corresponding PDB codes) of consecutive (black) and non-consecutive (grey) 5||6 β||α *cis* purine||purine stacks. The geometrical parameters ($\vec{d}_{ab}$, $\theta_{ab}$, $\tau_a$, $\tau_b$ and $\sigma_{ab}$) for each example stack are provided in parentheses respectively.

| | | α face (6) | |
|---|---|---|---|
| | | A | G |
| β face (5) | A | | A163(B)\|\|G164(B); 488d<br>(3.3 Å, 7.7°, 14.7°, 14.9°, 10.5°) |
| | | | A93(2)\|\|G398(2); 5ndv<br>(3.4 Å, 16.1°, 13.4°, 25.4°, 46.9°) |
| | G | | |

**Table S78**. Examples (i.e. identities of bases involved in stacking and corresponding PDB codes) of consecutive (black) and non-consecutive (grey) 6||5 β||α *cis* purine||purine stacks. The geometrical parameters ($\vec{d}_{ab}$, $\theta_{ab}$, $\tau_a$, $\tau_b$ and $\sigma_{ab}$) for each example stack are provided in parentheses respectively.

| | | α face (5) | |
|---|---|---|---|
| | | A | G |
| β face (6) | A | | A496(AA)\|\|G497(AA); 4woi<br>(4.0 Å, 10.7°, 38.2°, 38.4°, 54.2) |
| | | | A84(B)\|\|G95(B); 5xbl<br>(3.8 Å, 12.5°, 38.9°, 27.7°, 55.2°) |
| | G | | |

**Table S79**. Examples (i.e. identities of bases involved in stacking and corresponding PDB codes) of consecutive (black) and non-consecutive (grey) 5||5 β||α *cis* purine||purine stacks. The geometrical parameters ($\vec{d}_{ab}$, $\theta_{ab}$, $\tau_a$, $\tau_b$ and $\sigma_{ab}$) for each example stack are provided in parentheses respectively.

<table>
<tr><td colspan="2" rowspan="2"></td><td colspan="2">α face (5)</td></tr>
<tr><td>A</td><td>G</td></tr>
<tr><td rowspan="3">β face (5)</td><td rowspan="2">A</td><td rowspan="2"></td><td>G2319(14)||A2320(14);5ibb<br>(4.1 Å, 19.1°, 32.5°, 22.7°, 18.0°)</td></tr>
<tr><td>G1087 (BB)|| A1089 (BB);4v57<br>(3.8 Å, 10.5°, 23.1°, 28.9°, 49.3°)</td></tr>
<tr><td>G</td><td></td><td></td></tr>
</table>

**Table S80**. Examples (i.e. identities of bases involved in stacking and corresponding PDB codes) of consecutive (black) and non-consecutive (grey) 5||56, 6||56 β||α *trans* purine||purine stacks. The geometrical parameters ($\vec{d}_{ab}$, $\theta_{ab}$, $\tau_a$, $\tau_b$ and $\sigma_{ab}$) averaged for each ring:ring stacking contact (i.e. average parameters from 5||5, 5||6, 6||5 and 6||6 ring:ring contacts) for each example stack are provided in parentheses respectively.

<table>
<tr><td colspan="2" rowspan="2"></td><td colspan="2">α face (56), (56)</td></tr>
<tr><td>A</td><td>G</td></tr>
<tr><td rowspan="3">β face<br>(5),(6)</td><td rowspan="2">A</td><td rowspan="2"></td><td></td></tr>
<tr><td>A2404(BA)||G2441(BA);4w2g<br>(3.8 Å, 3.1°,24.0°, 24.7°, 163.2°)</td></tr>
<tr><td>G</td><td></td><td></td></tr>
</table>

**Table S81**. Examples (i.e. identities of bases involved in stacking and corresponding PDB codes) of consecutive (black) and non-consecutive (grey) 5||6, 6||56 β||α *trans* purine||purine stacks. The geometrical parameters ($\vec{d}_{ab}$, $\theta_{ab}$, $\tau_a$, $\tau_b$ and $\sigma_{ab}$) averaged for each ring:ring stacking contact (i.e. average parameters from 5||6, 6||5 and 6||6 ring:ring contacts) for each example stack are provided in parentheses respectively.

<table>
<tr><td colspan="2" rowspan="2"></td><td colspan="2">α face (6), (56)</td></tr>
<tr><td>A</td><td>G</td></tr>
<tr><td rowspan="3">β face (5), (6)</td><td rowspan="2">A</td><td rowspan="2"></td><td></td></tr>
<tr><td>G778(2)||A780(2); 5dgv<br>(4.0 Å,13.7°, 22.4°, 23.9°, 136.8°)</td></tr>
<tr><td>G</td><td></td><td></td></tr>
</table>

**Table S82**. Examples (i.e. identities of bases involved in stacking and corresponding PDB codes) of consecutive (black) and non-consecutive (grey) 5||5, 6||56 β||α *trans* purine||purine stacks. The geometrical parameters ($\vec{d}_{ab}$, $\theta_{ab}$, $\tau_a$, $\tau_b$ and $\sigma_{ab}$) averaged for each ring:ring stacking contact (i.e. average parameters from 5||5, 6||5 and 6||6 ring:ring contacts) for each example stack are provided in parentheses respectively.

<table>
<tr><td colspan="2" rowspan="2"></td><td colspan="2">α face (5), (56)</td></tr>
<tr><td>A</td><td>G</td></tr>
<tr><td rowspan="3">β face (5), (6)</td><td rowspan="2">A</td><td rowspan="2"></td><td></td></tr>
<tr><td>A2847(1)||G2898(1); 4u55<br>(3.7 Å, 20.0°, 23.1°, 21.2°, 101.1°)</td></tr>
<tr><td>G</td><td></td><td></td></tr>
</table>

**Table S83**. Examples (i.e. identities of bases involved in stacking and corresponding PDB codes) of consecutive (black) and non-consecutive (grey) 5||56, 6||6 β||α *trans* purine||purine stacks. The geometrical parameters ($\vec{d}_{ab}$, $\theta_{ab}$, $\tau_a$, $\tau_b$ and $\sigma_{ab}$) averaged for each ring:ring stacking contact (i.e. average parameters from 5||5, 5||6 and 6||6 ring:ring contacts) for each example stack are provided in parentheses respectively.

| | | α face (56), (6) | |
|---|---|---|---|
| | | A | G |
| β face (5), (6) | A | | |
| | | | A2457(0)\|\|G2508(0); 2aar<br>(4.1 Å, 9.8°, 19.5°, 20.5°, 101.6°) |
| | G | | |

**Table S84**. Examples (i.e. identities of bases involved in stacking and corresponding PDB codes) of consecutive (black) and non-consecutive (grey) 5||56, 6||5 β||α *trans* purine||purine stacks. The geometrical parameters ($\vec{d}_{ab}$, $\theta_{ab}$, $\tau_a$, $\tau_b$ and $\sigma_{ab}$) averaged for each ring:ring stacking contact (i.e. average parameters from 5||5, 5||6 and 6||5 ring:ring contacts) for each example stack are provided in parentheses respectively.

| | | α face (56), (5) | |
|---|---|---|---|
| | | A | G |
| β face (5), (6) | A | | |
| | | | A2457(A)\|\|G2508(A);1jzy<br>(3.9 Å, 19.5°, 21.8°, 19.6°, 101.0°) |
| | G | | |

**Table S85**. Examples (i.e. identities of bases involved in stacking and corresponding PDB codes) of consecutive (black) and non-consecutive (grey) 5||6, 6||6 β||α *trans* purine||purine stacks. The geometrical parameters ($\vec{d}_{ab}$, $\theta_{ab}$, $\tau_a$, $\tau_b$ and $\sigma_{ab}$) averaged for each ring:ring stacking contact (i.e. average parameters from 5||6 and 6||6 ring:ring contacts) for each example stack are provided in parentheses respectively.

<table>
<tr><td colspan="2" rowspan="2"></td><td colspan="2">α face (6), (6)</td></tr>
<tr><td>A</td><td>G</td></tr>
<tr><td rowspan="3">β face (5), (6)</td><td rowspan="2">A</td><td rowspan="2"></td><td></td></tr>
<tr><td>A812(2)||G858(2); 4u55<br>(3.9 Å, 4.4°, 22.4°, 25.3°, 135.4°)</td></tr>
<tr><td>G</td><td></td><td></td></tr>
</table>

**Table S86**. Examples (i.e. identities of bases involved in stacking and corresponding PDB codes) of consecutive (black) and non-consecutive (grey) 6||56 β||α *trans* purine||purine stacks. The geometrical parameters ($\vec{d}_{ab}$, $\theta_{ab}$, $\tau_a$, $\tau_b$ and $\sigma_{ab}$) averaged for each ring:ring stacking contact (i.e. average parameters from 6||5 and 6||6 ring:ring contacts) for each example stack are provided in parentheses respectively.

<table>
<tr><td colspan="2" rowspan="2"></td><td colspan="2">α face (56)</td></tr>
<tr><td>A</td><td>G</td></tr>
<tr><td rowspan="3">β face (6)</td><td rowspan="2">A</td><td rowspan="2"></td><td></td></tr>
<tr><td>G778(A)||A780(A); 6hhq<br>(3.6 Å, 8.0°, 19.4°, 25.3°, 129.0°)</td></tr>
<tr><td>G</td><td></td><td></td></tr>
</table>

**Table S87**. Examples (i.e. identities of bases involved in stacking and corresponding PDB codes) of consecutive (black) and non-consecutive (grey) 5||5, 6||6 β||α *trans* purine||purine stacks. The geometrical parameters ($\vec{d}_{ab}$, $\theta_{ab}$, $\tau_a$, $\tau_b$ and $\sigma_{ab}$) averaged for each ring:ring stacking contact (i.e. average parameters from 5||5 and 6||6 ring:ring contacts) for each example stack are provided in parentheses respectively.

| | | α face (5), (6) | |
|---|---|---|---|
| | | A | G |
| β face (5), (6) | A | | |
| | G | | |

**Table S88**. Examples (i.e. identities of bases involved in stacking and corresponding PDB codes) of consecutive (black) and non-consecutive (grey) 5||6, 6||5 β||α *trans* purine||purine stacks. The geometrical parameters ($\vec{d}_{ab}$, $\theta_{ab}$, $\tau_a$, $\tau_b$ and $\sigma_{ab}$) averaged for each ring:ring stacking contact (i.e. average parameters from 5||6 and 6||5 ring:ring contacts) for each example stack are provided in parentheses respectively.

| | | α face (6), (5) | |
|---|---|---|---|
| | | A | G |
| β face (5), (6) | A | | |
| | | | G2157(1)\|A2178(1);4u3n<br>(3.9 Å, 12.2°, 26.5, 34.0°, 164.4°) |
| | G | | |

**Table S89**. Examples (i.e. identities of bases involved in stacking and corresponding PDB codes) of consecutive (black) and non-consecutive (grey) 5||56 β||α *trans* purine||purine stacks. The geometrical parameters ($\vec{d}_{ab}$, $\theta_{ab}$, $\tau_a$, $\tau_b$ and $\sigma_{ab}$) averaged for each ring:ring stacking contact (i.e. average parameters from 5||5 and 5||6 ring:ring contacts) for each example stack are provided in parentheses respectively.

| | | α face (56) | |
|---|---|---|---|
| | | A | G |
| β face (5) | A | | |
| | | | A53(A)\|\|G83(A); 2hop<br>(3.8 Å, 15.9°, 29.2°, 19.1°, 140.8°) |
| | G | | |

**Table S90**. Examples (i.e. identities of bases involved in stacking and corresponding PDB codes) of consecutive (black) and non-consecutive (grey) 5||5, 6||5 β||α *trans* purine||purine stacks. The geometrical parameters ($\vec{d}_{ab}$, $\theta_{ab}$, $\tau_a$, $\tau_b$ and $\sigma_{ab}$) averaged for each ring:ring stacking contact (i.e. average parameters from 5||6 and 6||5 ring:ring contacts) for each example stack are provided in parentheses respectively.

| | | α face (5), (5) | |
|---|---|---|---|
| | | A | G |
| β face (5), (6) | A | | |
| | | | A2478(BA)\|\|G2529(BA); 4v9l (3.6 Å, 22.0°, 29.3°, 20.0°, 93.7°) |
| | G | | |

**Table S91**. Examples (i.e. identities of bases involved in stacking and corresponding PDB codes) of consecutive (black) and non-consecutive (grey) 6||6 β||α *trans* purine||purine stacks The geometrical parameters ($\vec{d}_{ab}$, $\theta_{ab}$, $\tau_a$, $\tau_b$ and $\sigma_{ab}$) for each example stack are provided in parentheses respectively.

| | | α face (6) | |
|---|---|---|---|
| | | A | G |
| β face (6) | A | | |
| | | | G721(AA)\|\|A733(AA);4v5e (3.9 Å, 15.3°, 36.3°, 29.0°, 159.5°) |
| | G | | |

**Table S92**. Examples (i.e. identities of bases involved in stacking and corresponding PDB codes) of consecutive (black) and non-consecutive (grey) 5||6 β||α *trans* purine||purine stacks. The geometrical parameters ($\vec{d}_{ab}$, $\theta_{ab}$, $\tau_a$, $\tau_b$ and $\sigma_{ab}$) for each example stack are provided in parentheses respectively.

| | | α face (6) | |
|---|---|---|---|
| | | A | G |
| β face (5) | A | | |
| | | | A803(1A)\|\|G1507(1A); 5hcq (3.9 Å, 15.9°, 39.5°, 26.2°, 120.3°) |
| | G | | |

**Table S93**. Examples (i.e. identities of bases involved in stacking and corresponding PDB codes) of consecutive (black) and non-consecutive (grey) 6||5 β||α *trans* purine||purine stacks. The geometrical parameters ($\vec{d}_{ab}$, $\theta_{ab}$, $\tau_a$, $\tau_b$ and $\sigma_{ab}$) for each example stack are provided in parentheses respectively.

| | | α face (5) | |
|---|---|---|---|
| | | A | G |
| β face (6) | A | | |
| | | | A2810(X)\|\|G2854(X); 3pip<br>(3.8 Å, 6.9°, 24.0°, 30.9°, 145.3°) |
| | G | | |

**Table S94**. Examples (i.e. identities of bases involved in stacking and corresponding PDB codes) of consecutive (black) and non-consecutive (grey) 5||5 β||α *trans* purine||purine stacks. The geometrical parameters ($\vec{d}_{ab}$, $\theta_{ab}$, $\tau_a$, $\tau_b$ and $\sigma_{ab}$) for each example stack are provided in parentheses respectively.

| | | α face (5) | |
|---|---|---|---|
| | | A | G |
| β face (5) | A | | |
| | | | A577(X)\|\|G2048(X);4wf9<br>(3.9 Å, 7.3°, 37.2°, 31.8°, 153.6°) |
| | G | | |

**Table S95**. Examples (i.e. identities of bases involved in stacking and corresponding PDB codes) of consecutive (black) and non-consecutive (grey) 5||56, 6||56 β||β *cis* purine||purine stacks, The geometrical parameters ($\vec{d}_{ab}$, $\theta_{ab}$, $\tau_a$, $\tau_b$ and $\sigma_{ab}$) averaged for each ring:ring stacking contact (i.e. average parameters from 5||5, 5||6, 6||5 and 6||6 ring:ring contacts) for each example stack are provided in parentheses respectively.

| | | β face (56), (56) | |
|---|---|---|---|
| | | A | G |
| β face (5),(6) | A | A61(2)\|\|A62(2); 5dgv<br>(4.2 Å, 22.0°, 20.4°, 26.9°, 64.3°) | A493(CA)\|\|G494(CA); 4v6c<br>(4.2 Å, 20.4°, 23.0°, 20.7°, 58.4°) |
| | | A665(AA)\|\|A733(AA); 4v7y<br>(3.6 Å, 10.0°, 23.0°, 20.8°, 1.0°) | A50(B) \|\| G68(B); 2h0x<br>(3.8 Å, 3.7°, 24.6°, 23.2°, 8.2°) |
| | G | | G1242(5)\|\|G1343(5);5dat<br>(4.0 Å, 15.4°,22.3°, 27.7°, 63.8°) |
| | | | G9(2x)\|\|G46(2x); 5hd1<br>(4.1 Å, 8.7°, 22.9°, 20.3°, 1.5°) |

**Table S96**. Examples (i.e. identities of bases involved in stacking and corresponding PDB codes) of consecutive (black) and non-consecutive (grey) 5||6, 6||56 β||β *cis* purine||purine stacks. , The geometrical parameters ($\vec{d}_{ab}$, $\theta_{ab}$, $\tau_a$, $\tau_b$ and $\sigma_{ab}$) averaged for each ring:ring stacking contact (i.e. average parameters from 5||6, 6||5 and 6||6 ring:ring contacts) for each example stack are provided in parentheses respectively.

| | | β face (6), (56) | |
|---|---|---|---|
| | | A | G |
| β \|\|β face (5), (6) | A | A1755 (2)\|\|A1756(2);5dgf<br>(4.1 Å, 13.7°, 17.4°, 13.9°, 33.2°) | G1878 (1)\|\| A1879 (1);5tbw<br>(4.0 Å, 21.8°, 24.3°, 16.2°, 1.1°) |
| | | A65(1)\|\|A77(1);5dc3<br>(4.0 Å, 4.3°, 24.8°, 24.5°, 28.9°) | A2806(1H)\|\|G2906(1H);5ndk<br>(4.1 Å, 19.0°, 27.4°, 18.0°, 20.6°) |
| | G | | G901(6)\|\|G902(6);5dgv<br>(4.0 Å, 22.9°, 23.9°, 28.9°, 74.6°) |
| | | | G2046(BA)\|\|G2623(BA); 4u24<br>(3.9Å, 5.5°, 30.3°, 26.3°, 46.0°) |

**Table S97**. Examples (i.e. identities of bases involved in stacking and corresponding PDB codes) of consecutive (black) and non-consecutive (grey) 5||5, 6||56 β||β *cis* purine||purine stacks. , The geometrical parameters ($\vec{d}_{ab}$, $\theta_{ab}$, $\tau_a$, $\tau_b$ and $\sigma_{ab}$) averaged for each ring:ring stacking contact (i.e. average parameters from 5||5, 6||5 and 6||6 ring:ring contacts) for each example stack are provided in parentheses respectively.

| | | β face (5), (56) | |
|---|---|---|---|
| | | A | G |
| β face (5), (6) | A | A1507(BA)\|\|A1508(BA); 4v9a<br>(3.9 Å, 21.4°, 24.3°,21.8°, 46.7°) | A646(DA)\|\|G647(DA); 4v9k<br>(4.0 Å, 15.2°, 20.1°, 25.0°, 51.1°) |
| | | A191(0)\|\|A204(0);3ccm<br>(4.1 Å, 9.6°, 24.2°, 26.7°, 26.3°) | A722(2a)\|\|G724(2a);6nd5<br>(3.7 Å, 10.2°, 20.6°, 22.4°, 69.9°) |
| | G | | G1242(5)\|\|G1243(5); 4u55<br>(3.7 Å, 18.4°, 20.6°, 24.8°, 72.1°) |
| | | | G1459(BA)\|\|G1461(BA); 4v5d<br>(3.7 Å, 5.8°, 24.2°, 25.0°, 43.8°) |

**Table S98.** Examples (i.e. identities of bases involved in stacking and corresponding PDB codes) of consecutive (black) and non-consecutive (grey) 5||56, 6||6 β||β *cis* purine||purine stacks. The geometrical parameters ($\vec{d}_{ab}$, $\theta_{ab}$, $\tau_a$, $\tau_b$ and $\sigma_{ab}$) averaged for each ring:ring stacking contact (i.e. average parameters from 5||5, 5||6 and 6||6 ring:ring contacts) for each example stack are provided in parentheses respectively.

| | | β face (56), (6) | |
|---|---|---|---|
| | | A | G |
| β face (5), (6) | A | | A646(1H)\|\|G647(1H); 4wt1<br>(4.0 Å, 22.6°, 30.8°, 16.5°, 76.3°) |
| | | | A1815(2A)\|\|G1817(2A); 6cfk<br>(3.7 Å, 5.6°, 23.1°, 25.1°, 55.3°) |
| | G | | |

**Table S99.** Examples (i.e. identities of bases involved in stacking and corresponding PDB codes) of consecutive (black) and non-consecutive (grey) 5||56, 6||5 β||β *cis* purine||purine stacks. , The geometrical parameters ($\vec{d}_{ab}$, $\theta_{ab}$, $\tau_a$, $\tau_b$ and $\sigma_{ab}$) averaged for each ring:ring stacking contact (i.e. average parameters from 5||5, 5||6 and 6||5 ring:ring contacts) for each example stack are provided in parentheses respectively.

| | | β face (56), (5) | |
|---|---|---|---|
| | | A | G |
| β face (5), (6) | A | A1556(1A)\|\|A1557(1A); 6cfk<br>(3.8 Å, 17.6°, 23.6°, 28.6°, 55.1°) | G1604(5)\|\|A1605(5); 4u4y<br>(3.6 Å, 9.7°, 14.4°, 16.8°, 21.0°) |
| | | A665(a)\|\|A733(a); 4jv5<br>(3.6 Å, 13.2°, 21.2°, 16.1°, 4.1°) | G3242(1)\|\|A3245(1); 4u4u<br>(3.9 Å, 11.2°, 15.4°, 17.0°, 14.6°) |
| | G | | |
| | | | G420(1)\|\|G2385(1); 4u4u<br>(3.7 Å, 22.7°, 18.8°, 31.3°, 8.4°) |

**Table S100.** Examples (i.e. identities of bases involved in stacking and corresponding PDB codes) of consecutive (black) and non-consecutive (grey) 5||6, 6||6 β||β *cis* purine||purine stacks. , The geometrical parameters ($\vec{d}_{ab}$, $\theta_{ab}$, $\tau_a$, $\tau_b$ and $\sigma_{ab}$) averaged for each ring:ring stacking contact (i.e. average parameters from 5||6 and 6||6 ring:ring contacts) for each example stack are provided in parentheses respectively.

<table>
<tr><td colspan="2" rowspan="2"></td><td colspan="2">β face (6), (6)</td></tr>
<tr><td>A</td><td>G</td></tr>
<tr><td rowspan="4">β face (5), (6)</td><td rowspan="2">A</td><td>A141(BA)||A142(BA); 4v8g<br>(4.1 Å, 22.3°, 21.2°, 15.2°, 67.3°)</td><td>A493(CA)||G494(CA); 4v7u<br>(3.9 Å, 22.9°, 14.8°, 23.4°, 67.6)</td></tr>
<tr><td>A43(6)||A378(6); 4u6f<br>(3.5 Å, 6.4°, 22.1°, 25.8°, 2.8°)</td><td>A14(C)||G22(C); 5wwr<br>(4.0 Å, 12.2°, 37.5°, 34.3°, 47.8°)</td></tr>
<tr><td rowspan="2">G</td><td rowspan="2"></td><td></td></tr>
<tr><td>G77(DA)||G110(DA); 4u24<br>(3.8 Å, 11.5°, 15.3°, 23.1°, 22.8°)</td></tr>
</table>

**Table S101.** Examples (i.e. identities of bases involved in stacking and corresponding PDB codes) of consecutive (black) and non-consecutive (grey) 6||56 β||β *cis* purine||purine stacks. , The geometrical parameters ($\vec{d}_{ab}$, $\theta_{ab}$, $\tau_a$, $\tau_b$ and $\sigma_{ab}$) averaged for each ring:ring stacking contact (i.e. average parameters from 6||5 and 6||6 ring:ring contacts) for each example stack are provided in parentheses respectively.

<table>
<tr><td colspan="2" rowspan="2"></td><td colspan="2">β face (56)</td></tr>
<tr><td>A</td><td>G</td></tr>
<tr><td rowspan="3">β face (6)</td><td rowspan="2">A</td><td rowspan="2"></td><td>A878(14)||G879(14); 4wq1<br>(4.2 Å, 15.0°, 22.3°, 25.9°, 76.4°)</td></tr>
<tr><td>A673(AA)||G734(AA); 4u24<br>(4.0 Å, 13.0°, 25.8°, 22.6°, 40.3)</td></tr>
<tr><td>G</td><td></td><td></td></tr>
</table>

**Table S102.** Examples (i.e. identities of bases involved in stacking and corresponding PDB codes) of consecutive (black) and non-consecutive (grey) 5||5, 6||6 β||β *cis* purine||purine stacks. , The geometrical parameters ($\vec{d}_{ab}$, $\theta_{ab}$, $\tau_a$, $\tau_b$ and $\sigma_{ab}$) averaged for each ring:ring stacking contact (i.e. average parameters from 5||5 and 6||6 ring:ring contacts) for each example stack are provided in parentheses respectively.

| | | β face (5), (6) | |
|---|---|---|---|
| | | A | G |
| β face (5), (6) | A | | |
| | G | | |

**Table S103.** Examples (i.e. identities of bases involved in stacking and corresponding PDB codes) of consecutive (black) and non-consecutive (grey) 5||6, 6||5 β||β *cis* purine||purine stacks. , The geometrical parameters ($\vec{d}_{ab}$, $\theta_{ab}$, $\tau_a$, $\tau_b$ and $\sigma_{ab}$) averaged for each ring:ring stacking contact (i.e. average parameters from 5||6 and 6||5 ring:ring contacts) for each example stack are provided in parentheses respectively.

| | | β face (6), (5) | |
|---|---|---|---|
| | | A | G |
| β face (5), (6) | A | A1755(6)\|\|A1756(6); 5dat (4.1 Å, 14.0°, 2.4°, 11.8°, 23.5°) | G1878(1)\|\| A1879(1);5dgf (4.2 Å, 22.8°, 28.9°, 30.4°, 13.2°) |
| | | A1306(CA)\|\|A1332(CA); 4v7w (4.2 Å, 21.4°, 10.5°, 10.5°, 24.3°) | G1541 (1)\|\| A1557 (1);5on6 (4.1 Å, 12.3°, 33.0°, 39.1°,43.5°) |
| | G | | |
| | | | G77(BA))\|\| G93(BA): 4v8q (3.7 Å, 19.3°, 17.7°, 34.9°, 37.8°) |

**Table S104.** Examples (i.e. identities of bases involved in stacking and corresponding PDB codes) of consecutive (black) and non-consecutive (grey) 5||56 β||β *cis* purine||purine stacks. , The geometrical parameters ($\vec{d}_{ab}$, $\theta_{ab}$, $\tau_a$, $\tau_b$ and $\sigma_{ab}$) averaged for each ring:ring stacking contact (i.e. average parameters from 5||5 and 5||6 ring:ring contacts) for each example stack are provided in parentheses respectively.

| | | β face (56) | |
|---|---|---|---|
| | | A | G |
| β face (5) | A | A61(2)\|\|A62(2); 5ndv<br>(3.9 Å, 18.2°, 17.3°, 21.2°, 55.1°) | A56(R)\|\|G57(R); 6b14<br>(4.2 Å, 13.0°, 25.9°, 37.2°, 12.1°) |
| | | A2333(DB)\|\|A2335(DB); 4v64<br>(3.7 Å, 12.5°, 19.6, 16.2°, 49.2°) | A1815(BA)\|\|G1817(BA); 4u24<br>(3.5 Å, 4.9°, 17.3°, 19.1°, 55.3°) |
| | G | | G138(1H)\|\|G139(1H); 4wq1<br>(4.2 Å, 21.7°, 14.3°, 25.6°, 67.7°) |
| | | | G2630(BA)\|\|G2894(BA); 4v8n<br>(3.8 Å, 10.6°, 26.8°, 20.4°, 16.8°) |

**Table S105.** Examples (i.e. identities of bases involved in stacking and corresponding PDB codes) of consecutive (black) and non-consecutive (grey) 5||5, 6||5 β||β *cis* purine||purine stacks. The geometrical parameters ($\vec{d}_{ab}$, $\theta_{ab}$, $\tau_a$, $\tau_b$ and $\sigma_{ab}$) averaged for each ring:ring stacking contact (i.e. average parameters from 5||5 and 6||5 ring:ring contacts) for each example stack are provided in parentheses respectively.

| | | β face (5), (5) | |
|---|---|---|---|
| | | A | G |
| β face (5), (6) | A | | A1050(BA)\|\|G1051(BA); 4v8b<br>(3.9 Å, 8.7°, 21.7°, 24.3°, 49.3°) |
| | | | A50(A)\|\|G169(A); 4e8t<br>(3.5 Å, 11.0°, 22.3°, 17.7°, 18.5°) |
| | G | | |

**Table S106.** Examples (i.e. identities of bases involved in stacking and corresponding PDB codes) of consecutive (black) and non-consecutive (grey) 6||6 β||β *cis* purine||purine stacks. The geometrical parameters ($\vec{d}_{ab}$, $\theta_{ab}$, $\tau_a$, $\tau_b$ and $\sigma_{ab}$) for each example stack are provided in parentheses respectively.

| | | β face (6) | |
|---|---|---|---|
| | | A | G |
| β face (6) | A | A144(F)\|\|A145(F);3k0j (4.4 Å, 1.2°, 33.4°, 33.2°, 0.9°) | A17(C)\|\| G18(C);5dea (4.2 Å, 6.0°, 36.5°, 39.6°, 20.5°) |
| | | A6(A)\|\|A113(A);4kqy (3.8 Å, 6.8°, 32.0°, 38.1°, 22.8°) | A507(B)\|\|G567(B);1g59 (3.6 Å, 11.5°, 36.1°, 26.7°, 6.1°) |
| | G | | G234 (6)\|\|G235 (6);5fci (4.5 Å, 20.2°, 29.4°, 27.9°, 46.0°) |
| | | | G528(B)\|\|G543(B);1g59 (3.5 Å, 4.1°, 12.3°, 16.4°, 19.8°) |

**Table S107.** Examples (i.e. identities of bases involved in stacking and corresponding PDB codes) of consecutive (black) and non-consecutive (grey) 5||6 β||β *cis* purine||purine stacks. The geometrical parameters ($\vec{d}_{ab}$, $\theta_{ab}$, $\tau_a$, $\tau_b$ and $\sigma_{ab}$) for each example stack are provided in parentheses respectively.

| | | β face (6) | |
|---|---|---|---|
| | | A | G |
| β face (5) | A | | G921(0)\|\|A922(0); 3ccu (4.0 Å, 22.6°, 28.5°, 32.8°, 9.4°) |
| | | | A1522(DA)\|\|G1524(DA); 4u24 (4.5 Å, 18.9°, 35.1°, 21.0°, 72.4° ) |
| | G | | |

**Table S108.** Examples (i.e. identities of bases involved in stacking and corresponding PDB codes) of consecutive (black) and non-consecutive (grey) 6||5 β||β *cis* purine||purine stacks. The geometrical parameters ($\vec{d}_{ab}$, $\theta_{ab}$, $\tau_a$, $\tau_b$ and $\sigma_{ab}$) for each example stack are provided in parentheses respectively.

| | | β face (5) | |
|---|---|---|---|
| | | A | G |
| β face (6) | A | A22(Ax)\|\|A23(Ax);4v7j (3.6 Å, 17.3°, 9.3°, 25.8°, 58.6°) | A2598(H)\|\|G2599(H); 3dh3 (4.0 Å, 15.1°, 26.5°, 34.1°, 53.7°) |
| | | A975(BA)\|\|A990(BA); 4u24 (3.2 Å, 8.8°, 23.7°, 14.9°, 69.7°) | A876(0)\|\|G878(0); 3ccj (3.9 Å, 4.0°, 39.0°, 36.6°, 69.3°,) |
| | G | | G1032(CA)\|\|G1033(CA); 4v57 (4.0 Å, 22.9°, 21.5°, 31.1°, 30.4°) |
| | | | G1009(AA)\|\|G1021(AA); 4v8n (4.0 Å, 18.6°, 32.8°, 30.8°, 38.5°) |

**Table S109.** Examples (i.e. identities of bases involved in stacking and corresponding PDB codes) of consecutive (black) and non-consecutive (grey) 5||5 β||β *cis* purine||purine stacks. The geometrical parameters ($\vec{d}_{ab}$, $\theta_{ab}$, $\tau_a$, $\tau_b$ and $\sigma_{ab}$) for each example stack are provided in parentheses respectively.

| | | β face (5) | |
|---|---|---|---|
| | | A | G |
| β face (5) | A | A113(A)\|\|A114(A);1kc8 (3.8 Å, 15.4°, 31.8°, 20.2°, 72.8°) | G1503(2)\|\|A1504(2);5dat (3.6 Å, 22.4°, 19.2°, 21.0°,65.2°) |
| | | A1543(B)\|\|A1545(B);6b4v (3.6 Å, 12.0°, 27.0°, 19.6°, 8.8°) | A560(AA)\|\|G566(AA); 4u24 (3.8 Å, 23.0°, 37.8°, 20.4°, 73.2°) |
| | G | | G142(6)\|\|G143(6);5on6 (4.1 Å, 9.3°, 23.6°, 28.3°, 42.0°) |
| | | | G7790)\|\|G778(0);1xbp (3.3 Å, 9.3°, 26.7°, 18.6°, 25.0°) |

**Table S110**. Examples (i.e. identities of bases involved in stacking and corresponding PDB codes) of consecutive (black) and non-consecutive (grey) 5||56, 6||56 β||β *trans* purine||purine stacks The geometrical parameters ($\vec{d}_{ab}$, $\theta_{ab}$, $\tau_a$, $\tau_b$ and $\sigma_{ab}$) averaged for each ring:ring stacking contact (i.e. average parameters from 5||5, 5||6, 6||5 and 6||6 ring:ring contacts) for each example stack are provided in parentheses respectively.

| | | β face (56), (56) | |
|---|---|---|---|
| | | A | G |
| β face (5), (6) | A | A14(4k)\|\|A15(4k);4wzo<br>(3.8 Å, 7.8°, 25.3°, 22.9°, 95.6°) | A27(CB)\|\|G28(CB);4v8f<br>(3.8 Å, 6.8°, 24.7°, 21.2°, 103.4°) |
| | | A1773(DA)\|\|A1839(DA);5j88<br>(3.9 Å, 8.6°, 28.0°, 30.3°, 149.3°) | G2097(A)\|\|A2612(A) ;1ffz<br>(4.2 Å, 7.6°, 22.2°, 23.4°, 125.6°) |
| | G | | |
| | | | G94(B)\|\|G138(B); 2gcv<br>(3.7 Å, 7.4°, 24.8°, 26.0°, 101.2) |

**Table S111.** Examples (i.e. identities of bases involved in stacking and corresponding PDB codes) of consecutive (black) and non-consecutive (grey) 5||6, 6||56 β||β *trans* purine||purine stacks. The geometrical parameters ($\vec{d}_{ab}$, $\theta_{ab}$, $\tau_a$, $\tau_b$ and $\sigma_{ab}$) averaged for each ring:ring stacking contact (i.e. average parameters from 5||6, 6||5 and 6||6 ring:ring contacts) for each example stack are provided in parentheses respectively.

| | | β face (6), (56) | |
|---|---|---|---|
| | | A | G |
| β face (5), (6) | A | | |
| | | A265(DA)\|\|A428(DA); 4w2g<br>(3.9 Å, 15.2°, 20.8°,24.9°,140.2°) | A14(CX)\|\| G34(CW); 4v5d<br>(3.9 Å, 6.5°, 25.3°, 30.5°, 153.1°) |
| | G | | |
| | | | G94(B)\|\|G138(B); 2h0w<br>(3.7 Å, 11.7°, 20.6°, 24.2°, 96.3°) |

**Table S112**. Examples (i.e. identities of bases involved in stacking and corresponding PDB codes) of consecutive (black) and non-consecutive (grey) 5||5, 6||56 β||β *trans* purine||purine stacks. The geometrical parameters ($\vec{d}_{ab}$, $\theta_{ab}$, $\tau_a$, $\tau_b$ and $\sigma_{ab}$) averaged for each ring:ring stacking contact (i.e. average parameters from 5||5, 6||5 and 6||6 ring:ring contacts) for each example stack are provided in parentheses respectively.

| | | β face (5), (56) | |
|---|---|---|---|
| | | A | G |
| β face (5), (6) | A | A1586(14)\|\|A1587(14); 6gsk<br>(3.8 Å, 15.2, 18.6°, 25.3°, 105.1°) | A2469(1H)\|\|G2470(1H); 5el5<br>(3.8 Å, 11.2°, 21.7°,26.1°, 112.3°) |
| | | A9(A)\|\|A27(A); 2zy6<br>(3.7 Å, 5.1°, 26.0°, 22.9°, 144.3°) | A6(A)\|\|G24(A); 3vrs<br>(3.8 Å, 3.1°, 25.9°, 28.0°, 120.7°) |
| | G | | G1(A)\|\|G2(A); 2cky<br>(3.9 Å, 4.6°, 26.8°, 28.7°, 101.3°) |
| | | | G1947(0)\|\|G1970(0); 3ccl<br>(3.6 Å, 2.0°, 27.4°, 27.0°, 123.4°) |

**Table S113.** Examples (i.e. identities of bases involved in stacking and corresponding PDB codes) of consecutive (black) and non-consecutive (grey) 5||56, 6||6 β||β *trans* purine||purine stacks. The geometrical parameters ($\vec{d}_{ab}$, $\theta_{ab}$, $\tau_a$, $\tau_b$ and $\sigma_{ab}$) averaged for each ring:ring stacking contact (i.e. average parameters from 5||5, 5||6 and 6||6 ring:ring contacts) for each example stack are provided in parentheses respectively.

| | | β face (56), (6) | |
|---|---|---|---|
| | | A | G |
| β face (5), (6) | A | | A26(1K)\|\|G27(1K); 5el6<br>(3.6 Å, 15.6°, 21.7°, 23.4°, 96.1°) |
| | | | A21(C)\|\|G46(C); 5wwr<br>(3.6 Å, 7.2°, 20.9°, 25.3°, 164.1°) |
| | G | | |

**Table S114.** Examples (i.e. identities of bases involved in stacking and corresponding PDB codes) of consecutive (black) and non-consecutive (grey) 5||56, 6||5 β||β *trans* purine||purine stacks. The geometrical parameters ($\vec{d}_{ab}$, $\theta_{ab}$, $\tau_a$, $\tau_b$ and $\sigma_{ab}$) averaged for each ring:ring stacking contact (i.e. average parameters from 5||5, 5||6 and 6||5 ring:ring contacts) for each example stack are provided in parentheses respectively.

| | | β face (56), (5) | |
|---|---|---|---|
| | | A | G |
| β face (5), (6) | A | A5(n)\|\|N6(q);1cvj<br>(4.0 Å, 16.7°, 22.9°, 21.5°, 104.8°) | |
| | | A1429(X)\|\|1603(X);2zjr<br>(3.9 Å, 6.1°, 23.1°, 22.1°, 117.1°) | A2448(0)\|\|G2461(X);1xbp<br>(4.2 Å, 17.0°, 24.5°, 16.1°, 128.5°) |
| | G | | |

**Table S115.** Examples (i.e. identities of bases involved in stacking and corresponding PDB codes) of consecutive (black) and non-consecutive (grey) 5||6, 6||6 β||β *trans* purine||purine stacks. The geometrical parameters ($\vec{d}_{ab}$, $\theta_{ab}$, $\tau_a$, $\tau_b$ and $\sigma_{ab}$) averaged for each ring:ring stacking contact (i.e. average parameters from 5||6 and 6||6 ring:ring contacts) for each example stack are provided in parentheses respectively.

| | | β face (6), (6) | |
|---|---|---|---|
| | | A | G |
| β face (5), (6) | A | | |
| | | A448(A)\|\|A487(A); 1ibm<br>(3.5 Å, 6.9°, 19.2°, 23.6°, 117.5°) | A1239(CA)\|\|G1241(CA); 4u24<br>(3.9 Å, 15.5°, 30.7°,26.4°, 104.2°) |
| | G | | |
| | | | G1699(BB)\|\|G1763(BB); 4v64<br>(4.1 Å, 15.1°, 31.4°, 31.0°, 112.1°) |

**Table S116.** Examples (i.e. identities of bases involved in stacking and corresponding PDB codes) of consecutive (black) and non-consecutive (grey) 6||56 β||β *trans* purine||purine stacks. The geometrical parameters ($\vec{d}_{ab}$, $\theta_{ab}$, $\tau_a$, $\tau_b$ and $\sigma_{ab}$) averaged for each ring:ring stacking contact (i.e. average parameters from 6||5 and 6||6 ring:ring contacts) for each example stack are provided in parentheses respectively.

| | | β face (56) | |
|---|---|---|---|
| | | A | G |
| β face (6) | A | | A2469(YA)\|\|G2470(YA);6buw (3.9 Å, 9.2°, 21.9°, 20.8°, 111.4°) |
| | | | A815(A)\|\|G1529(A); 5wns (3.6 Å, 9.1°, 17.4°, 21.7°, 111.5°) |
| | G | | |

**Table S117.** Examples (i.e. identities of bases involved in stacking and corresponding PDB codes) of consecutive (black) and non-consecutive (grey) 5||5, 6||6 β||β *trans* purine||purine stacks. The geometrical parameters ($\vec{d}_{ab}$, $\theta_{ab}$, $\tau_a$, $\tau_b$ and $\sigma_{ab}$) averaged for each ring:ring stacking contact (i.e. average parameters from 5||5 and 6||6 ring:ring contacts) for each example stack are provided in parentheses respectively.

| | | β face (5), (6) | |
|---|---|---|---|
| | | A | G |
| β face (5), (6) | A | | |
| | | A1092(1a)\|\|A1183(1a);5wit (3.8 Å, 18.7°, 33.0°, 15.5°, 164.1°) | G570(a)\|\|A873(a);4jv5 (3.8 Å, 10.1°, 36.0°, 28.0°, 132.7°) |
| | G | | |
| | | | G1947(0)\|\|G1970(0);1qvf (3.9 Å, 9.9°, 25.2°, 32.4°, 117.7°) |

**Table S118.** Examples (i.e. identities of bases involved in stacking and corresponding PDB codes) of consecutive (black) and non-consecutive (grey) 5||6, 6||5 β||β *trans* purine||purine stacks. The geometrical parameters ($\vec{d}_{ab}$, $\theta_{ab}$, $\tau_a$, $\tau_b$ and $\sigma_{ab}$) averaged for each ring:ring stacking contact (i.e. average parameters from 5||6 and 6||5 ring:ring contacts) for each example stack are provided in parentheses respectively.

| | | β face (6), (5) | |
|---|---|---|---|
| | | A | G |
| β face (6), (5) | A | | |
| | G | | |

**Table S119.** Examples (i.e. identities of bases involved in stacking and corresponding PDB codes) of consecutive (black) and non-consecutive (grey) 5||56 β||β *trans* purine||purine stacks. The geometrical parameters ($\vec{d}_{ab}$, $\theta_{ab}$, $\tau_a$, $\tau_b$ and $\sigma_{ab}$) averaged for each ring:ring stacking contact (i.e. average parameters from 5||5 and 5||6 ring:ring contacts) for each example stack are provided in parentheses respectively.

| | | β face (56) | |
|---|---|---|---|
| | | A | G |
| β face (5) | A | | |
| | | A42(B)‖A71(B);3skw<br>(3.9 Å, 22.8°, 21.6°, 16.6°, 144.6°) | A1502(CA)‖G1504(CA); 4u24<br>(3.5 Å, 20.5°, 18.4°, 30.5°, 112.3°) |
| | G | | |
| | | | G2249(1)‖G2272(1); 4u51<br>(3.5 Å, 13.3°, 25.3°, 24.1°, 109.9°) |

**Table S120.** Examples (i.e. identities of bases involved in stacking and corresponding PDB codes) of consecutive (black) and non-consecutive (grey) 5||5, 6||5 β||β *trans* purine||purine stacks. The geometrical parameters ($\vec{d}_{ab}$, $\theta_{ab}$, $\tau_a$, $\tau_b$ and $\sigma_{ab}$) averaged for each ring:ring stacking contact (i.e. average parameters from 5||5 and 6||5 ring:ring contacts) for each example stack are provided in parentheses respectively.

| | | β face (5), (5) | |
|---|---|---|---|
| | | A | G |
| β face (5), (6) | A | | A933(DA)‖G934(DA); 4v8b (3.6 Å, 6.5°, 28.2°, 24.6°, 100.1°) |
| | | | A872(AA)‖G874(AA); 4v64 (3.5 Å, 19.3°, 33.1°, 19.0°, 123.4°) |
| | G | | |

**Table S121.** Examples (i.e. identities of bases involved in stacking and corresponding PDB codes) of consecutive (black) and non-consecutive (grey) 6||6 β||β *trans* purine||purine stacks. The geometrical parameters ($\vec{d}_{ab}$, $\theta_{ab}$, $\tau_a$, $\tau_b$ and $\sigma_{ab}$) for each example stack are provided in parentheses respectively.

| | | β face (6) | |
|---|---|---|---|
| | | A | G |
| β face (6) | A | | |
| | | A766(0)‖A897(0);1vqn (4.2 Å, 11.0°, 24.9°, 35.8°, 108.5°) | A196(BA)‖G805(BA); 4u24 (3.8 Å , 18.4°, 32.0°, 20.1°, 163.3°) |
| | G | | |

**Table S122.** Examples (i.e. identities of bases involved in stacking and corresponding PDB codes) of consecutive (black) and non-consecutive (grey) 5||6 β||β *trans* purine||purine stacks. The geometrical parameters ($\vec{d}_{ab}$, $\theta_{ab}$, $\tau_a$, $\tau_b$ and $\sigma_{ab}$) for each example stack are provided in parentheses respectively.

| | | β face (6) | |
|---|---|---|---|
| | | A | G |
| β face (5) | A | | A25(E)‖G26(F); 1q2r (4.0 Å, 2.3°, 31.3°, 32.9°, 109.3°) |
| | | | A1875(A)‖G1877(A); 1q81 (3.7 Å, 4.9°, 21.5°, 24.9°, 103.6°) |
| | G | | |

**Table S123.** Examples (i.e. identities of bases involved in stacking and corresponding PDB codes) of consecutive (black) and non-consecutive (grey) 6||5 β||β *trans* purine||purine stacks. The geometrical parameters ($\vec{d}_{ab}$, $\theta_{ab}$, $\tau_a$, $\tau_b$ and $\sigma_{ab}$) for each example stack are provided in parentheses respectively.

| | | β face (5) | |
|---|---|---|---|
| | | A | G |
| β face (6) | A | A1359(BA)\|\|A1360(BA); 4v8b (4.0 Å, 16.8°, 37.8°, 21.4°, 107.1°) | A2469(DA)\|\|G2470(DA);4v8b (3.3 Å. 16.6°, 14.9°, 28.0°, 101.5°) |
| | | A1272(DA)\|\|A1618(DA); 4u24 (3.7 Å, 13.2°, 38.3°, 32.9°, 156.1°) | A397(A)\|\|G548(A);5wns (3.9 Å, 2.6°, 38.2°, 37.8°, 126.5°) |
| | G | | G401(B)\|\|G402(B); 1zl3 (3.5 Å, 11.0°, 20.4°, 18.9°, 113.9°) |
| | | | G129A(A)\|\|G190G(A); 5wns (3.7 Å, 5.1°, 25.2°, 30.1°, 110.0°) |

**Table S124.** Examples (i.e. identities of bases involved in stacking and corresponding PDB codes) of consecutive (black) and non-consecutive (grey) 5||5 β||β *trans* purine||purine stacks. The geometrical parameters ($\vec{d}_{ab}$, $\theta_{ab}$, $\tau_a$, $\tau_b$ and $\sigma_{ab}$) for each example stack are provided in parentheses respectively.

| | | β face (5) | |
|---|---|---|---|
| | | A | G |
| β face (5) | A | A2169(1H)\|\|A21704(1H);5wqr (3.6 Å, 9.9°, 28.2°, 34.9°, 91.4°) | A26(1L)\|\|G27(1L);4wro (3.4 Å, 18.0, 28.1°, 10.2°, 101.2°) |
| | | A1074(A)\|\|A1164(A);4gkk (4.1 Å, 22.9°, 13.1°, 32.7°, 165.0°) | A1005(A)\|\|G1026(A);1xnr (4.3 Å, 21.2°, 8.4°, 26.8°, 145.5°) |
| | G | | G81(DB)\|\|G82(DB);4v5a (3.7 Å, 22.5°, 29.3°, 35.5°, 96.8°) |
| | | | G1947(0)\|\|G1970(0);3cxc (3.6 Å, 12.8°, 18.3°, 27.9°, 115.0°) |

**Table S125.** Percent occurrence frequency of three most dominant topologies of purine||pyrimidine stacks.

| **Topology** | **Percent occurrence** |
|---|---|
| 5\|\|6, 6\|\|6 α\|\|α cis | 36.0% |
| 5\|\|6 β\|\|β cis | 23.6% |
| 6\|\|6 α\|\|α cis | 21.3% |

**Table S126.** Examples (i.e. identities of bases involved in stacking and corresponding PDB codes) of consecutive (black) and non-consecutive (grey) 5||6, 6||6 α||α *cis* purine||pyrimidine stacks The geometrical parameters ($\vec{d}_{ab}$, $\theta_{ab}$, $\tau_a$, $\tau_b$ and $\sigma_{ab}$) averaged for each ring:ring stacking contact (i.e. average parameters from 5||6 and6||6 ring:ring contacts) for each example stack are provided in parentheses respectively.

<table>
<tr><td colspan="2" rowspan="2"></td><td colspan="2">α face (6)</td></tr>
<tr><td>C</td><td>U</td></tr>
<tr><td rowspan="4">α face (5),(6)</td><td rowspan="2">A</td><td>A14(A)||C15(A); 3gca<br>(3.5 Å, 7.9°, 18.7°, 16.9°, 39.4°)</td><td>A6(A)||U7(A); 4kqy<br>(4.0 Å, 14.2°, 15.7°, 25.5°, 16.4°)</td></tr>
<tr><td>A44(A)||C46(A);4wfl<br>(4.0 Å, 2.0°, 31.2°, 29.5°, 6.3°)</td><td>A2020(DA)||U2022(DA); 1vy6<br>(3.5 Å, 10.1°, 24.8°, 19.2°, 77.7°)</td></tr>
<tr><td rowspan="2">G</td><td>G28(A)||C29(A); 4kqy<br>(3.8 Å,11.7°, 22.9°, 16.4°, 49.5°)</td><td>G2(3)||U3(3); 1pgl<br>(3.7 Å, 4.8°, 24.9°, 20.5°, 26.8°)</td></tr>
<tr><td>G616(BA)||C618(BA); 1vy6<br>(3.9 Å, 6.0°, 21.6°,23.7°, 33.9°)</td><td>G1182 (AA)|| U1159 (AA); 1vy6<br>(4.2 Å, 12.9°, 17.5°, 18.5°, 69.2°)</td></tr>
</table>

**Table S127.** Examples (i.e. identities of bases involved in stacking and corresponding PDB codes) of consecutive (black) and non-consecutive (grey) 6||6 α||α *cis* purine||pyrimidine stacks. The geometrical parameters ($\vec{d}_{ab}$, $\theta_{ab}$, $\tau_a$, $\tau_b$ and $\sigma_{ab}$) for each example stack are provided in parentheses respectively.

<table>
<tr><td colspan="2" rowspan="2"></td><td colspan="2">α face (6)</td></tr>
<tr><td>C</td><td>U</td></tr>
<tr><td rowspan="4">α face (6)</td><td rowspan="2">A</td><td>A655(A)||C656(A); 2uxd<br>(3.4 Å, 12.4°, 13.3°, 7.6°, 22.4°)</td><td>A323(6)||U324(6); 4u3n<br>(3.9 Å 19.5°, 15.1°, 34.6°, 18.6°)</td></tr>
<tr><td>A315(A)||C330(A); 2uxd<br>(4.2 Å, 7.7°, 24.9°, 32.0°, 32.0°)</td><td>A1021(14)||U1023(14);4wqr<br>(3.4 Å, 7.3°, 14.2°, 9.1°, 6.0°)</td></tr>
<tr><td rowspan="2">G</td><td>G644(A)||C645(A; 2uxd<br>(3.4 Å, 7.5°, 21.5°, 14.4°, 37.2°)</td><td>G1767(0)||U1768(0); 1ond<br>(3.7 Å, 7.6°, 27.5°, 32.3°, 29.4°)</td></tr>
<tr><td>C905(0)||G1354(0); 3g6e<br>(4.0 Å, 9.9°, 28.0°, 35.2°, 18.4°)</td><td>G71(P)||U136(P); 3g8s<br>(3.7 Å, 5.4°, 24.9°, 25.7°, 18.1°)</td></tr>
</table>

**Table S128.** Examples (i.e. identities of bases involved in stacking and corresponding PDB codes) of consecutive (black) and non-consecutive (grey) 5||6 α||α *cis* purine||pyrimidine stacks. . The geometrical parameters ($\vec{d}_{ab}$, $\theta_{ab}$, $\tau_a$, $\tau_b$ and $\sigma_{ab}$) for each example stack are provided in parentheses respectively.

| | | α face (6) | |
|---|---|---|---|
| | | C | U |
| α face (5) | A | A1922(1)\|\|C1923(1);5obm (3.6 Å ,6.1°, 29.6°, 24.5°, 73.2°) | A980(1)\|\|U981(1);5tga (3.4 Å, 18.6°, 25.6°, 13.0°, 12.4°) |
| | | A440(AA)\|\|C442(AA); 4v8b (3.7 Å, 20.2°, 11.3°, 30.5°, 70.4) | A222(DA)\|\|U224(DA); 4u24 (3.2 Å, 17.9°, 9.0°, 24.6°, 70.4°) |
| | G | G1131(A)\|C1132(A);2uuc (4.5 Å, 20.4°, 35.5°, 31.6°, 5.6°) | G1198(AA)\|\|U1199(AA);4v7m (4.0 Å, 22.9°, 23.2°, 23.4°, 18.3°) |
| | | G224(0)\|\|C226(0); 2aar (3.9 Å, 9.9°, 18.8°, 28.2°, 79.9°) | G10(A)\|\|U13(A); 4y1m (4.1 Å, 9.6°, 33.6°, 24.3°, 57.7°) |

**Table S129.** Examples (i.e. identities of bases involved in stacking and corresponding PDB codes) of consecutive (black) and non-consecutive (grey) 5||6, 6||6 α||α *trans* purine||pyrimidine stacks. The geometrical parameters ($\vec{d}_{ab}$, $\theta_{ab}$, $\tau_a$, $\tau_b$ and $\sigma_{ab}$) averaged for each ring:ring stacking contact (i.e. average parameters from 5||6 and 6||6 ring:ring contacts) for each example stack are provided in parentheses respectively.

| | | α face (6) | |
|---|---|---|---|
| | | C | U |
| α face (5), (6) | A | A3135(YA)\|\|C3136(YA);6buh (4.0 Å, 4.7°, 25.2°, 23.7°, 92.9°) | A73(BB)\|\|U74(BB);4v9l (3.9 Å, 16.4°, 15.7°, 23.0°, 95.5°) |
| | | A58(C)\|\|C61(C); 5wwr (3.8 Å, 8.0°,21.6°, 19.0°, 137.0°) | A788(DA)\|\|U464(DA);1vy6 (3.7 Å, 8.5°, 16.7°, 20.7°, 116.8°) |
| | G | | G2112(BA)\|\|U2113(BA);4v9c (3.5 Å, 10.3°, 16.7°, 25.3°, 90.9°) |
| | | G838(A)\|\|G840(A); 4lf4 (4.0 Å,14.7°, 17.1°, 27.1°, 111.6°) | G517(AA)\|\|U531(AA);4u1u (3.8 Å, 16.4°, 33.9°, 21.5°, 105.0°) |

**Table S130.** Examples (i.e. identities of bases involved in stacking and corresponding PDB codes) of consecutive (black) and non-consecutive (grey) 6||6 α||α *trans* purine||pyrimidine stacks. The geometrical parameters ($\vec{d}_{ab}$, $\theta_{ab}$, $\tau_a$, $\tau_b$ and $\sigma_{ab}$) for each example stack are provided in parentheses respectively.

| | | α face (6) | |
|---|---|---|---|
| | | C | U |
| α face (6) | A | | |
| | | A1355(x)\|\|C1358(x);5dm7<br>(3.6 Å, 18.9°,22.2°,37.6°, 121.3°) | A2956(5)\|\|U2141(5);5dc3<br>(4.2 Å, 4.8°, 35.5°, 30.7°, 171.9°) |
| | G | | |
| | | G1378(A)\|\|C2747(A);1kc8<br>(3.5 Å, 14.8°, 10.5°, 11.6°,179.4°) | G2073(0)\|\|U2607(0);1kqs<br>(3.4 Å, 17.6°, 22.6°, 10.1°, 145.7°) |

**Table S131.** Examples (i.e. identities of bases involved in stacking and corresponding PDB codes) of consecutive (black) and non-consecutive (grey) 5||6 α||α *trans* purine||pyrimidine stacks. The geometrical parameters ($\vec{d}_{ab}$, $\theta_{ab}$, $\tau_a$, $\tau_b$ and $\sigma_{ab}$) for each example stack are provided in parentheses respectively.

| | | α face (6) | |
|---|---|---|---|
| | | C | U |
| α face (5) | A | A205(AA)\|\|C206(AA); 4ybb<br>(4.0 Å,4.6°, 27.9°, 31.7°, 101.4°) | A59(3k)\|\|U60(3k); 5el6<br>(3.9 Å, 5.1°, 22.1°, 23.1°, 110.5°) |
| | | A1239(A)\|\|C124=98(A);2uxd<br>(4.1 Å, 12.0°, 34.8°, 25.1°,136.1°) | A1776(X)\|\|U1778(X); 4ioa<br>(3.7 Å, 18.7°, 14.9°, 28.4°, 104.3°) |
| | G | G1026(2a)\|\|C1027(2a); 6fkr<br>(4.1 Å ,18.9°, 15.7°, 33.9°, 108.9°) | G834(6)\|\|U835(6); 5ndv<br>(3.4 Å, 19.5°, 11.6°, 27.7°, 96.5°) |
| | | G336(2)\|\|338(2); 4u53<br>(4.2 Å, 21.3°, 32.8°, 14.7°, 114.3°) | G517(CA)\|\|U531(CA); 4v67<br>(3.8 Å, 9.4°, 19.3°, 27.8°, 129.7°) |

**Table S132.** Examples (i.e. identities of bases involved in stacking and corresponding PDB codes) of consecutive (black) and non-consecutive (grey) 5||6, 6||6 α||β *cis* purine||pyrimidine stacks. The geometrical parameters ($\vec{d}_{ab}$, $\theta_{ab}$, $\tau_a$, $\tau_b$ and $\sigma_{ab}$) averaged for each ring:ring stacking contact (i.e. average parameters from 5||6 and 6||6 ring:ring contacts) for each example stack are provided in parentheses respectively.

| | | β face (6) | |
|---|---|---|---|
| | | C | U |
| α face (5) ,(6) | A | A34(W)‖C35(W);2gtt<br>(4.2 Å, 18.4°, 31.0°, 17.1°, 10.7°) | A161(CA)‖U162(CA);4v9p<br>(4.1 Å, 2.9°, 27.5°, 30.0°, 75.0°) |
| | | A665(BA)‖C732(BA); 4v9o<br>(3.9 Å, 6.3°, 25.1°, 25.3°, 4.2°) | A9(1w)‖U45(1w);4y4p<br>(4.0 Å, 12.2°, 18.1°, 23.2°, 11.1°) |
| | G | G892(A)‖C893(A);1q82<br>(3.8 Å, 19.0°, 19.6°, 25.1°, 75.7°) | G612(DA)‖U613(DA);4v87<br>(3.9 Å, 5.6°, 23.2°, 21.8°, 19.1°) |
| | | G58(B)|C60(B);5xbl<br>(3.9 Å, 13.8°, 24.3°, 24.6°, 4.8°) | G40(B)‖U67(B);3b4c<br>(4.0 Å, 11.5°, 25,0°, 24.6°, 0.9°) |

**Table S133.** Examples (i.e. identities of bases involved in stacking and corresponding PDB codes) of consecutive (black) and non-consecutive (grey) 6||6 α||β *cis* purine||pyrimidine stacks. The geometrical parameters ($\vec{d}_{ab}$, $\theta_{ab}$, $\tau_a$, $\tau_b$ and $\sigma_{ab}$) for each example stack are provided in parentheses respectively.

| | | β face (6) | |
|---|---|---|---|
| | | C | U |
| α face (6) | A | A70(X)‖C71(X); 2gtt<br>(3.6 Å, 16.1°, 18.4°, 13.1°, 1.0°) | A2845(1)‖U2846(1); 4u55<br>(3.4 Å, 1.2°, 19.1°, 19.7°, 30.0°) |
| | | C133(X)‖A139(X);2zjr<br>(4.0 Å, 18.4°, 35.8°,25.1°,15.2°) | A9(x)‖U45(x); 4w2e<br>(3.7 Å, 12.0°, 26.1°, 38.0°, 15.5°) |
| | G | C2825(0)‖G2826(0); 3ccv<br>(3.7 Å, 13.1°, 17.1°, 26.3°, 58.1) | G98(BB)‖U99(BB); 4v64<br>(3.9 Å, 15.9°, 19.1°, 33.3°, 11.9°) |
| | | C72(9)‖G110(9); 3ccv<br>(4.1 Å, 8.1°, 35.0°, 38.9°, 15.6°) | U2074(DA)‖G2436(DA); 1vy7<br>(4.1 Å, 7.5°, 30.5°, 36.2°, 31.0°) |

**Table S134.** Examples (i.e. identities of bases involved in stacking and corresponding PDB codes) of consecutive (black) and non-consecutive (grey) 5||6 α||β*cis* purine||pyrimidine stacks. The geometrical parameters ($\vec{d}_{ab}$, $\theta_{ab}$, $\tau_a$, $\tau_b$ and $\sigma_{ab}$) for each example stack are provided in parentheses respectively.

| | | β face (6) | |
|---|---|---|---|
| | | C | U |
| α face (5) | A | A1157(1G)\|\|C1158(1G);6gsl (4.4 Å, 22.4°, 32.3°, 21.2°, 47.9°) | A411(X)\|\|U412(X);5nrg (3.7 Å, 18.1°, 36.0°, 20.1°, 49.2°) |
| | | A2176(A)\|\|C2197(A); 5ml7 (4.0 Å, 20.0°, 15.1°, 35.2°, 87.1°) | A559(CA)\|\|U561(CA); 4v8b (4.2 Å, 22.4°, 33.1°, 14.2°,32.8°) |
| | G | G275(A)\|\|C276(A);2a64 (4.2 Å, 21.1°, 20.8°,29.1°, 43.1°) | G1084(A)\|\|U1085(A);5wns (3.7 Å, 16.7°, 18.8°, 17.2°, 76.1°) |
| | | - | G60(DA)\|\|U62(DA); 4u24 (3.4 Å, 8.7°, 16.4°, 15.4°, 72.4°) |

**Table S135.** Examples (i.e. identities of bases involved in stacking and corresponding PDB codes) of consecutive (black) and non-consecutive (grey) 5||6, 6||6 α||β *trans* purine||pyrimidine stacks. The geometrical parameters ($\vec{d}_{ab}$, $\theta_{ab}$, $\tau_a$, $\tau_b$ and $\sigma_{ab}$) averaged for each ring:ring stacking contact (i.e. average parameters from 5||6 and 6||6 ring:ring contacts) for each example stack are provided in parentheses respectively.

| | | β face (6) | |
|---|---|---|---|
| | | C | U |
| α face (5), (6) | A | A3114(1)\|\|C3115(1); 5on6 (4.0 Å, 21.9°, 14.8°, 16.8°, 91.6°) | A3(B)\|\|U(B); 6dpm (4.0 Å, 17.7°, 14.8°, 17.7°, 101.7°) |
| | | A105(A)\|\|C261(A); 4faq (4.1 Å, 7.3°, 37.4°, 32.8°, 162.8°) | A225(a)\|\|U247(a); 1gid (3.8 Å, 2.8°, 26.0°, 28.0°, 130.7°) |
| | G | G553(A2)\|\|C554(A2);4v88 (3.6 Å, 21.7°, 21.4°, 17.7°, 99.8°) | G1059(DA)\|\|U1060(DA); 4v9l (3.9 Å, 14.5°, 18.1°, 16.7°, 103.6°) |
| | | G1613(YA)\|\|C1617(YA); 5j30 (3.9 Å, 10.4°, 21.5°, 27.4°, 126.8°) | G2816(5)\|\|U2869(5);6u6f (3.8 Å, 13.3°, 20.7°, 22.8°, 106°) |

**Table S136.** Examples (i.e. identities of bases involved in stacking and corresponding PDB codes) of consecutive (black) and non-consecutive (grey) 6||6 α||β *trans* purine||pyrimidine stacks. The geometrical parameters ($\vec{d}_{ab}$, $\theta_{ab}$, $\tau_a$, $\tau_b$ and $\sigma_{ab}$) for each example stack are provided in parentheses respectively.

| | | β face (6) | |
|---|---|---|---|
| | | C | U |
| α face (6) | A | | |
| | | C2197(5)\|\|A2242(5); 4u3u (3.6 Å, 12.0°, 24.5°, 15.8°, 157.6°) | U768(A)\|\|A895(A); 1q86 (4.0 Å, 13.5°, 24.7°, 35.2°, 121.9°) |
| | G | | U-3(B)\|\|G-4(B); 4far (3.9 Å, 1.2°, 30.8°, 29.6°, 111.3°) |
| | | G1613(DA)\|\|C1617(DA); 4u24 (3.3 Å, 7.1°, 10.5°, 17.6°, 133.4°) | G2761(5)\|\|U2795(5); 4u56 (4.1 Å, 18.5°, 24.0°, 19.1°, 177.0°) |

.

**Table S137.** Examples (i.e. identities of bases involved in stacking and corresponding PDB codes) of consecutive (black) and non-consecutive (grey) 5||6 α||β *trans* purine||pyrimidine stacks. The geometrical parameters ($\vec{d}_{ab}$, $\theta_{ab}$, $\tau_a$, $\tau_b$ and $\sigma_{ab}$) for each example stack are provided in parentheses respectively.

| | | β face (6) | |
|---|---|---|---|
| | | C | U |
| α face (5) | A | | A1350(5)\|\|U1351(5); 5ndv (4.4 Å, 12.8°, 16.5°, 17.3°, 94.9°) |
| | | A105(A)\|\|C261(A); 4e8q (4.5 Å, 2.4°, 38.2°, 39.7°, 162.2°) | A84(BA)\|\|U99(BA); 4v8b (4.4 Å, 10.7°, 23.4°, 33.6°, 151.2) |
| | G | | G1059(1A)\|\|U1060(1A); 5doy (3.7 Å, 14.3°, 11.3°, 16.6°, 101.5°) |
| | | G64(AA)\|\|C99(AA); 4v64 (4.1 Å, 5.0°, 39.0°, 35.3°, 148.2°) | G3022(B)\|\|U3055(B); 1q81 (3.5 Å, 22.8°, 30.8°, 9.3°, 143.0°) |

**Table 138.** Examples (i.e. identities of bases involved in stacking and corresponding PDB codes) of consecutive (black) and non-consecutive (grey) 5||6, 6||6 β||α *cis* purine||pyrimidine stacks. The geometrical parameters ($\vec{d}_{ab}$, $\theta_{ab}$, $\tau_a$, $\tau_b$ and $\sigma_{ab}$) averaged for each ring:ring stacking contact (i.e. average parameters from 5||6 and 6||6 ring:ring contacts) for each example stack are provided in parentheses respectively.

| | | α face (6) | |
|---|---|---|---|
| | | C | U |
| β face (5), (6) | A | A71(R)\|\|C72(R);4yb1 (3.8 Å,8.9°, 21.4°, 22.2°,56.7°) | A80(a)\|\|U81(a);4yb0 (4.0 Å, 3.7°, 28.0°, 28.9°, 69.7°) |
| | | A866(DA)\|\|C914(DA);4v9n (4.4 Å, 13.7°, 29.0°, 36.9°, 7.0°) | A1524(5)\|\|U1607(5);4u3u (4.1 Å, 15.1°, 34.5°, 27.3°, 3.7°) |
| | G | G892(0)\|\|C893(0);1yj9 (3.8 Å, 22.5°,22.1°, 25.2°, 77.3°) | G1678(ba)\|\|U1679(ba);1vy6 (4.0 Å, 4.9°, 32.1°, 30.4°, 62.1°) |
| | | G1059(0)\|\|C1127(0);5xbl (3.9 Å, 18.3°,15.8°,20.1°, 72.3°) | G2997(1)\|\|U3396(1);4u3u (3.7 Å, 4.5°, 20.7°, 19.2°, 62.8°) |

**Table S139.** Examples (i.e. identities of bases involved in stacking and corresponding PDB codes) of consecutive (black) and non-consecutive (grey) 6||6 β||α *cis* purine||pyrimidine stacks. The geometrical parameters ($\vec{d}_{ab}$, $\theta_{ab}$, $\tau_a$, $\tau_b$ and $\sigma_{ab}$) for each example stack are provided in parentheses respectively.

| | | α face (6) | |
|---|---|---|---|
| | | C | U |
| β face (6) | A | C1054(x)\|\|A1055(x);2zjq (3.9 Å, 10.7°, 25.2°, 15.0°, 2.6°) | A1062(6)\|\|U1063(6);5ndv (3.6 Å, 19.5°, 17.1°, 6.2°, 80.1°) |
| | | A1859(1)\|\|C1872(1);5dat (4.2 Å, 11.4°. 33.7°, 34.6°, 14.7°) | A247(0)\|\|U265(0);1yjn (3.5 Å, 4.8°, 19.1°, 23.7°, 62.8°) |
| | G | G101(1)\|\|C102(1);4u50 (4.5 Å, 22.0°, 26.4°, 6.6°, 63.9°) | G2782(X)\|\|U2783(X);5jvg (3.8 Å, 7.7°, 23.2°, 17.3°, 69.1°) |
| | | G1544(5)\|\|C1550(5);4u50 (4.0 Å, 7.8°, 38.2°, 30.5°, 33.9°) | G820(0)\|\|U1831(0);3cc2 (3.7 Å, 15.9°, 33.4°, 17.7°, 18.2°) |

**Table S140.** Examples (i.e. identities of bases involved in stacking and corresponding PDB codes) of consecutive (black) and non-consecutive (grey) 5||6 β||α *cis* purine||pyrimidine stacks. The geometrical parameters ($\vec{d}_{ab}$, $\theta_{ab}$, $\tau_a$, $\tau_b$ and $\sigma_{ab}$) for each example stack are provided in parentheses respectively.

| | | α face (6) | |
|---|---|---|---|
| | | C | U |
| β face (5) | A | A2768(0)\|\|C2769(0);3ccl (3.6 Å,10.6°, 17.7°,10.6°, 25.4°) | A1062(6)\|\|U1063(6);5obm (4.0 Å, 13.9°, 17.2°, 9.8°, 43.0°) |
| | | A912(2A)\|\|C959(2A);5hcr (4.3 Å, 18.0°, 37.7°, 30.3°, 16.4°) | A1346(CA)\|\|U1348(CA);4u24 (3.5 Å, 8.5°, 24.2°, 22.5°, 64.0°) |
| | G | G177(CA)\|\|C178(CA);4v7t (4.1 Å, 18.2°, 39.5.0°, 34.3°, 31.1°) | G1724(2A)\|\|U1725(2A); 5hcq (3.9 Å, 11.9°, 22.7°, 32.2°, 70.0°) |
| | | G2345(DA)\|\|C2347(DA); 4u24 (3.5 Å, 16.6°, 32.4°, 17.9°, 44.3°) | G39(a)\|\|U498(a);1xmo (4.5 Å, 20.1°, 21.4°, 36.4°, 40.4°) |

.

**Table S141.** Examples (i.e. identities of bases involved in stacking and corresponding PDB codes) of consecutive (black) and non-consecutive (grey) 5||6, 6||6 β||α *trans* purine||pyrimidine stacks. The geometrical parameters ($\vec{d}_{ab}$, $\theta_{ab}$, $\tau_a$, $\tau_b$ and $\sigma_{ab}$) averaged for each ring:ring stacking contact (i.e. average parameters from 5||6 and 6||6 ring:ring contacts) for each example stack are provided in parentheses respectively.

| | | α face (6) | |
|---|---|---|---|
| | | C | U |
| β face (5), (6) | A | A468(CA)\|\|C469(CA);4v7s (3.9 Å, 13.8°, 22.2°, 32.8°, 102.1°) | A1241(BB)\|\|U1242(BB);4v64 (3.7 Å, 5.4°, 21.3°, 25.4°, 102.5°) |
| | | A21(F)\|\|C48(F);4yco (3.6 Å, 6.6°, 23.0°, 24.9°, 126.9°) | A21(B)\|\|U48(B);4ycp (3.6 Å, 6.9°, 22.4°, 27.4°,136.2°) |
| | G | G27(1L)\|\|C28(1L);4wsm (3.5 Å, 3.3°, 20.9°, 19.5°, 103.5°) | |
| | | G2482(0)\|\|C2536(0);3ccj (3.5 Å, 3.5°, 19.1°, 17.8°, 115.7°) | G170(B)\|\|U173(B);3bo2 (3.8 Å, 4.4°, 26.2°, 22.3°,118.3°) |

**Table S142.** Examples (i.e. identities of bases involved in stacking and corresponding PDB codes) of consecutive (black) and non-consecutive (grey) 6||6 β||α *trans* purine||pyrimidine stacks. The geometrical parameters ($\vec{d}_{ab}$, $\theta_{ab}$, $\tau_a$, $\tau_b$ and $\sigma_{ab}$) for each example stack are provided in parentheses respectively.

| | | α face (6) | |
|---|---|---|---|
| | | C | U |
| β face (6) | A | A718(DB)\|\|C719(DB); 4v64<br>(3.9 Å, 19.0°, 24.0°, 37.3°, 110.9°) | A1230(6)\|\|U1231(6); 5tga<br>(4.3 Å, 17.4°, 24.3°,31.9°, 107.8°) |
| | | A65(9)\|\|C113(9);3ccj<br>(3.4 Å, 13.1°, 18.7°, 12.0°, 171.9°) | A2727(0)\|\|U2756(0); 2qex<br>(3.5 Å, 5.1°, 32.0°, 30.4°, 119.2°) |
| | G | G1036(1a)\|\|C1037(1a); 6cae<br>(3.7 Å, 13.2°, 23.2°, 21.5°, 101.7°) | G59(D)\|\|U60(D); 2csx<br>(3.5 Å, 12.4°, 16.1°, 14.1°, 104.7°) |
| | | G1138(A)\|\|C1140(A); 2uub<br>(3.9 Å, 21.4°, 31.4°, 32.3°, 121.5°) | G1119(0)\|\|U1244(0); 1jj2<br>(3.5 Å, 9.4°, 19.4°, 18.5°, 123.2°) |

**Table S143.** Examples (i.e. identities of bases involved in stacking and corresponding PDB codes) of consecutive (black) and non-consecutive (grey) 5||6 β||α *trans* purine||pyrimidine stacks. The geometrical parameters ($\vec{d}_{ab}$, $\theta_{ab}$, $\tau_a$, $\tau_b$ and $\sigma_{ab}$) for each example stack are provided in parentheses respectively.

| | | α face (6) | |
|---|---|---|---|
| | | C | U |
| β face (5) | A | | |
| | | A1342(14)\|\|C1345(14); 4wsm<br>(4.0 Å, 6.4°, 33.1°, 37.6°, 161.8°) | A2521(0)\|\|U2498(0);1xbp<br>(3.2 Å, 11.1°, 15.5°, 7.1°, 108.8°) |
| | G | | |
| | | G2482(A)\|\|C2536(A);1ffz<br>(3.2 Å, 11.1°, 10.9°,6.2°, 123.5°) | G551(2)\|\|U582(2); 4u55<br>(3.8 Å, 13.8°, 21.9°, 9.9°, 127.2°) |

**Table S144.** Examples (i.e. identities of bases involved in stacking and corresponding PDB codes) of consecutive (black) and non-consecutive (grey) 5||6, 6||6 β||β *cis* purine||pyrimidine stacks. The geometrical parameters ($\vec{d}_{ab}$, $\theta_{ab}$, $\tau_a$, $\tau_b$ and $\sigma_{ab}$) averaged for each ring:ring stacking contact (i.e. average parameters from 5||6 and 6||6 ring:ring contacts) for each example stack are provided in parentheses respectively.

| | | β face (6) | |
|---|---|---|---|
| | | C | U |
| β face (5),(6) | A | A34(X)\|\|C35(X); 2gtt (3.9 Å, 21.8°, 20.8°, 17.4°, 54.3°) | A1504(0)\|\|U1503(0); 1qvg (3.7 Å, 12.7°, 16.9°, 17.4°, 23.0°) |
| | | A243(A)\|\|C245(A); 5wns (3.9 Å, 8.5°, 26.1°, 27.3°, 55.1°) | A9(A)\|\|U63(A); 5fkf (4.0 Å, 7.5°23.6°, 25.3°,26.9° ) |
| | G | G6(E)\|\|C7(E); 3rc8 (4.1 Å, 17.2°, 32.3°, 20.3°. 53.4°) | G1580(BA)\|\|U1581(BA); 4w2h (3.8 Å, 10.8°, 22.3°, 20.2°,55.6°) |
| | | G37(A)\|\|C39(A); 6dnr (3.8 Å, 7.7°, 24.9°, 23.6°, 28.1°) | G976(AA)\|\|U1358(AA); 4v64 (4.0 Å, 11.0°, 32.5°, 26.0°, 68.2°) |

**Table S145.** Examples (i.e. identities of bases involved in stacking and corresponding PDB codes) of consecutive (black) and non-consecutive (grey) 6||6 β||β *cis* purine||pyrimidine stacks. The geometrical parameters ($\vec{d}_{ab}$, $\theta_{ab}$, $\tau_a$, $\tau_b$ and $\sigma_{ab}$) for each example stack are provided in parentheses respectively.

| | | β face (6) | |
|---|---|---|---|
| | | C | U |
| β face (6) | A | C967(AA)\|\|A968(AA);4v4h (3.8 Å, 17.4°, 16.7°, 13.6°, 23.7°) | U10(R)\|\|A11(R) ;4fts (4.3 Å, 23.0°, 18.8°, 39.5°, 55.8°) |
| | | A48(4)\|\|C62(4);6hhq (4.0 Å, 11.4°, 27.8°, 38.6°, 24.8°) | A243(CA)\|\|U245(CA); 4u24 (3.4 Å, 6.7°, 28.6°, 26.6°, 48.3°) |
| | G | C2306(BB)\|\|G2307(BB); 4v64 (4.4 Å, 8.0°, 34.1°, 38.4°, 47.5°) | U751(A)\|\|G752(A);4x64 (4.3 Å, 22.5°, 3.3°, 25.8°, 63.5°) |
| | | G33(B)\|\|C60(B);3slq (3.8 Å, 20.6°, 23°, 18.7°, 61.0) | G46(CB)\|\|U55(CB);4v8f (3.9 Å, 7.4°, 37.8°, 34.6°, 40.7°) |

**Table S146.** Examples (i.e. identities of bases involved in stacking and corresponding PDB codes) of consecutive (black) and non-consecutive (grey) 5||6 β||β *cis* purine||pyrimidine stacks. The geometrical parameters ($\vec{d}_{ab}$, $\theta_{ab}$, $\tau_a$, $\tau_b$ and $\sigma_{ab}$) for each example stack are provided in parentheses respectively.

| | | β face (6) | |
|---|---|---|---|
| | | C | U |
| β face (5) | A | A537(B)\|\|C536(B);1g59<br>(3.7 Å, 9.1°, 23.4°, 30.1°, 25.2°) | A1490(B)\|\|U1489(B);2oe5<br>(4.1 Å, 14.3°, 33.5°, 29.3°, 20.1°) |
| | | A2628(5)\|\|C2798(5); 4u55<br>(4.1 Å, 5.7°, 37.7°, 38.3°, 13.4°) | A1(D)\|\|U3(E); 3bo2<br>(4.5 Å, 18.7°, 28.1°, 35.3°, 13.7°) |
| | G | C1725(0)\|\|G1726(0);3g4s<br>(3.9 °, 14.8°, 29.5°, 35.5°, 24.2°) | G31(A)\|\|U30(A);3g4m<br>(4.5 Å, 6.3°, 30.3°, 36.0°, 26.3°) |
| | | G2259(BA)\|\|C2427(BA); 4v8b<br>(3.7 Å, 4.6°, 29.3°, 33.3°, 0.2°) | G976(AA)\|\|U1358(AA); 4u24<br>(4.1 Å, 15.2°, 37.0°, 25.6°, 58.7°) |

**Table S147.** Examples (i.e. identities of bases involved in stacking and corresponding PDB codes) of consecutive (black) and non-consecutive (grey) 5||6, 6||6 β||β *trans* purine||pyrimidine stacks. The geometrical parameters ($\vec{d}_{ab}$, $\theta_{ab}$, $\tau_a$, $\tau_b$ and $\sigma_{ab}$) averaged for each ring:ring stacking contact (i.e. average parameters from 5||6 and 6||6 ring:ring contacts) for each example stack are provided in parentheses respectively.

| | | β face (6) | |
|---|---|---|---|
| | | C | U |
| β face (5) ,(6) | A | A40(CW)\|\|C41(CW);4v8n<br>(3.7 Å, 20.4°, 22.2°, 20.1°, 102.8°) | |
| | | A35(A)\|\|C61(A);2xnz<br>(4.0 Å, 11.2°, 20.7°, 30.1°, 169.8°) | A215(2)\|\|U242(2);4u55<br>(4.2 Å, 11.6°, 29.5°, 19.9°, 168.4°) |
| | G | | |
| | | G1378(A)\|\|C2724(A); 1m1k<br>(3.8 Å, 13.1°, 19.0°, 18.4°, 177.4°) | G535(0)\|\|U2063(0);1vqo<br>(3.9 Å, 5.7°, 30.3°, 29.6°, 131.6°) |

**Table S148.** Examples (i.e. identities of bases involved in stacking and corresponding PDB codes) of consecutive (black) and non-consecutive (grey) 6||6 β||β *trans* purine||pyrimidine stacks. The geometrical parameters ($\vec{d}_{ab}$, $\theta_{ab}$, $\tau_a$, $\tau_b$ and $\sigma_{ab}$) for each example stack are provided in parentheses respectively.

| | | β face (6) | |
|---|---|---|---|
| | | C | U |
| β face (6) | A | A1419(BA)\|\|C1420(BA); 4v9h (4.3 Å, 17.1°, 39.2°, 24.0°, 116.9°) | |
| | | A35(A)\|\|C61(A); 3gao (3.7 Å, 10.4°, 14.0°, 24.0°, 169.7°) | A23(B)\|\|U48(B); 5swd (3.5 Å, 14.9°, 29.2°, 15.3°, 158.5°) |
| | G | | |
| | | G1378(0)\|\|C2747(0); 3cpw (3.6 Å, 16.8°, 12.6°, 16.7°, 176.1°) | G2950(1)\|\|U2979(1); 4u4y (3.5 Å, 6.4°, 11.0°, 15.9°, 123.1°) |

**Table S149.** Examples (i.e. identities of bases involved in stacking and corresponding PDB codes) of consecutive (black) and non-consecutive (grey) 5||6 β||β *trans* purine||pyrimidine stacks. The geometrical parameters ($\vec{d}_{ab}$, $\theta_{ab}$, $\tau_a$, $\tau_b$ and $\sigma_{ab}$) for each example stack are provided in parentheses respectively.

| | | β face (6) | |
|---|---|---|---|
| | | C | U |
| β face (5) | A | C546(1H)\|\|A547(1H);4wzd (3.8 Å, 14.3°, 16.3°, 29.4°, 94.0°) | A1536(BA)\|\|U1535(BA);4v67 (4.1 Å, 20.3°, 24.1°, 4.6°, 96.3°) |
| | | A532(DA)\|\|C2021(DA); 4u24 (3.9 Å, 21.6°, 17.4°, 37.3°, 127.6°) | A215(2)\|\|U242(2); 4u6f (3.8 Å, 9.6°, 31.9°, 22.4°, 165.0°) |
| | G | G117(A)\|\|C116(A);4faq (4.0 Å, 9.8°, 27.5°, 33.2°, 103.5°) | G183(AA)\|\|U182(AA);4v9b (4.0 Å,12.8°, 29.1°, 29.8°, 98.6°) |
| | | G1093(RA)\|\|C1072(RA); 1vvj (4.1 Å, 4.8°, 39.9°, 35.1°, 110.1°) | G1127(CA)\|\|U1125(CA); 4u24 (4.3 Å, 11.0°, 32.6°, 26.2°, 112.2°) |

**Table S150.** Percent occurrence frequency of three most dominant topologies of pyrimidine||pyrimidine stacks.

| **Topology** | **Percent occurrence** |
|---|---|
| 6\|\|6 α\|\|β or β\|\|α cis | 93.5% |
| 6\|\|6 β\|\|β cis | 1.8% |
| 6\|\|6 β\|\|β trans | 1.6% |

**Table S151.** Examples (i.e. identities of bases involved in stacking and corresponding PDB codes) of consecutive (black) and non-consecutive (grey) 6||6 α||α *cis* pyrimidine||pyrimidine stacks. The geometrical parameters ($\vec{d}_{ab}$, $\theta_{ab}$, $\tau_a$, $\tau_b$ and $\sigma_{ab}$) for each example stack are provided in parentheses respectively.

| | | α face (6) | |
|---|---|---|---|
| | | C | U |
| α face (6) | C | C2666(BB)\|\|C2667(BB); 4v4q (3.6 Å, 14.3°, 12.9°, 6.5°, 54.1°) | C846(B)\|\|U(B); 6boh (4.1 Å, 16.1°, 24.0°, 39.3°, 7.8°) |
| | | C587(BA)\|\|C671(BA); 1vy6 (4.2 Å, 15.8°, 39.9°, 32.6°, 28.6°) | C1773(0)\|\|U2588(0);1y69 (4.2 Å, 9.9°, 37.2°, 38.7°, 17.2°) |
| | U | | U9(v)\|\|U10(v); 5uk4 (3.7 Å, 6.0°, 32.4°, 26.3°, 55.3°) |
| | | | U1944(BA)\|\|U1955(BA); 4u24 (3.8 Å, 15.8°, 32.7°, 22.2°, 22.0°) |

**Table S152.** Examples (i.e. identities of bases involved in stacking and corresponding PDB codes) of consecutive (black) and non-consecutive (grey) 6||6 α||α *trans* pyrimidine||pyrimidine stacks. The geometrical parameters ($\vec{d}_{ab}$, $\theta_{ab}$, $\tau_a$, $\tau_b$ and $\sigma_{ab}$) for each example stack are provided in parentheses respectively.

| | | α face (6) | |
|---|---|---|---|
| | | C | U |
| α face (6) | C | C83(1L)\|\|C84(1L);4wr6 (3.9 Å, 18.3°, 19.6°, 26.7°, 103.7°) | |
| | | C2006(0)\|\|C1982(0);1w2b (4.0 Å, 10.9°, 22.9°, 25.1°, 118.2°) | C1159(6)\|\|U1285(6);4u4u (4.0 Å, 10.1°, 28.4°, 38.5°, 170.2°) |
| | U | | |
| | | | U4(A)\|\|U1(A);4rgf (4.1 Å, 4.6°, 33.2°, 29.3°, 160.6°) |

**Table S153.** Examples (i.e. identities of bases involved in stacking and corresponding PDB codes) of consecutive (black) and non-consecutive (grey) 6||6 α||β *cis* pyrimidine||pyrimidine stacks. The geometrical parameters ($\vec{d}_{ab}$, $\theta_{ab}$, $\tau_a$, $\tau_b$ and $\sigma_{ab}$) for each example stack are provided in parentheses respectively.

| | | β face (6) | |
|---|---|---|---|
| | | C | U |
| α face (6) | C | C22(X)\|\|C23(X); 2gtt<br>(4.0 Å, 12.8°, 22.0°, 16.0°, 43.6°) | C5(D)\|\|U6(D);6fpx<br>(4.0 Å, 19.8°, 22.1°, 17.1°, 43.8°) |
| | | C45(x)\|\|C47(x);2gtt<br>(4.1 Å, 8.7°, 35.4°, 29.9°, 78.3°) | C13(X)\|\|U94(Z);6dn1<br>(3.8 Å, 14.9°, 33.8°, 19.7°, 74.1°) |
| | U | | U40(A)\|\|U41(A); 4kqy<br>(3.8 Å, 6.8°, 36.1°, 31.9°, 26.9°) |
| | | | U41(r)\|\|U43(r); 3pu4<br>(3.9 Å, 12.0°, 29.0°, 24.7°,63.7° ) |

**Table S154.** Examples (i.e. identities of bases involved in stacking and corresponding PDB codes) of consecutive (black) and non-consecutive (grey) 6||6 α||β *trans* pyrimidine||pyrimidine acks. The geometrical parameters ($\vec{d}_{ab}$, $\theta_{ab}$, $\tau_a$, $\tau_b$ and $\sigma_{ab}$) for each example stack are provided in parentheses respectively.

| | | β face (6) | |
|---|---|---|---|
| | | C | U |
| α face (6) | C | C2163(1H)\|\|C2164(1H); 5elh<br>(4.0 Å, 11.1°, 33.5°, 39.1°, 99.0°) | C-4(A)\|\|U-5(A); 4g6p<br>(4.4 Å, 7.6°, 31.0°, 37.4°, 92.2°) |
| | | C9(R)\|\|C11(R); 2iz8<br>(4.0 Å, 12.8°, 11.5°, 18.3°, 101.6°) | C562(A)\|\|884(A); 5wns<br>(4.0 Å, 8.0°, 24.2°, 16.6°, 128.4°) |
| | U | | U54(AD)\|\|U55(AD); 4v9b<br>(3.8 Å, 14.6°, 26.6°, 32.0°, 90.2°) |
| | | | U562(CA)\|\|U884(CA); 4v7v<br>(3.8 Å, 13.5°, 17.7°, 29.6°, 126.6°) |

**Table 155.** Examples (i.e. identities of bases involved in stacking and corresponding PDB codes) of consecutive (black) and non-consecutive (grey) 6||6 β||α *cis* pyrimidine||pyrimidine stacks. The geometrical parameters ($\vec{d}_{ab}$, $\theta_{ab}$, $\tau_a$, $\tau_b$ and $\sigma_{ab}$) for each example stack are provided in parentheses respectively.

| | | α face (6) | |
|---|---|---|---|
| | | C | U |
| β face (6) | C | | C512(D)‖U513(D);1g59<br>(4.0 Å, 11.0°, 37.6°, 35.2°, 30.3°) |
| | | | C48(D)‖U59(D);2zm5<br>(3.9 Å, 19.4°, 20.5°, 31.1°, 11.8°) |
| | U | | |

**Table S156.** Examples (i.e. identities of bases involved in stacking and corresponding PDB codes) of consecutive (black) and non-consecutive (grey) 6||6 β||α *trans* pyrimidine||pyrimidine stacks. The geometrical parameters ($\vec{d}_{ab}$, $\theta_{ab}$, $\tau_a$, $\tau_b$ and $\sigma_{ab}$) for each example stack are provided in parentheses respectively.

| | | α face (6) | |
|---|---|---|---|
| | | C | U |
| β face (6) | C | | C1(X)‖U2(X);4khp<br>(4.2 Å, 6.6°, 34.5°, 33.1°, 108.8°) |
| | | | C208(A)‖U210(A);2hhh<br>(3.9 Å, 9.9°, 34.6°, 31.5°, 130.0°) |
| | U | | |

**Table S157.** Examples (i.e. identities of bases involved in stacking and corresponding PDB codes) of consecutive (black) and non-consecutive (grey) 6||6 β||β *cis* pyrimidine||pyrimidine stacks. The geometrical parameters ($\vec{d}_{ab}$, $\theta_{ab}$, $\tau_a$, $\tau_b$ and $\sigma_{ab}$) for each example stack are provided in parentheses respectively.

| | | β face (6) | |
|---|---|---|---|
| | | C | U |
| β face (6) | C | C5(D)\|\|C5(D); 6fpk (4.2 Å, 14.6°, 28.5°, 30.6°, 25.7°) | C1149(AA)\|\|U1150(AA); 4w5s (4.3 Å, 14.8°, 24.8°, 39.5°, 30.4°) |
| | | C1728(DA)\|\|C1730(DA);4u1v (4.3 Å, 7.8°, 27.4°, 33.1°, 27.0°) | C1054(A)\|\|U1096(A); 6cao (4.3 Å, 15.4°, 13.4°, 14.1°, 40.8°) |
| | U | | U2888(A)\|\|U2889(A); 1k8a (3.8 Å, 18.9°, 18.3°, 12.9°, 66.5°) |
| | | | U33(E)\|\|U35(E);4j1g (3.5 Å,18.2°, 38.4°, 20.3°, 67.4°) |

**Table S158.** Examples (i.e. identities of bases involved in stacking and corresponding PDB codes) of consecutive (black) and non-consecutive (grey) 6||6 β||β *trans* pyrimidine||pyrimidine stacks. The geometrical parameters ($\vec{d}_{ab}$, $\theta_{ab}$, $\tau_a$, $\tau_b$ and $\sigma_{ab}$) for each example stack are provided in parentheses respectively.

| | | β face (6) | |
|---|---|---|---|
| | | C | U |
| β face (6) | C | | C1018(0)\|\|U1019(0);1xbp (4.3 Å, 15.6°, 19.4°, 19.2°, 100.0°) |
| | | C130(0)\|\|C141(0);3g4s (4.1 Å, 16.4°, 13.1°, 17.8°, 127.5°) | C372(CA)\|\|U387(CA); 4u24 (4.1 Å, 7.5°, 24.4°,22.0°, 168.3°) |
| | U | | U269(14)\|\|U270(14); 4wra (3.9 Å, 15.7°, 35.4°, 23.4°, 158.8°) |

.

**Table S159**. Comparison of the method of detection of all topologies of A||G stacks by our methods with other available methods.

| Stack | our method | FR3D | 3DNA | MC-Annotate |
|---|---|---|---|---|
| A900(A)\|\|G901(A); 6hhq | 5\|\|56, 6\|\|56 α\|\|α *cis* consecutive | s33 | Part of n = 4 stack | adjacent_5p inward |
| A1287(DA)\|\|G1288(DA); 4u24 | 5\|\|6, 6\|\|56 α\|\|α *cis* consecutive | s33 | Part of n = 5 stack | adjacent_5p inward |
| G973(AA)\|\|A974(AA); 6i7v | 5\|\|5, 6\|\|56 α\|\|α *cis* consecutive | s33 | Part of n = 2 stack | adjacent_5p inward |
| G1806(X)\|\|A1807(X); 4io9 | 5\|\|56, 6\|\|6 α\|\|α *cis* consecutive | s33 | Part of n = 5 stack | adjacent_5p inward |
| A2841(0)\|\|G2842(0);3i55 | 5\|\|56, 6\|\|5 α\|\|α *cis* consecutive | s33 | Part of n = 8 stack | adjacent_5p inward pairing |
| A7(C)\|\|G8(C); 4z0c | 5\|\|6, 6\|\|6 α\|\|α *cis* consecutive | s33 | Part of n = 3 stack | adjacent_5p inward |
| A1287(DA)\|\|G1288(DA); 4v7t | 6\|\|56 α\|\|α *cis* consecutive | s33 | Part of n = 5 stack | adjacent_5p inward |
| G1814(GA)\|\|A1815(GA);4v9o | 5\|\|5, 6\|\|6 α\|\|α *cis* consecutive | s33 | Part of n = 3 stack | adjacent_5p inward |
| A427(x)\|\|G428(x);4wfa | 5\|\|6, 6\|\|5 α\|\|α cis consecutive | s33 | Part of n = 5 stack | adjacent_5p inward |
| A1260(5)\|\|G1261(5); 4u55 | 5\|\|56 α\|\|α *cis* consecutive | s33 | Part of n = 4 stack | adjacent_5p inward |
| A2841(0)\|\|G2842(0); 1vq5 | 5\|\|5, 6\|\|5 α\|\|α *cis* consecutive | s33 | Part of n = 5 stack | adjacent_5p inward pairing |
| G1348(DA)\|\|A1349(DA);4v5e | 6\|\|6 α\|\|α *cis* consecutive | s33 | Part of n = 2 stack | adjacent_5p inward |
| A2841(0)\|\|G2842(0); 3ccj | 5\|\|6 α\|\|α *cis* consecutive | s33 | Part of n = 5 stack | adjacent_5p inward pairing |
| A1287(CA)\|\|G1288(CA); 4v9p | 6\|\|5 α\|\|α *cis* consecutive | s33 | Part of n = 5 stack | adjacent_5p inward |
| A2841(A)\|\|G2842(A); 1nji | 5\|\|5 α\|\|α *cis* consecutive | s33 | Part of n = 8 stack | adjacent_5p inward pairing |
| A1070(0)\|\|G1071(0);1yj9 | 5\|\|56, 6\|\|56 α\|\|α *trans* consecutive | s33 | Part of n = 4 stack | adjacent_5p inward |
| A21(1x)\|\|G46(1x); 5hcq | 5\|\|6, 6\|\|56 α\|\|α *trans* non-consecutive | s33 | Part of n = 7 stack | non-adjacent-inward |
| G75(3)\|\|A76(3); 5dgf | 5\|\|5, 6\|\|56 α\|\|α *trans* consecutive | s33 | Part of n = 5 stack | adjacent_5p inward pairing |
| G2308(BA)\|\|A2309(BA); 4v7x | 5\|\|56, 6\|\|6 α\|\|α *trans* consecutive | s33 | Part of n = 2 stack | adjacent_5p inward |
| A21(IB)\|\|G46(IB); 5j4d | 5\|\|56, 6\|\|5 α\|\|α *trans* non-consecutive | s33 | Part of n = 6 stack | non-adjacent-inward pairing |
| A66(BB)\|\|G107(BB); 4u24 | 5\|\|6, 6\|\|6 α\|\|α *trans* non-consecutive | s33 | Part of n = 5 stack | non-adjacent inward |
| A34(DB)\|\|G44(DB); 4u27 | 6\|\|56 α\|\|α *trans* non-consecutive | s33 | Part of n = 2 stack | non-adjacent inward pairing |
| G1019(14)\|\|A1020(14);5ndk | 5\|\|5, 6\|\|6 α\|\|α *trans* consecutive | s33 | Part of n = 4 stack | adjacent_5p inward |
| G60(A)\|\| A61 (A); 4nya | 5\|\|56 α\|\|α *trans* consecutive | s33 | Part of n = 4 stack | adjacent_5p inward |
| G90(X)\|\|A91 (X); 5dm6 | 5\|\|5, 6\|\|5 α\|\|α *trans* consecutive | s33 | Part of n = 2 stack | adjacent_5p inward pairing |
| G720 (2A)\|\| A850 (2A);4z8c | 6\|\|6 α\|\|α *trans* non-consecutive | s33 | Part of n = 6 stack | non-adjacent inward |
| A1028(DA)\|\|G1125(DA); 4u27 | 5\|\|6 α\|\|α *trans* non-consecutive | s33 | Part of n = 3 stack | non-adjacent inward |
| A22(CD)\|\|G47(CD); 4v8b | 6\|\|5 α\|\|α *trans* non-consecutive | s33 | Part of n = 6 stack | non-adjacent inward |

| | | | | |
|---|---|---|---|---|
| A889 (CA)‖G888 (CA); 4v67 | 5‖5 α‖α *trans* consecutive | s33 | Part of n = 4 stack | adjacent_5p inward |
| A5(A)‖G6(A);4lvv | 5‖56, 6‖56 α‖β *cis* consecutive | s35 | Part of n = 5 stack | adjacent_5p upward |
| A495(0)‖ G496(0); 1yit | 5‖6, 6‖56 α‖β *cis* consecutive | s35 | Part of n = 9 stack | adjacent_5p upward |
| A503(BA)‖G506(BA); 4u24 | 5‖5, 6‖56 α‖β *cis* non-consecutive | s35 | Part of n = 11 stack | non-adjacent upward |
| A1632(DA)‖G1633(DA); 4u24 | 5‖56, 6‖6 α‖β *cis* consecutive | s35 | Part of n = 4 stack | adjacent_5p upward |
| A583(1G)‖G584(1G);4wt1 | 5‖56, 6‖5 α‖β *cis* consecutive | s35 | Part of n = 5 stack | adjacent_5p upward |
| A1433(5)‖G1434(5); 4u51 | 5‖6, 6‖6 α‖β *cis* consecutive | s35 | Part of n = 2 stack | adjacent_5p upward |
| A9(A)‖G10(A); 3vrs | 6‖56 α‖β *cis* consecutive | s35 | - | adjacent_5p upward |
| A1042(CA)‖A1043(CA); 4u24 | 5‖5, 6‖6 α‖β cis consecutive | s35 | Part of n = 8 stack | adjacent_5p upward |
| A14(CW)‖G15(CW); 4v8n | 5‖56 α‖β *cis* consecutive | s35 | Part of n = 9 stack | adjacent_5p upward |
| A20(C)‖G21(C); 2d2l | 5‖5, 6‖5 α‖β *cis* consecutive | s35 | Part of n = 9 stack | adjacent_5p upward |
| A20(AX)‖G21(AX);4v5a | 6‖6 α‖β *cis* consecutive | s35 | Part of n = 3 stack | adjacent_5p upward |
| A1433(5)‖G1434(5); 4u55 | 5‖6 α‖β *cis* consecutive | s35 | Part of n = 2 stack | adjacent_5p upward |
| A18(C)‖G19(C); 2d2l | 6‖5 α‖β *cis* consecutive | s35 | - | adjacent_5p upward |
| A8(A)‖G9(A);4lvw | 5‖5 α‖β *cis* consecutive | s35 | Part of n = 4 stack | adjacent_5p upward |
| A2725(BA)‖G2727(BA);4v8a | 5‖56, 6‖56 α‖β *trans* non-consecutive | s35 | Part of n = 2 stack | non-adjacent upward pairing |
| A1069(DA)‖G1074(DA); 4u24 | 5‖6, 6‖56 α‖β *trans* non-consecutive | s35 | Part of n = 5 stack | non-adjacent upward |
| A119(CA)‖G240(CA); 4v7s | 5‖5, 6‖56 α‖ β *trans* non-consecutive | s35 | Part of n = 8 stack | non-adjacent upward |
| A2764(CA)‖G2766(CA); 4wqf | 5‖56, 6‖6 α‖β trans non- consecutive | s35 | Part of n = 6 stack | non-adjacent upward |
| A2369(0)‖G2371(0);3cc7 | 5‖56, 6‖5 α‖β *trans* non- consecutive | s35 | Part of n = 2 stack | non-adjacent upward pairing |
| A45(0)‖G147(0);1ffk | 5‖6, 6‖6 α‖β *trans* non- consecutive | s35 | Part of n = 2 stack | non-adjacent upward pairing |
| A1069(BA)‖G1074(BA); 4u24 | 6‖56 α‖β *trans* non- consecutive | s35 | Part of n = 9 stack | non-adjacent upward |
| A451(CA)‖G481(CA);4v8f | 5‖6, 6‖5 α‖β *trans* non- consecutive | s35 | Part of n = 8 stack | - |
| A1067(AA)‖G1068(AA); 4v87 | 5‖56 α‖β *trans* consecutive | s35 | Part of n = 5 stack | adjacent_5p upward |
| A35(A)‖G62(A);3g4m | 6‖6 α‖β *trans* non- consecutive | s35 | Part of n = 3 stack | non-adjacent upward |
| A451(AA)‖G481(AA); 4u27 | 5‖6 α‖β *trans* non-consecutive | s35 | Part of n = 8 stack | non-adjacent upward pairing |
| A221(BA)‖G266(BA); 4u24 | 6‖5 α‖β *trans* non-consecutive | s35 | Part of n = 5 stack | non-adjacent upward |
| A460(6)‖G461(6);5fci | 5‖5 α‖β *trans* consecutive | s35 | Part of n = 3 stack | adjacent_5p upward |
| G1525(0)‖A1526(0);3ccj | 5‖56, 6‖56 β‖α *cis* consecutive | s53 | Part of n = 5 stack | adjacent_5p downward pairing |

| | | | | |
|---|---|---|---|---|
| A109(XA)‖G326(XA);4www | 5‖6, 6‖56 β‖α *cis* non-consecutive | s53 | Part of n = 10 stack | non-adjacent downward pairing |
| G1878(5)‖1879(5); 5fcj | 5‖5, 6‖56 β‖α *cis* consecutive | s53 | Part of n = 6 stack | adjacent_5p downward pairing |
| A1(D)‖G2(D); 6dcl | 5‖56, 6‖6 β‖α *cis* consecutive | s53 | Part of n = 2 stack | adjacent_5p downward |
| A2513(A)‖G2564(A); 1ffz | 5‖56, 6‖5 β‖α *cis* non-consecutive | s53 | Part of n = 3 stack | non-adjacent downward pairing |
| A33(A)‖G34(A); 3d2g | 5‖6, 6‖6 β‖α *cis* consecutive | s53 | Part of n = 3 stack | adjacent_5p downward |
| A496(BA)‖G497(BA); 4ybb | 6‖56 β‖α *cis* consecutive | s53 | Part of n = 6 stack | adjacent_5p downward |
| G2334(14)‖A2335(14); 5ndk | 5‖5, 6‖6 β‖α *cis* consecutive | s53 | Part of n = 5 stack | adjacent_5p downward |
| G2319(1H)‖A2320(1H); 5el4 | 5‖56 β‖α *cis* consecutive | s53 | Part of n = 5 stack | adjacent_5p downward |
| A2478(RA)‖G2529(RA); 4lt8 | 5‖5, 6‖5 β‖α *cis* non-consecutive | s53 | Part of n = 3 stack | non-adjacent downward |
| A159(A)‖G160(A);4fax | 6‖6 β‖α *cis* consecutive | s53 | Part of n = 6 stack | adjacent_5p downward |
| A163(B)‖G164(B); 488d | 5‖6 β‖α *cis* consecutive | s53 | - | adjacent_3p downward |
| A496(AA)‖G497(AA); 4woi | 6‖5 β‖α *cis* consecutive | s53 | Part of n = 6 stack | - |
| G2319(14)‖A2320(14);5ibb | 5‖5 β‖α *cis* consecutive | s53 | Part of n = 5 stack | adjacent_5p downward |
| A2404(BA)‖G2441(BA);4w2g | 5‖56, 6‖56 β‖α *trans* non-consecutive | s53 | Part of n = 5 stack | non-adjacent downward |
| G778(2)‖A780(2); 5dgv | 5‖6, 6‖56 β‖α *trans* non-consecutive | s53 | Part of n = 3 stack | non-adjacent downward |
| A2847(1)‖G2898(1); 4u55 | 5‖5, 6‖56 β‖α *trans* non-consecutive | s53 | Part of n = 3 stack | non-adjacent downward |
| A2457(0)‖G2508(0); 2aar | 5‖56, 6‖6 β‖α *trans* non-consecutive | s53 | Part of n = 3 stack | non-adjacent downward |
| A2457(A)‖G2508(A);1jzy | 5‖56, 6‖5 β‖α *trans* non- consecutive | s53 | Part of n = 8 stack | non-adjacent downward |
| A812(2)‖G858(2); 4u55 | 5‖6, 6‖6 β‖α *trans* non-consecutive | s53 | Part of n = 12 stack | non-adjacent downward |
| G778(A)‖A780(A); 6hhq | 6‖56 β‖α *trans* non-consecutive | s53 | Part of n = 3 stack | non-adjacent downward |
| G2157(1)\|A2178(1);4u3n | 5‖6, 6‖5 β‖α *trans* non-consecutive | s53 | Part of n = 3 stack | non-adjacent downward pairing |
| A53(A)‖G83(A); 2hop | 5‖56 β‖α *trans* non-consecutive | s53 | Part of n = 4 stack | non-adjacent downward |
| A2478(BA)‖G2529(BA); 4v9l | 5‖5, 6‖5 β‖α *trans* non-consecutive | s53 | Part of n = 3 stack | non-adjacent downward |
| G721(AA)‖A733(AA);4v5e | 6‖6 β‖α *trans* non-consecutive | s53 | - | non-adjacent downward |
| A803(1a)‖G1507(1a); 5hcq | 5‖6 β‖α *trans* non-consecutive | s53 | Part of n = 3 stack | non-adjacent downward |
| A2810(X)‖G2854(X); 3pip | 6‖5 β‖α *trans* non-consecutive | s53 | Part of n = 5 stack | non-adjacent downward pairing |
| A577(X)‖G2048(X);4wf9 | 5‖5 β‖α *trans* non-consecutive | s53 | - | non-adjacent downward |
| A493(CA)‖G494(CA); 4v6c | 5‖56, 6‖56 β‖β *cis* consecutive | s55 | Part of n = 6 stack | adjacent_5p outward |

| | | | | |
|---|---|---|---|---|
| G1878 (1)\|\| A1879 (1);5tbw | 5\|\|6, 6\|\|56 β\|\|β *cis* consecutive | s55 | Part of n = 6 stack | adjacent_5p outward |
| A646(DA)\|\|G647(DA); 4v9k | 5\|\|5, 6\|\|56 β\|\|β *cis* consecutive | s55 | Part of n = 4 stack | adjacent_5p outward |
| A646(1H)\|\|G647(1H); 4wt1 | 5\|\|56, 6\|\|6 β\|\|β *cis* consecutive | s55 | Part of n = 4 stack | adjacent_5p outward |
| G1604(5)\|\|A1605(5); 4u4y | 5\|\|56, 6\|\|5 β\|\|β *cis* consecutive | s55 | Part of n = 4 stack | adjacent_5p outward |
| A493(CA)\|\|G494(CA); 4v7u | 5\|\|6, 6\|\|6 β\|\|β *cis* consecutive | s55 | Part of n = 4 stack | adjacent_5p outward |
| A878(14)\|\|G879(14); 4wq1 | 6\|\|56 β\|\|β *cis* consecutive | s55 | Part of n = 5 stack | adjacent_5p outward |
| G1878(1)\|\| A1879(1);5dgf | 5\|\|6, 6\|\|5 β\|\|β *cis* consecutive | s55 | Part of n = 6 stack | adjacent_5p outward |
| A56(R)\|\|G57(R); 6b14 | 5\|\|56 β\|\|β *cis* consecutive | s55 | Part of n = 3 stack | adjacent_5p outward |
| A1050(BA)\|\|G1051(BA); 4v8b | 5\|\|5, 6\|\|5 β\|\|β *cis* consecutive | s55 | Part of n = 6 stack | adjacent_5p outward |
| A17(C)\|\| G18(C);5dea | 6\|\|6 β\|\|β *cis* consecutive | s55 | Part of n = 4 stack | - |
| G921(0)\|\|A922(0); 3ccu | 5\|\|6 β\|\|β *cis* consecutive | s55 | Part of n = 2 stack | adjacent_5p outward |
| A2598(H)\|\|G2599(H); 3dh3 | 6\|\|5 β\|\|β *cis* consecutive | s55 | Part of n = 2 stack | adjacent_5p outward |
| G1503(2)\|\|A1504(2);5dat | 5\|\|5 β\|\|β *cis* consecutive | s55 | Part of n = 3 stack | adjacent_5p outward |
| A27(CB)\|\|G28(CB);4v8f | 5\|\|56,6\|\|56 β\|\|β *trans* consecutive | s55 | Part of n = 3 stack | adjacent_5p outward |
| A14(CX)\|\| G34(CW); 4v5d | 5\|\|6, 6\|\|56 β\|\|β *trans* non-consecutive | s55 | Part of n = 9 stack | non-adjacent outward |
| A2469(1H)\|\|G2470(1H); 5el5 | 5\|\|5, 6\|\|56 β\|\|β *trans* consecutive | s55 | Part of n = 5 stack | adjacent_5p outward |
| A26(1K)\|\|G27(1K); 5el6 | 5\|\|56, 6\|\|6 β\|\|β *trans* consecutive | s55 | Part of n = 7 stack | adjacent_5p outward |
| A2448(0)\|\|G2461(X);1xbp | 5\|\|56, 6\|\|5 β\|\|β *trans* non-consecutive | s55 | Part of n = 7 stack | non-adjacent outward |
| A1239(CA)\|\|G1241(CA); 4u24 | 5\|\|6, 6\|\|6 β\|\|β *trans* non-consecutive | s55 | Part of n = 2 stack | non-adjacent outward |
| A2469(YA)\|\|G2470(YA);6buw | 6\|\|56 β\|\|β *trans* consecutive | s55 | Part of n = 5 stack | adjacent_5p outward |
| G570(a)\|\|A873(a);4jv5 | 5\|\|5, 6\|\|6 β\|\|β *trans* non-consecutive | s55 | Part of n = 2 stack | non-adjacent outward |
| A1502(CA)\|\|G1504(CA); 4u24 | 5\|\|56 β\|\|β *trans* non-consecutive | s55 | - | non-adjacent outward |
| A933(DA)\|\|G934(DA); 4v8b | 5\|\|5, 6\|\|5 β\|\|β *trans* consecutive | s55 | Part of n = 4 stack | adjacent_5p outward |
| A196(BA)\|\|G805(BA); 4u24 | 6\|\|6 β\|\|β *trans* non-consecutive | s55 | Part of n = 2 stack | non-adjacent outward |
| A1875(A)\|\|G1877(A); 1q81 | 5\|\|6 β\|\|β *trans* non-consecutive | s55 | Part of n = 3 stack | non-adjacent outward |
| A2469(DA)\|\|G2470(DA);4v8b | 6\|\|5 β\|\|β *trans* consecutive | s55 | Part of n = 5 stack | adjacent_5p outward |
| A26(1L)\|\|G27(1L);4wro | 5\|\|5 β\|\|β *trans* consecutive | s55 | Part of n = 2 stack | adjacent_5p outward |

**Table S160.** Comparison of stacking interaction identified in the Loop E of bacterial 5S rRNA (PDB code: 364d) using our method and FR3D.

| Stack | FR3D | our method |
|---|---|---|
| 71‖72 | S35 | 6‖5 β‖β *cis* |
| 73‖74 | S35 | 5‖6, 6‖6 α‖α *cis* |
| 74‖75 | S35 | 6‖5 β‖β *cis* |
| 75‖76 | S35 | 5‖5, 6‖56 α‖β *cis* |
| 76‖77 | S35 | 5‖6, 6‖6 α‖α *cis* |
| 78‖79 | S35 | 6‖5 α‖α *cis* |
| 105‖104 | S53 | 5‖5 β‖α *cis* |
| 104‖103 | S53 | 5‖6 β‖β *cis* |
| 103‖102 | S53 | 6‖5, 6‖6 α‖α *cis* |
| 102‖101 | S53 | 5‖6 β‖α *cis* |
| 101‖100 | S53 | 56‖6 β‖α *cis* |
| 100‖99 | S53 | 5‖6 β‖α *cis* |
| 98‖97 | S53 | 5‖6 β‖β *cis* |
| 104‖73 | S55 | 6‖5 β‖β *trans* |
| 102‖75 | S55 | not detected |
| 99‖78 | S55 | 5‖56, 6‖56 β‖β *trans* |
